# Genome sequencing and molecular networking analysis of the wild fungus *Anthostomella pinea* reveal its ability to produce a diverse range of secondary metabolites

**DOI:** 10.1101/2023.10.20.563261

**Authors:** R. Iacovelli, T. He, J. L. Allen, T. Hackl, K. Haslinger

## Abstract

**Background:** Filamentous fungi are prolific producers of bioactive molecules and enzymes with important applications in industry. Yet, the vast majority of fungal species remain undiscovered or uncharacterized. Here we focus our attention to a wild fungal isolate that we identified as *Anthostomella pinea*. The fungus belongs to a complex polyphyletic genus in the family of *Xylariaceae*, which is known to comprise endophytic and pathogenic fungi that produce a plethora of interesting secondary metabolites. Despite that, *Anthostomella* is largely understudied and only two species have been fully sequenced and characterized at a genomic level.

**Results:** In this work, we used long-read sequencing to obtain the complete 53.7 Mb genome sequence including the full mitochondrial DNA. We performed extensive structural and functional annotation of coding sequences, including genes encoding enzymes with potential applications in biotechnology. Among others, we found that the genome of *A. pinea* encodes 91 biosynthetic gene clusters, more than 600 CAZymes, and 164 P450s. Furthermore, untargeted metabolomics and molecular networking analysis of the cultivation extracts revealed a rich secondary metabolism, and in particular an abundance of sesquiterpenoids and sesquiterpene lactones. We also identified the polyketide antibiotic xanthoepocin, to which we attribute the anti–Gram-positive effect of the extracts that we observed in antibacterial plate assays.

**Conclusions:** Taken together, our results provide a first glimpse into the potential of *Anthstomella pinea* to provide new bioactive molecules and biocatalysts and will facilitate future research into these valuable metabolites.

## Background

Fungi are ubiquitous organisms and major components of all ecosystems on Earth, although they more commonly colonize soil and plant tissues. To thrive and adapt to environmental stressors (e.g., extreme temperatures, low water and nutrient availabilities, predators), they can adopt different lifestyles ranging from parasites and opportunistic pathogens to specialized biomass decomposers and mutualistic symbionts [1–3]. The latter include endophytic fungi, that is, fungi that can live inside plant tissues throughout their life cycle and establish a beneficial or neutral relationship with their host, thereby not causing any detrimental effect or disease [4,5]. In exchange for nutrients and a habitat, endophytic symbionts offer several benefits to their host: for example, they can promote growth or protect from biotic (predators, pathogens) or abiotic stressors (UV radiation) [1,6,7]. Generally, they exert these functions through the production of small bioactive molecules called secondary metabolites (SMs), which include polyketides, alkaloids, peptides, terpenoids, and phenolic compounds [8,9]. Often SMs possess biological activities that translate into pharmaceutical applications, such as antimicrobial, antioxidant and anticancer activities [8]. Among the most encountered endophytic fungi are members of the family *Xylariaceae*, which also comprises saprotrophic, pathogenic, and endolichenic (that live inside lichen tissues) fungi [10–12]. Endophytic fungi, and in particular *Xylariaceae*, have been extensively studied in the last decades to discover and exploit new pharmaceutically relevant compounds [4,7,10,13,14]. Despite this, the vast majority of the *Xylariaceae* species studied so far belong to the eponymous genus *Xylaria* and a few others, while many are yet to be characterized [10].

In this work, we isolated a filamentous fungus from sections of dry lichen thalli collected in Cheney, Washington, USA. We identified the fungus as *Anthostomella pinea*, previously described as an epiphytic fungus of pine trees [15]. *Anthostomella* is a large and complex polyphyletic genus that comprises more than 400 species [16] within the *Xylariaceae* family, though little to nothing is known about their genetic makeup and metabolome. To the best of our knowledge, only a few publications report the discovery and characterization of metabolites from *Anthostomella* fungi [17–19], and complete genomic data is currently available for only two species [20,21]. Therefore, we set out to perform whole genome sequencing and untargeted metabolomics analyses to shed light on the ability of the fungus to produce biotechnologically relevant secondary metabolites and enzymes that can be used as biocatalysts in industrial applications. To do so, we employed state-of-the-art long read sequencing technologies that allowed us to obtain a high-quality annotated genome of the fungus. We then used high-resolution tandem mass-spectrometry coupled to molecular networking analysis to analyze the intra- and extra-cellular metabolome. With these approaches, we revealed that *A. pinea* is a prolific producer of sesquiterpenoids and sesquiterpene lactones, and that its genome encodes an abundance of biotechnologically relevant enzymes such as carbohydrate-active enzymes (CAZymes), cytochrome P450s (CYP), and unspecific peroxygenases (UPOs). We also detected antimicrobial activity of the extracts, which we putatively attribute to the anti–Gram-positive compound xathoepocin, produced in small amounts by the fungus. Lastly, based on the genomics data, we hypothesize the biosynthetic route of this compound, which could prove useful in future efforts to reconstitute the pathway in a heterologous host for large scale production and engineering of structural variants.

## Results and Discussion

### Isolation of *Anthostomella pinea* F5 from sections of lichen thallus and morphological characterization

Lichenized fungi, those that form obligate mutualistic symbioses with a photosynthetic partner, are known for their rich SM profiles [22] and equally rich endophytic fungal communities [23]. The wolf lichens, *Letharia*, are a genus of lichens that have received substantial recent research attention due to their unique fungal associates [24,25] and genomic architecture [26]. All wolf lichens produce certain SMs in great abundance, as is evidenced by their bright yellow-green color. Thus, we chose to investigate the biosynthetic potential of one of the most common and widespread wolf lichens in western North America, *Letharia lupina*, and its associates. To obtain axenic fungal cultures, we first collected fresh specimen from a *Pinus ponderosa* growing by the Cheney Wetlands Trail in Cheney, Washington, USA (47.483392, −117.553087), and shipped them at room temperature to the research facilities in Groningen, The Netherlands. Next, we incubated surface-sterilized sections of the lichen thallus in sterile water, based on a modified version of the Yamamoto method [27]. After 4 weeks, filamentous growth was observed on the lichen sections and ten isolates (F1-F10) were excised and transferred to MYA plates: three showed robust growth of white mycelium (F5, F6, and F7), while the other seven showed much poorer growth and dark-green mycelium. From the initial morphological observations and preliminary barcoding with the ITS region, we confirmed that isolates F1-4 and F8-10 were most likely the same species but because of poor ITS alignments and slow growth they were not further investigated (data not shown). Similarly, we determined F5, F6, and F7 to be the same species, and we therefore continued only with isolate F5.

For further morphological characterization we inoculated F5 on different nutrient media, including MEA, PDA, YES, CYA, and DG18, which features a low water activity and is used to grow and characterize xerophilic fungi [28]. Isolate F5 showed robust growth in all the tested media but CYA, where it stopped growing approximately after 14 days (Fig. 1A). In all cases, it produced white vegetative mycelium which appeared less compact in DG18 and YES, likely attributable to stressful conditions (i.e., low water activity, and poorer availability of nutrients, respectively). Furthermore, F5 secreted yellow extrolites into the medium, which is particularly visible on the reverse side of the plates. On MEA, PDA, and DG18, the isolate showed a strong orange pigmentation at the inoculum site and around the ageing zone, visible both on the observe and reverse sides, but much more prominent on the latter (Fig. 1A). Lastly, extensive guttation was observed on the surface of the colonies on PDA. These observations are a strong indication that the isolate is able to produce and secrete secondary metabolites [29,30] in laboratory conditions.

**Figure 1.**
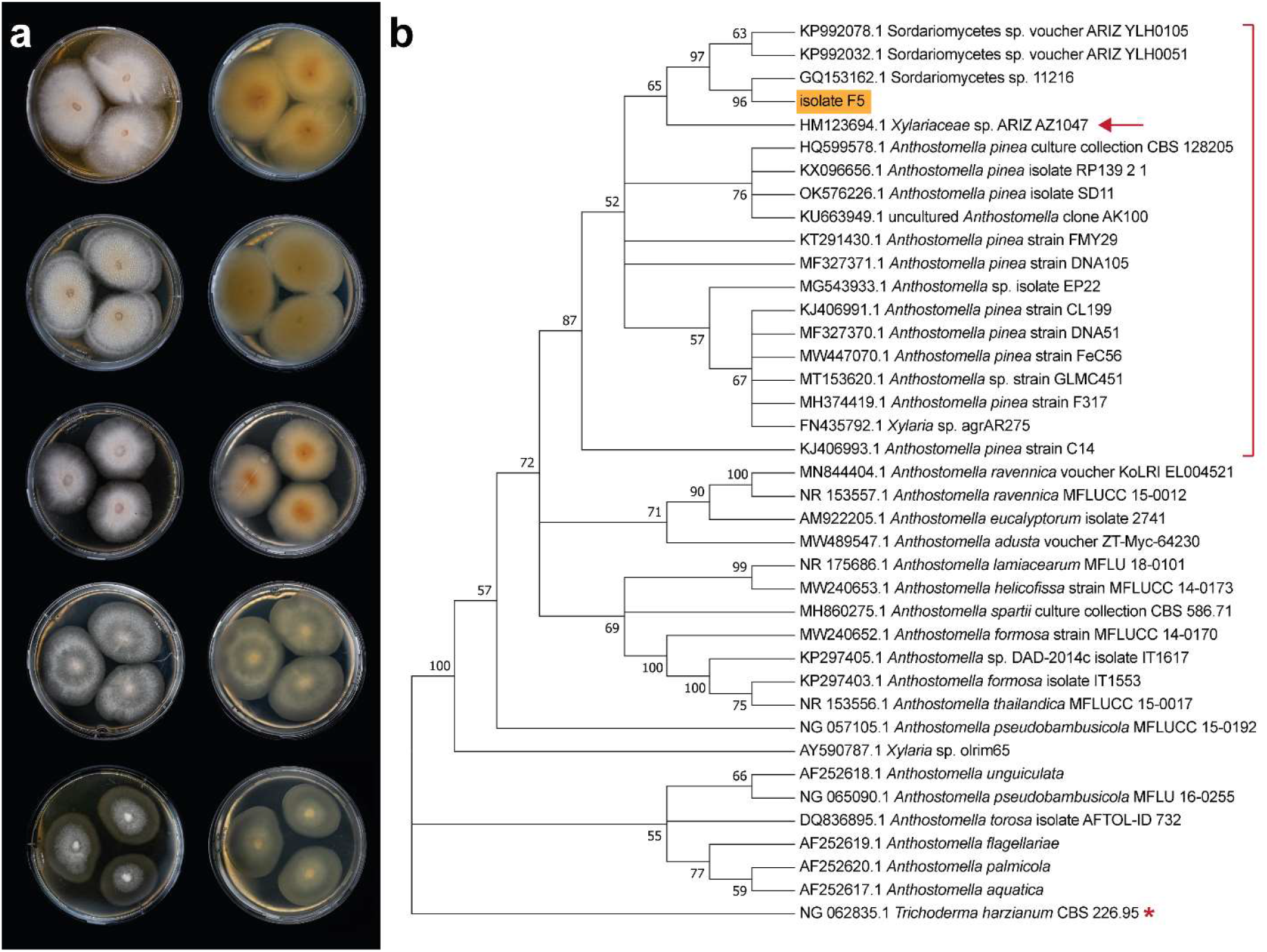
**(a)** Morphological characterization of F5 isolate via cultivation on solid medium. From top to bottom: MEA, PDA, DG18, YES, and CYA; obverse (left) and reverse (right). The cultures were incubated for 28 days at 20 °C. **(b)** Maximum-likelihood phylogenetic tree based on ITS sequences retrieved from the NCBI database. The hypothesized cluster of A. pinea strains is highlighted by the red bracket. Isolate F5 is highlighted in orange, next to isolate AZ1047 which showed almost identical morphology (arrow). T. harzanium (starred) was used as an outgroup to root the tree.

Next, we extracted genomic DNA from the fungal colonies to amplify the Internally Transcribed Spacer (ITS), a universal barcode region for phylogenetic analysis and taxonomic assignment [31]. We analyzed the manually curated 661 bp-long sequence by blastn with standard settings to search for hits within the kingdom of Fungi. The best 25 hits (ID > 95%) are shown in Table S1. The highest ranked hit for an accepted species was *Anthostomella pinea* strain CBS128205, an ex-type strain isolated from pine needles (Fungal Planet 53, 23 December 2010). Thus, we compared the 25 best hits and all the ITS sequences of other *Anthostomella* species available on NCBI (18 in total) by constructing a maximum-likelihood phylogenetic tree using *Trichoderma harzanium*, a phylogenetically unrelated Sordariomycete, as an outgroup (Fig. 1B). We observed that F5 clustered mainly with different strains of *A. pinea* and with several unidentified Sordariomycete species. Interestingly, F5 showed very similar morphology to isolate AZ1047 which was itself identified as *A. pinea* (Fig S1) [32,33] and for which barcode sequences of the *rbp2* and *act* genes were also available. Sequence alignments with the respective barcodes from F5 (obtained from whole genome sequencing) showed near-perfect matches (Fig S2 and S3), confirming F5 to be an isolate of *A. pinea* closely related to isolate AZ1047. Based on the phylogenetic tree, we hypothesize that the three not further classified Sordariomycete isolates that cluster together with F5 are themselves ascribable to *Anthostomella pinea* (Fig. 1B). Since no additional barcodes are available for these isolates, we cannot confirm this hypothesis.

### Whole genome assembly and annotation

Once we determined isolate F5 to be a species of *Anthostomella*, we decided to obtain its full genomic sequence. For that, we extracted high-molecular-weight genomic DNA (Table S2) and subsequently performed long-read sequencing with the new, highly accurate (Q20) chemistry of Oxford Nanopore Technologies (ONT). We generated 1.95 million raw reads (6.12 Gbp) which we filtered to remove all the sequences below 2,000 bp, achieving a final number of 0.8 million reads for a total of 5.17 Gbp. Despite the reduction in sequence information, both the mean read quality and the N50 value increased: from 15.9 to 17.3 and from 6,949 to 8,291, respectively (Table S3). We assembled the filtered reads into the complete genome of *A. pinea* F5, with a total size of 53.7 Mbp (∼80 × coverage). We polished the assembly with Racon and Medaka to produce the final draft of the genome, with a contig N50 value of 5,428,629 and a 97.9% completeness score (BUSCO) [34] (Table 1 and Table S4), indicating that we obtained a highly contiguous and complete genome. Lastly, we used a subset of 10,000 reads to assemble the mitochondrial genome as well, which we fully annotated with the web-based tool GeSeq [35] (Figure 2).

**Figure 2.**
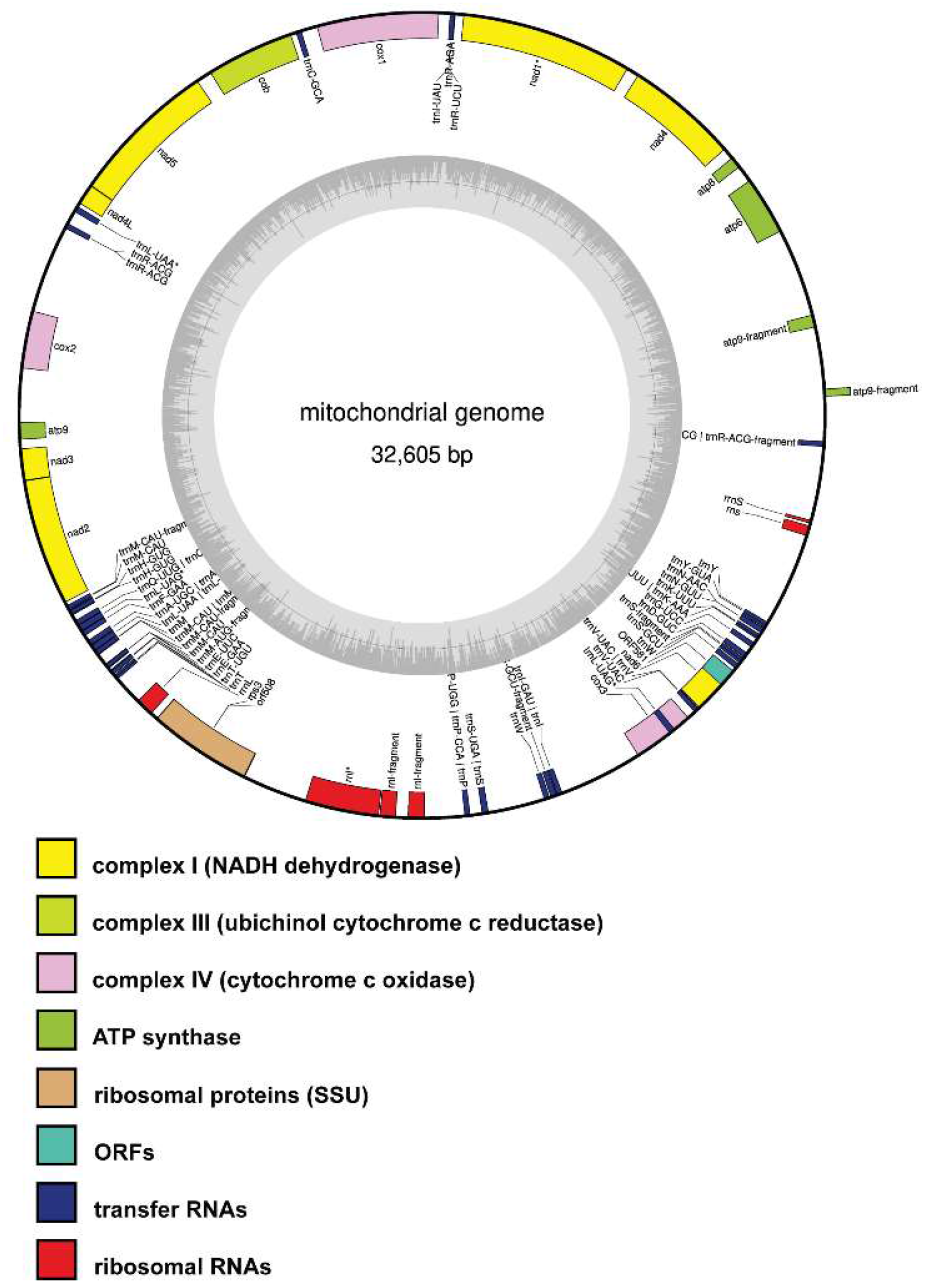
Complete structural and functional annotation of the assembled mitochondrial genome from A. pinea F5.

**Table 1.** Assembly statistics and genome annotations.

|  |  |
| --- | --- |
| Coverage | ~80× |
| # contigs | 19 |
| Largest contig | 8,924,063 |
| Total length | 53,727,737 |
| N50 | 5,428,629 |
| N90 | 2,115,983 |
| L50 | 4 |
| L90 | 10 |
| GC (%) | 51.85 |
| BUSCO (ascomycota) | 98.1% (C) |
| BUSCO (sordariomycetes) | 97.9% (C) |
| Protein coding genes | 14,734 |
| rRNA genes | 48 |
| tRNA genes | 199 |
| Proteins with TM segments | 2923 |
| Proteins with SP | 2109 |
| Proteins with predicted PFAM family/domain | 11,040 |
| CAZymes | 608 |
| P450s | 164 |
| UPOs | 6 |

We then proceeded with structural and functional annotation of the genome, which revealed 14,734 protein coding genes, of which 11,040 could be assigned to PFAM domains/families (Additional file 2). Furthermore, we identified 247 RNA genes, 48 for rRNA and 199 for tRNA, respectively (Table 1). Gene Ontology (GO) analysis revealed that the top 50 terms grouped in the following categories: biological processes (24 %), molecular functions (50 %), and cellular components (26 %). Particularly interesting was the abundance of biotechnologically relevant genes: 171 genes were annotated as “secondary metabolite biosynthetic process”, 229 as “monooxygenase activity”, and 221 as “carbohydrate metabolic process” (Table S5 and Fig 3). Thus, we set out to investigate this type of genes more deeply with specific bioinformatic tools.

**Figure 3.**
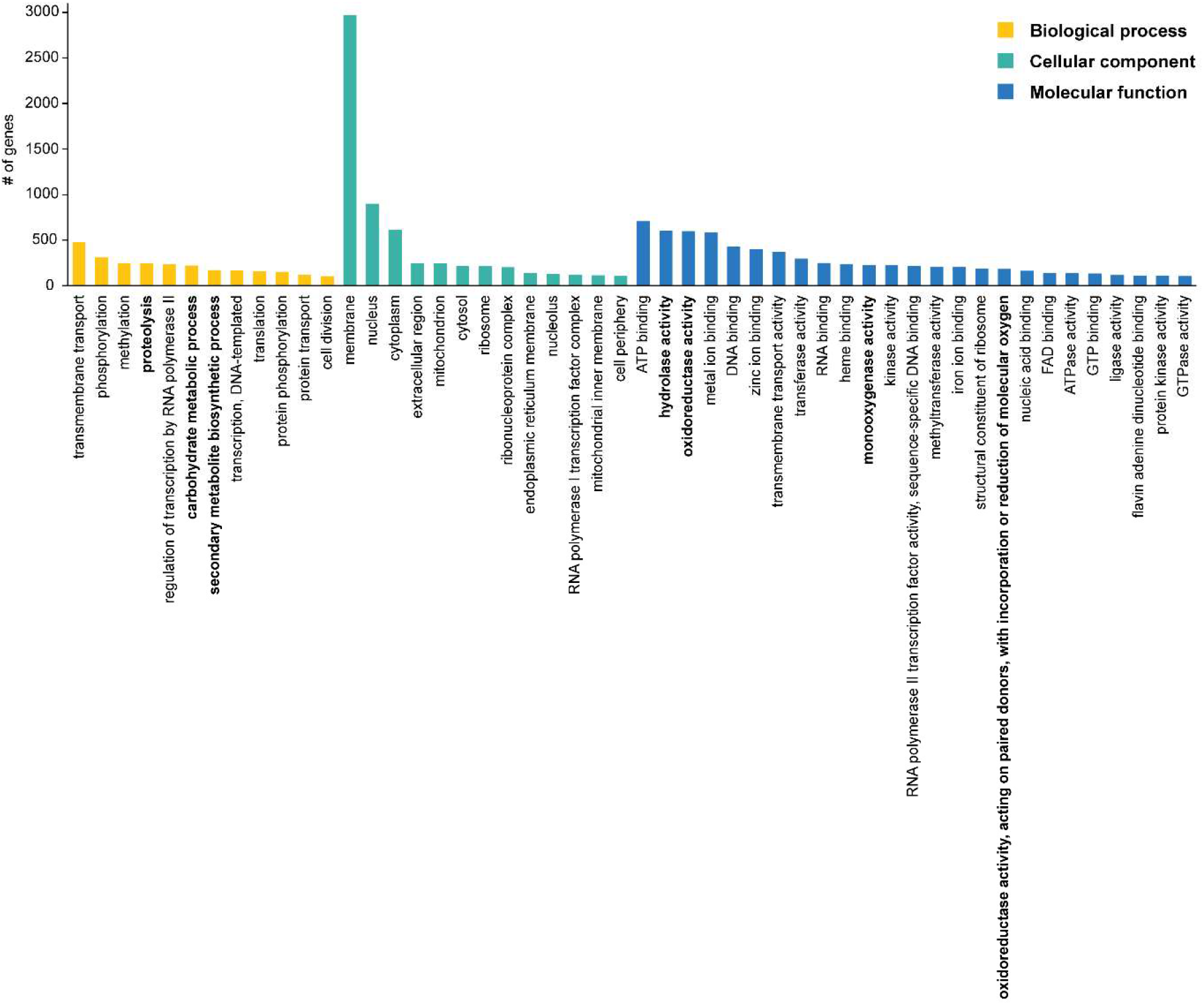
Gene Ontology (GO) analysis of the genome of A. pinea F5 (top 50 terms).

### The genome of *A. pinea* F5 encodes a variety of biotechnologically-relevant enzymes

First, we investigated the CAZyme families in the “carbohydrate metabolic process” to assess the ability of *A. pinea* F5 to degrade lignocellulosic material [36]. We used three different tools, HMMMER, DIAMOND, and dbCAN_sub, incorporated into the webserver dbCAN2 [37]. In total, the genome of *A. pinea* F5 encodes 610 CAZymes. Among the six major CAZymes families [38,39] (glycoside hydrolases (GH), glycosyltransferases (GT), polysaccharide lyases (PL), carbohydrate esterases (CE), auxiliary activities (AA), and carbohydrate-binding modules (CBMs), the GH and AA families were most abundant with 281 and 157 members, respectively (Additional file 2). We then examined the substrate specificity of the CAZymes as predicted by dbCAN_sub [37]. In total, *A. pinea* F5 possesses 146 plant biomass-degrading enzymes, with a particularly high number of enzymes active on xylan (35) and cellulose (49) (Table 2). Interestingly, we also found nine CAZymes that are predicted to be active on lichenin (Additional file 2), a complex glucan that occurs in the cell wall of certain lichen-forming fungi when they are associated with their photosynthetic partner [40]. This might suggest that *A. pinea* F5 is indeed able to colonize lichen tissues and grows as an endo/epilichenic fungus.

**Table 2.**
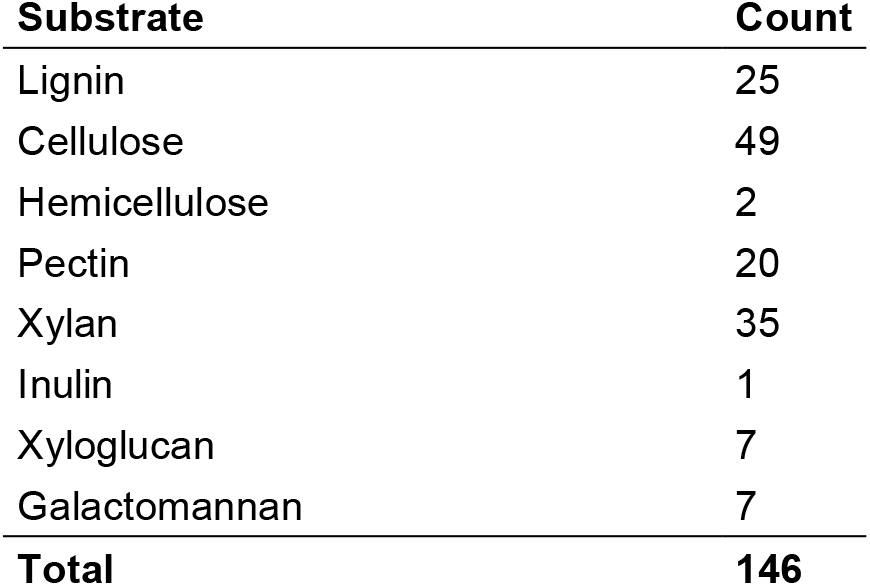
Substrate specificity of CAZymes with predicted plant biomass-degrading activity.

| Substrate | Count |
| --- | --- |
| Lignin | 25 |
| Cellulose | 49 |
| Hemicellulose | 2 |
| Pectin | 20 |
| Xylan | 35 |
| Inulin | 1 |
| Xyloglucan | 7 |
| Galactomannan | 7 |
| **Total** | **146** |

Next, we investigated the abundance of cytochromes P450, an important class of enzymes that play essential roles in fungal metabolism. P450s naturally show very broad substrate scopes, and are capable of performing a wide range of reactions involved in the biosynthesis of bioactive molecules as well as in the degradation of pollutants and xenobiotics, and are therefore considered powerful biocatalysts [41,42]. The bioinformatic analysis revealed that the genome of *A. pinea* F5 encodes 164 P450s distributed across 42 families (Additional file 2). The 4 most represented families (≥10 counts) are CYP3 (21), CYP52 (19), CYP570 (12), and CYP65 (10). Interestingly, at least two of these families (CYP3 and CYP65) are involved in the degradation of xenobiotics [43,44], which may suggest that *A. pinea* can act as “detoxifier” for its environment. For other lichens and lichen-associated fungi it is well established that they fulfill this important ecological role by hyperaccumulating xenobiotics, while remaining unharmed [45,46].

Lastly, we looked at another group of enzymes that has recently made the headlines in the fields of biocatalysis and green chemistry: unspecific peroxygenases (UPOs). So far, UPOs have only been found in the fungal kingdom [47,48] and have attracted considerable attention given their simplicity, stability, and broad range of reactions that they can catalyze. These include, among others, epoxidation, aromatic hydroxylation, ether cleavage, and sulfoxidation, demonstrating the versatility and biocatalytic potential of such enzymes [49]. To search for UPOs in the genome of *A. pinea* F5 we performed two pHMMER analyses using two well characterized UPOs as queries: *Aae*UPO from *Agrocybe aegerita* [50], prototype enzyme of family II (long UPOs); and *Hsp*UPO from *Hypoxylon* sp. EC38, member of family I (short UPO) of which the crystal structure was recently published [51]. Both searches produced the same seven putative UPO sequences with a E-value between 0.0016 and 1.30 × 10^-77^ (Additional file 2). To determine whether these genes might actually encode for UPOs, we performed a multiple sequence alignment of the seven predicted proteins and the two query sequences. All but one showed the typical motifs “PCP” and “ExD” [52] confirming that these proteins are most likely UPOs (Figure 4). Based on length, we can classify three of them as short UPOs (g075280, g022190, g110720) and three as long UPOs (g137960, g143780, g025740).

**Figure 4.**
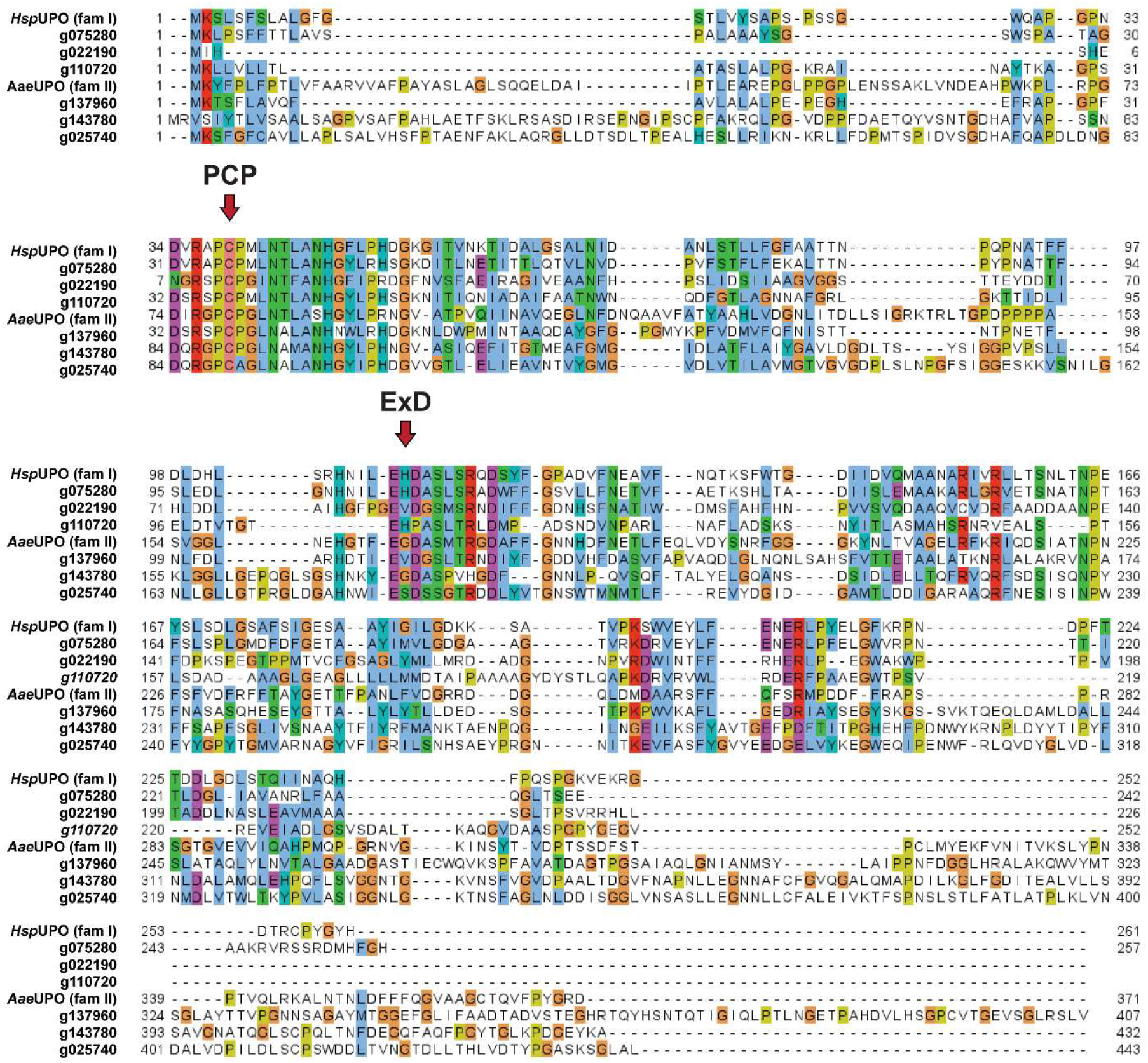
Multiple sequence alignment analysis of putative UPOs from A. pinea F5 and prototype UPOs. The sequences of family I and family II prototype UPOs were retrieved from the Uniprot database. The conserved motifs “PCP” and “ExD” are highlighted by red arrows. The entry of g137950 was removed from the alignment since it lacked the conserved motifs.

Overall, by applying tailored bioinformatic approaches we show that the genome of *A. pinea* contains a wide range of enzymes that might have interesting biotechnological applications, including biocatalysis and bioremediation.

### fungiSMASH predicts an abundance of secondary metabolite biosynthetic gene clusters in the genome of *A. pinea* F5

Fungi are recognized to be prolific producers of secondary metabolites, bioactive molecules with crucial ecological roles and with important applications in medicine and industry [8]. To investigate the secondary metabolism of *A. pinea* F5, we searched its genome for BGCs with the web-based tool fungiSMASH 7 [53]. The software predicted 91 clusters, grouped in the following categories: 23 NRPS (nonribosomal peptide synthetases), 23 fungal RiPP (ribosomally synthesized and post-translationally modified peptides), 22 PKS (polyketide synthases), 14 terpene, 2 indole, and 7 mixed BGCs. Out of these, 71 show no similarities to any known cluster, indicating that they might be involved in unknown biosynthetic pathways (Additional file 3). Among the ones with known similarities, a terpene BGC is predicted to produce the anticancer compound clavaric acid [54], while another terpene BGC and a PKS BGC are predicted to produce the pigments monascorubrin [55] and aurofusarin [56], respectively. These might explain the yellow-to-orange coloration that we observed when performing morphological studies (Fig. 1). Furthermore, *A. pinea* F5 possesses several more BGCs compared to the average for Pezizomycotina (42.8) and the family *Xylariaceae* (71.2) [57]. Such an abundance of BGCs encoded in the genome suggests that this species is likely able to produce a large number of secondary metabolites.

### *A. pinea* F5 is a prolific producer of sesquiterpenoids and sesquiterpene lactones

Despite the large abundance of BGCs, the biosynthesis of secondary metabolites is often regulated and triggered only under specific environmental stimuli [8]. Therefore, we next investigated whether *A. pinea* F5 is actually able to produce secondary metabolites by means of untargeted metabolomics. For that, we grew the fungus in different solid and liquid media for 28 days, and subsequently extracted both extra- and intra-cellular metabolites for LC-MS analysis (Figure S5). The metabolite profile looked almost identical for the different solid media, while the liquid medium showed some differences. To gain further information about the extracts, we processed the raw MS data with MZmine [58] and performed a feature-based molecular networking (FBMN) analysis using the Global Natural Products Social Molecular Networking platform (GNPS) [59,60]. The network was manually curated by deleting multiple nodes with identical mass to charge ratio (m/z) and near-identical RT, artifacts generated during pre-processing with MZmine, and keeping only the node that showed the highest signal. Nodes present in the blank extracts (sterile media) were also deleted. We colored the nodes by occurrence in the extracts of the different media and adjusted the node size according to the m/z, with bigger nodes representing higher m/z. This resulted in a polished network (Additional file 4) consisting of 299 nodes connected by 422 edges, shown in Fig. 5a. By using the additional GNPS tool MolNet Enhancer [61], three of the clusters were annotated as different types of sesquiterpenoids (STs) and sesquiterpenoid lactones (STLs): namely, guaianes (including dioxanes)(I), germacranolides (II), and eudesmanolides (III) (Fig. 5a) (Additional file 5). Guaianes are bicyclic compounds characterized by a 7-membered ring fused to a 5-membered ring. Compounds from II and III are lactones, with the main difference being that germacranolides have a 10-membered ring fused to the lactone function, while in eudesmanolides the 10-membered ring is fused in the middle resulting in two 6-membered rings [62].

**Figure 5.**
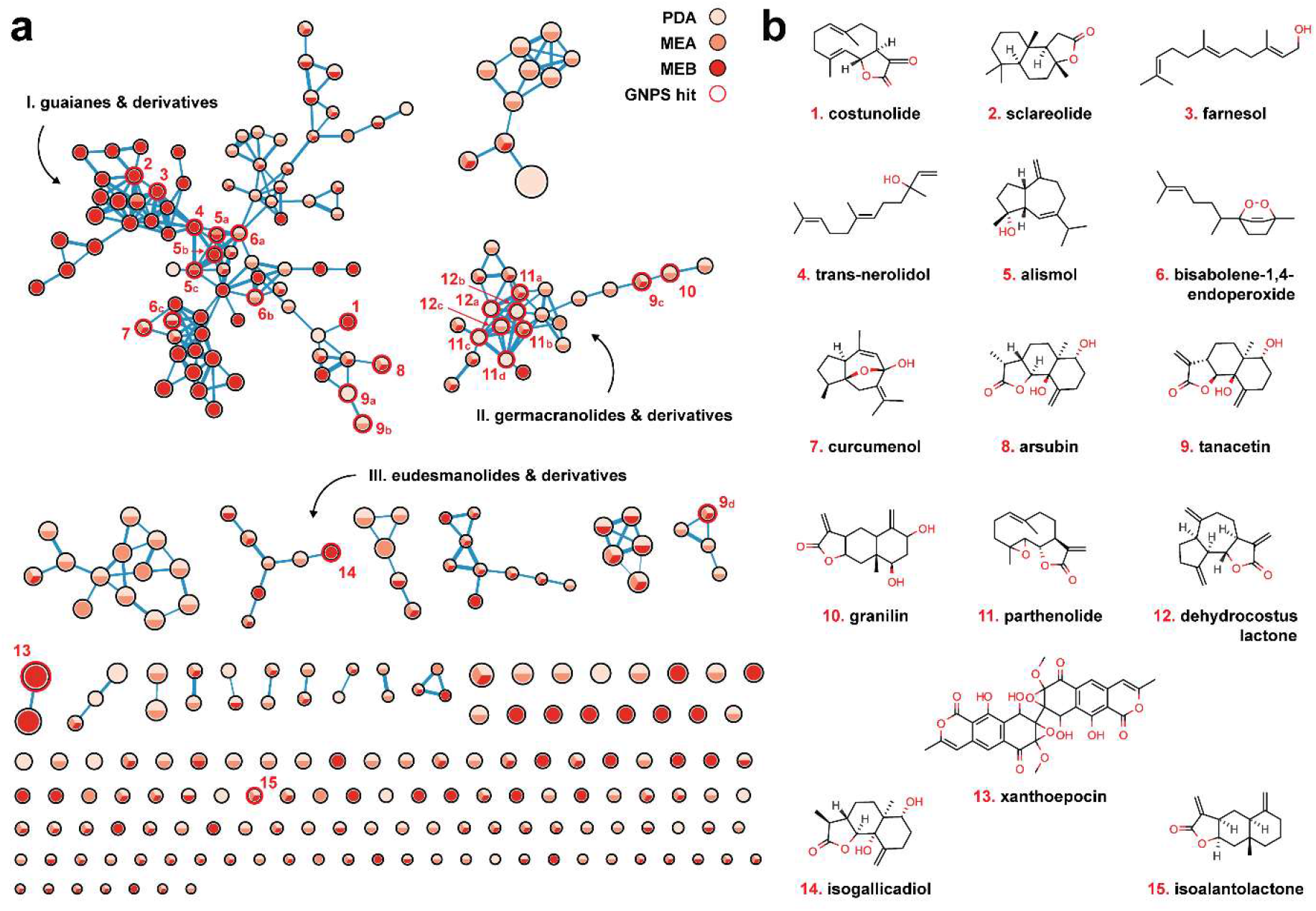
**(a)** Molecular network of extracted secondary metabolites from cultures of A. pinea F5. Nodes are colored based on their occurrence in the extracts from the different media and sized based on the m/z ratio. Hits from the MS/MS GNPS library search are highlighted in red and numbered. **(b)** Chemical structures of the metabolites identified via library search, numbered correspondingly.

Simultaneously with the FBMN workflow, we also ran an MS/MS spectral library search to identify nodes within the network by direct match with spectral data available on the platform. With that, we were able to annotate several nodes within the ST(L)s groups, as well as a few additional nodes in the network (Fig. 5a and 5b) (Table 3).

**Table 3.** Spectral matches from GNPS MS/MS library search. ST = sesquiterpene; STL = sesquiterpene lactone; PK = polyketide.

| Hit # | Compound name | m/z | Adduct | Error (ppm) | Type |
| --- | --- | --- | --- | --- | --- |
| 1 | Costunolide | 233.1535 | [M+H] | -2.57 | Germacranolide STL |
| 2 | Sclareolide | 251.2005 | [M+H] | -2.39 | Eudesmanolide STL |
| 3 | Farnesol | 223.2054 | [M+H] | -3.58 | Linear ST |
| 4 | Trans-nerolidol | 205.1949 | [M-H<sub>2</sub>O+H] | -3.41 | Linear ST |
| 5a,b,c | Alismol | 203.1798 | [M-H<sub>2</sub>O+H] | -0.98 | Guaiane ST |
| 6a | Bisabolene-1,4-endoperoxide | 201.1638 | [M-2H<sub>2</sub>O+H] | -2.49 | Bisabolane ST |
| 6b | Bisabolene-1,4-endoperoxide | 219.1732 | [M-H<sub>2</sub>O+H] | -3.19 | Bisabolane ST |
| 6c | Bisabolene-1,4-endoperoxide | 237.1851 | [M+H] | -1.69 | Bisabolane ST |
| 7 | Curcumenol | 235.1695 | [M+H] | -1.28 | Guaiane ST |
| 8 | Arsubin | 267.1590 | [M+H] | -2.26 | Eudesmanolide STL |
| 9a,b,c,d | Tanacetin | 265.1435 | [M+H] | -1.89 | Eudesmanolide STL |
| 10 | Granilin | 265.1436 | [M+H] | -1.51 | Eudesmanolide STL |
| 11a,b | Parthenolide | 231.1382 | [M-H<sub>2</sub>O+H] | -1.30 | Germacranolide STL |
| 11c,d | Parthenolide | 231.1377 | [M-H<sub>2</sub>O+H] | -3.46 | Germacranolide STL |
| 12a | Dehydrocostus lactone | 231.1376 | [M+H] | -3.89 | Guaianolide STL |
| 12b,c | Dehydrocostus lactone | 231.1382 | [M+H] | -1.30 | Guaianolide STL |
| 13 | Xanthoepocin | 607.1082 | [M+H] | -0.99 | Bis-naphthopyrone PK |
| 14 | Isogallicadiol | 267.1593 | [M+H] | -1.22 | Eudesmanolide STL |
| 15 | Isoalantolactone | 233.1533 | [M+H] | -3.86 | Eudesmanolide STL |

Surprisingly, the library search revealed that the distinction between the three clusters is not as marked as predicted by MolNet Enhancer, given that several hits found in cluster I (**1**, **2**, **7**, **8**) showed the structure of compounds from II and III, and in cluster II at least one hit (**12**) has a typical guaiane-like structure, though fused to a lactone (guaianolide). An important thing to note is that some of the hits are identified multiple times in the network and library search (**5**, **6**, **9**, **11**, **12**), likely due to the presence of several closely related isomers and derivatives that fragment at the MS^1^ level already. This results in different nodes showing identical m/z values and the same MS/MS fragmentation pattern at different RTs (Additional file 7). Because we can’t be certain which of the hits is the actual molecule that was identified through spectral match, we cannot propagate the annotation to related unknown nodes and identify them unambiguously.

Interestingly, other nodes in cluster 1 matched to compounds that do not show neither guaiane-type nor eudesmanolide/germacranolide-type structures. These include farnesol and trans-nerolidol (**3**, **4**), linear sesquiterpenoids likely derived from the hydration of farnesyl cation and its isomer nerolidyl cation, respectively. Lastly, bisabolene-1,4-endoperoxide (**5**) was also matched to three nodes in this cluster, a compound so far only observed in plants that has potential anti-tumor activity [63,64]. Another node in cluster I was matched to curcumenol (**6**), a bioactive molecule isolated from the edible rhizome of *Curcuma zedoaria* (white turmeric) which shows potent anti-inflammatory properties [65]. Both compound **5** and **6** have an endoperoxide bridge within their chemical structure, a feature that is typical of many other bioactive compounds [66], including the well-known antimalaria drug artemisinin [67].

Because of the abundance of STs and STLs in the extracts of *A. pinea* F5, we took a closer look at the terpenoid BGCs encoded in its genome. We performed blastP [68] analyses on the terpene cyclases or synthases in each of the 14 terpene BGCs to gain insight into their putative function (Additional file 8). We observed that eight of these enzymes are closely related to ST synthases. Together with the presence of various tailoring enzymes in the respective BGCs (particularly regions 1.3, 12.3 and 14.6), this confirms that *A. pinea* can produce different ST scaffolds and further diversify them into a broad range of products (Fig. 6). Although we cannot conclusively assign all the library hits to the respective matched nodes, it is evident that *A. pinea* produces a wide variety of sesquiterpenoids with different scaffolds and various functional groups. Most of the compounds matched through library search are bioactive and potentially interesting for pharmaceutical applications [69]. Given that the majority of the nodes in the network are unidentified, we can speculate that the fungus is likely producing completely new compounds, which remain to be explored in the future.

**Figure 6.**
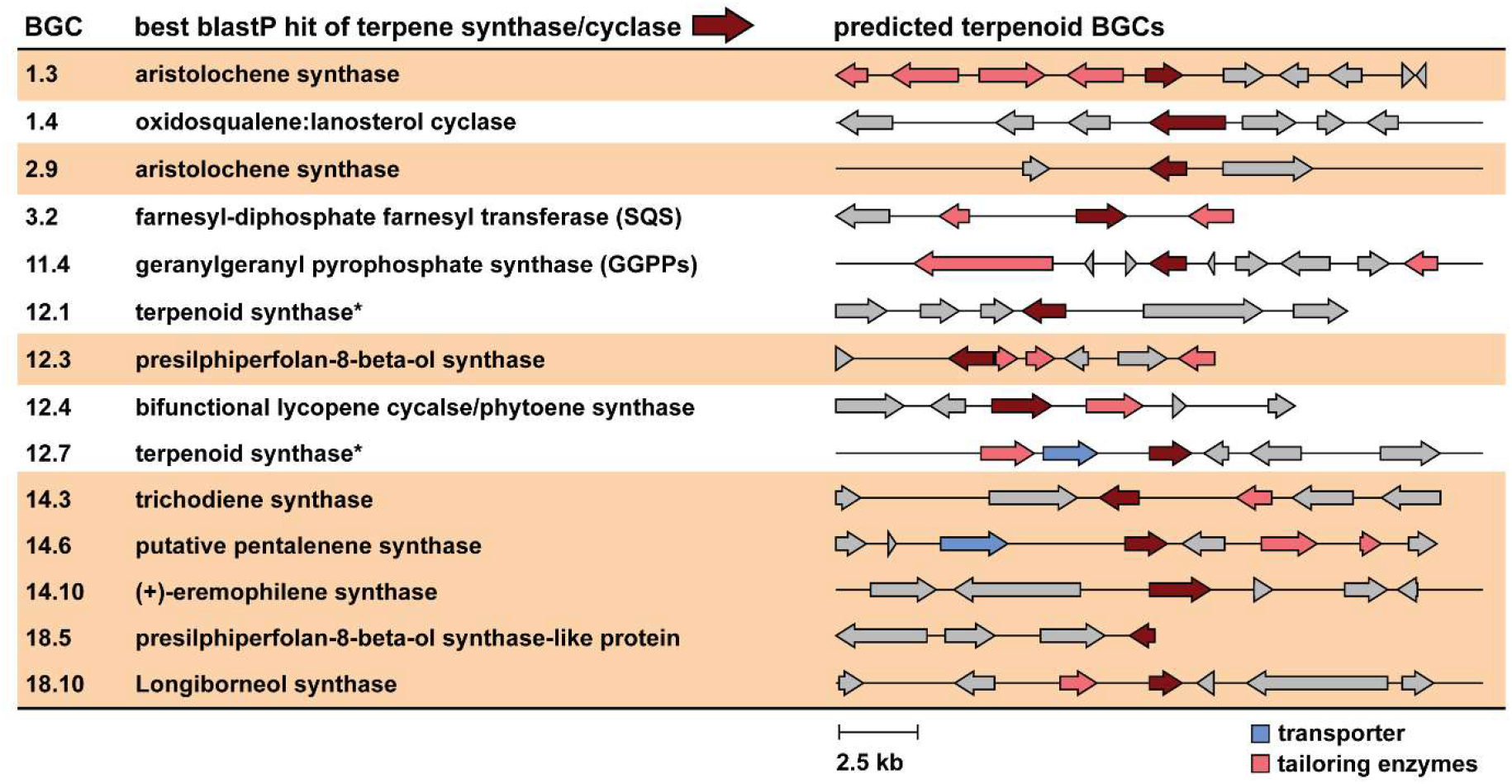
Analysis of terpenoid BGCs as predicted by fungiSMASH v7.0. Clusters highlighted in tan brown correspond to putative STs and STLs biosynthetic gene clusters, based on blastP analysis of their core terpene synthase/cyclase genes.

### *A. pinea* F5 produces the antimicrobial compound xanthoepocin in liquid medium

Besides sesquiterpenoids and sesquiterpene lactones, we identified one other node through spectral library search—node **13**, only found in extracts from liquid MEB medium (Fig. 5a and 5b). The feature was matched to xanthoepocin, a polyketide antibiotic originally isolated from *Penicillium simplicissimum* and commonly found in other *Penicillium* species [70,71]. We also found a related node which showed a negative mass difference of 18.011, consistent with loss of a water molecule from xanthoepocin which happens already at MS^1^ level (Additional files 4 and 6, Fig 5a). Xanthoepocin has a homodimeric structure composed of two napthopyrone scaffolds, typically associated with pigmentation and sporulation in fungi and with a wide range of biological activities [72–74]. Xanthoepocin in particular has been shown to possess potent antibiotic activity against Gram-positive bacteria, including resistant strains of *E. faecium* and MRSA [71]. Informed by these findings, we performed antimicrobial plate assays with the same extracts that we used for the LC-MS analysis. Indeed, we observed antibiotic activity against the indicator Gram-positive species *Micrococcus luteus* for the MEB extract, albeit only when concentrated 10x (Fig. 7). We also performed agar overlay experiments with 28-days old colonies of *A. pinea* on PDA and MEA and we observed complete growth inhibition of *M. luteus* under both conditions (Fig. S6). This antibacterial activity may be caused by yet another differentially produced secondary metabolite. However, it is also possible that the fungus produces xanthoepocin on solid media as well, yet we were not able to detect it via LC-MS analysis. This could be caused by a low extraction efficiency, low starting concentration of the compound, and further degradation of the compound with exposure to light [71]. Because of the scarcity of newly discovered antibiotics and its potency against multi-drug resistant bacteria, xanthoepocin has recently regained attention for the potential development of photodynamic antimicrobial therapies [71]. Therefore, we decided to delve deeper into the genomics data to attempt to identify the BGC involved in its biosynthesis.

**Figure 7.**
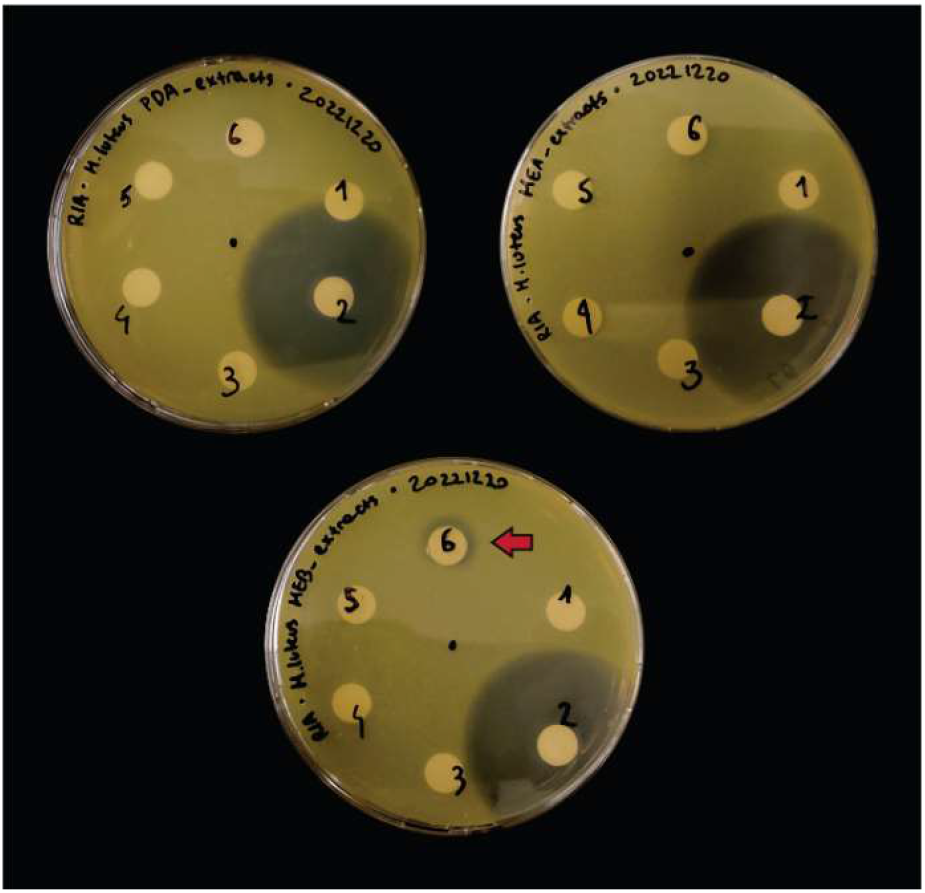
Antibacterial assays of the extracts from A. pinea F5. Top left: PDA extracts; top right: MEA extracts; bottom: MEB extracts. The assays were carried out on LB media overlayed with cultures of indicator strain M. luteus, grown overnight at 30 °C. Numbers correspond to: 1 - 50% MeOH + FA 0.1%; 2 - Ampicillin 20 µg; 3 - medium extract; 4 - medium extract 10x; 5 - fungal culture extract; 6 - fungal culture extract 10x. Detected antibacterial activity (from extract of A. pinea grown on MEB) is highlighted by a red arrow.

### Xanthoepocin is likely biosynthesized by a polyketide BGC that includes a fungal laccase

As discussed in the section above, napthopyrones are widespread across fungi, and the biosynthetic route for many of them is already known [56,72,73]. Generally, the core polyketide structure is synthesized by large, iterative, non-reducing PKS enzymes and further modified by tailoring enzymes (e.g., monooxygenases, or dehydrogenases) to yield the monomeric structure [56]. In the case of bis-naphthopyrones a final coupling step is required to achieve the dimeric structure, which is typically performed by fungal laccases, multicopper oxidases that catalyze intramolecular phenolic coupling via generation of radical intermediates [75,76]. Thus, we analyzed the data from the fungiSMASH prediction searching for a BGC that would encode the necessary enzymes, and successfully identified region 18.3 as the putative xanthoepocin BGC (Fig. 8a, Additional file 3). The cluster encodes all the enzymes predicted to be involved in the biosynthetic route: a NR-PKS (*xaeA*), a flavin-containing monooxygenase (FMO, *xaeO*), an O-methyltransferase (OMT, *xaeM*), an alcohol dehydrogenase (ADH, *xaeD*), and a laccase (*xaeL*). A drug-resistance transporter is present in the region as well (*xaeT*). As a further confirmation, fungiSMASH predicted region 18.3 to be similar to the aurofusarin BGC [56], as well as the BGCs of ustilaginoidins [77] and viriditoxin [78], which are all bis-napthopyrone compounds.

**Figure 8.**
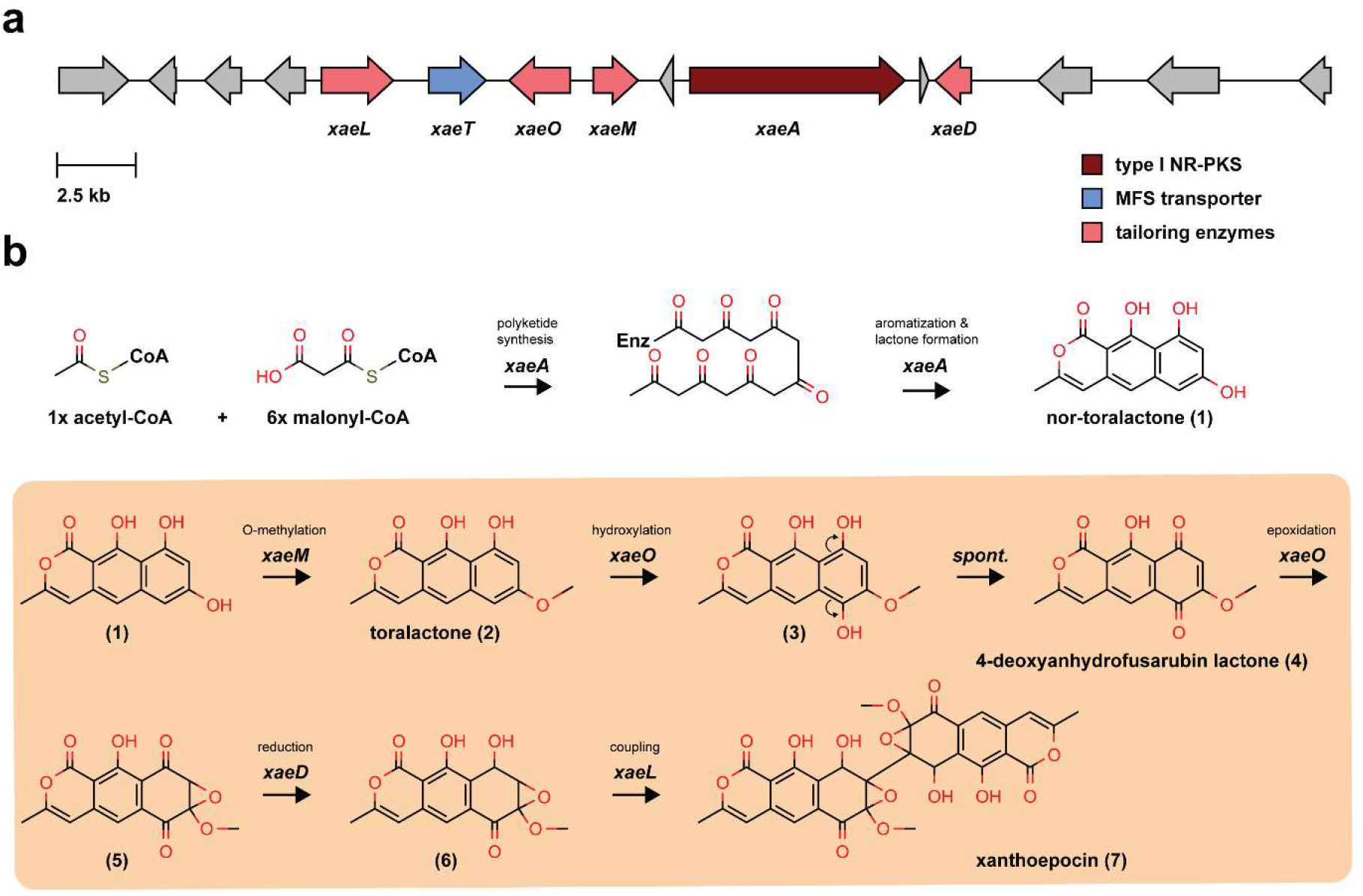
**(a)** Putative BGC (ID 18.3) of xanthoepocin as predicted by fungiSMASH v7.0. **(b)** Proposed biosynthetic pathway of xanthoepocin based on the function of the BGC genes and the chemical structure of the final product. While the biosynthesis of the polyketide scaffold surely occurs as the first step, the order of the remaining steps (brown box) remains to be determined.

Based on the structural features of xanthoepocin and the genes that we identified in the BGC, we propose a step-by-step biosynthetic route (Fig. 8b). The synthesis of the heptaketide scaffold and its subsequent lactonization catalyzed by the PKS are the first steps of the pathway, while the intramolecular coupling is the last. Aside from those, the succession of the remaining steps is only hypothesized, and it could occur in a different order (amber box). Intermediates **1** and **2** are already described in literature [79] as precursors of another naphtopyrone-derived fungal natural product, cercosporin [79,80]—while compound **4** has been prepared synthetically [81]. Indeed, bis-napthopyrone precursors and intermediates often show very similar structures that are differentiated during the late steps of the biosynthetic pathways via functionalization, oxidation/reduction, chain shortening, and coupling reactions [74]. The identification of the xanthoepocin BGC should be validated experimentally (e.g., via gene knockout or heterologous expression) as it could prime efforts to reconstitute the pathway in a model host. This would provide access to higher amounts of compound for biological testing, as well as potentially new variants with engineered desirable properties.

## Conclusions

Fungi are extremely appealing organisms for the discovery of new metabolites and biocatalysts to be used for industrial applications. To date, about 156,000 extant species have been described [82] out of an estimated 2-11 million total species [83]—and only a few thousands genomes have been fully sequenced and are currently available [20,21].

In this work, we report a high-quality whole genome assembly of a wild fungal isolate that we identified as *Anthostomella pinea* based on genetic barcoding. Following structural and functional genome annotation, we used an extensive bioinformatic pipeline to search for enzymes with potential biotechnological applications. First, we identified more than 600 CAZymes, an important group of enzymes with diverse applications in industry [84–87]. We also predicted 164 P450s—powerful biocatalysts that can be used for the regio- and stereospecific oxidation of hydrocarbons [42,88]—and six UPOs, members of a unique family of fungal enzymes that can catalyze a wide range of oxyfunctionalization reactions and that are extremely attractive for their robustness and versatility [47,49,52].

We then explored secondary metabolism in *A. pinea* by combining bioinformatic predictions of biosynthetic gene clusters and molecular networking analyses [53,59,60]. Our investigation revealed that the fungus possesses a rich biosynthetic machinery and can produce a wide variety of small molecules, particularly sesquiterpenoids and sesquiterpene lactones. Through spectral library matches, we putatively identified 14 sesquiterpenes and sesquiterpene lactones and one bis-napthopyrone polyketide with antibiotic activity which is currently being re-evaluated for biological testing [71]. For the latter, we also identified the putative BGC and proposed a step-by-step biosynthetic mechanism based on its chemical structure and the predicted activities of the BGC enzymes.

Overall, our compressive analysis showcases the rich biosynthetic capacity of the fungus *A. pinea*, a species thus far underexplored. Our findings pave the way to explore targeted approaches for the discovery and production of bioactive molecules, such as genome editing and heterologous expression of genes and BGCs of interest. Lastly, this work may prompt further investigation into the genomic and metabolic diversity of phylogenetically related fungi.

## Materials and Methods

### Sampling of lichen thalli

One large thallus of *Letharia lupina* was collected along the Cheney Wetland Trail, Cheney, Spokane County, Washington, USA (47.483392, −117.553087). The lichen was growing on a *Pinus ponderosa* branch along the edge of a pond and appeared robust and healthy (that is, lacking large galls, perithecia, or discoloration that would indicate parasite infection or tissue necrosis). Collection was completed with a sterile nitrile glove and the thallus was placed in a paper bag. After drying in the bag at room temperature on the lab bench in the original collection bag for ∼48 hours it was shipped to Groningen, The Netherlands.

### Fungal isolation and preliminary barcoding

The specimen of *A. pinea* was isolated from section of dry thalli of the lichen *Letharia lupina* using a modified version of the Yamamoto method [27]. Briefly, a sample of dry thallus was washed three times with sterile ddH_2_O under a biosafety cabinet, after which it was damp-dried with sterile cheesecloth and cut into ∼1 cm long sections using a metal scalpel. These sections were incubated in sterile ddH_2_O for 4 weeks in the dark at room temperature (18-23 °C) before being inspected under a stereoscopic microscope. When filamentous growth was observed at either end of one of the sections, this was excised and placed on Malt-Yeast Extract Agar (MYA) plates (malt exctract 20 g/L; yeast extract 2 g/L; microagar 15 g/L). In total, 10 isolates were obtained from as many growth points. The plates with the isolates were incubated at room temperature in the dark for up to six weeks to allow for sufficient growth of all the isolates.

To obtain template DNA for preliminary ITS barcoding, small sections of mycelia of approximately 4 mm^2^ were scraped from the growing edges of the isolates and placed in 20 µL of DNA dilution buffer (Tris-HCl 10 mM; NaCl 10 mM, EDTA 1 mM, pH = 7.5) in 0,2 mL PCR tubes. The suspensions were then incubated at 95 °C for 10 minutes immediately followed by 2 minutes of incubation on ice, after which they were centrifuged at max speed for 1 minute to pellet the mycelia. The supernatant was transferred to clean tubes for storage, and 1 µL was used for direct colony PCR with with the ITSF1 (5′-CTTGGTCATTTAGAGGAAGTAA-3′) and ITS4 (5′-TCCTCCGCTTATTGATATGC-3′) primers. The PCR reaction was performed in a thermal cycler with 2x Plant Phire Master Mix (ThermoFisher Scientific, Waltham, MA, United States) with the following program: 5 min at 98 °C; 34 cycles of 10 s at 98 °C, 10 s at 60 °C, 40 s at 72 °C; 5 min at 72 °C. Two microliters of the PCR product were taken for 1% agarose gel electrophoresis analysis to confirm successful amplification of the ITS region. The PCR product was then purified using the QIAquick PCR Purification Kit (Qiagen, Venlo, the Netherlands) and sent to Macrogen Europe (Amsterdam, the Netherlands) for Sanger sequencing. The resulting sequences were analyzed with the Basic Local Alignment Search Tool (BLAST) [89] against the nucleotide collection of the National Center for Biotechnology Information (NCBI) to identify the best match for the fungal isolate based on %ID, coverage, and E-value. The original plates with isolates F5-7 (putatively identified as *A. pinea*) were stored at 4 °C and used for inoculation of fresh medium.

### ITS-based species assignment

For species assignment, we extracted genomic DNA from 40 mg of mycelium freshly scraped from an agar plate, using the Nucleospin Microbial DNA kit (Bioké, Leiden, the Netherlands) in combination with NucleoSpin Bead Tubes Type C (Bioké, Leiden, the Netherlands) for tissue disruption in a MM 301 vibratory mill (Retsch GmbH, Haan, Germany). The extraction was carried out according to manufacturer’s instructions except for the disruption time, where 2 cycles of 1 min each were used. Next, a PCR reaction was performed with 1 µL of extracted gDNA and the primers ITSF1 and ITS4, in a thermal cycler with a 2 × Q5 PCR master mix (New England Biolabs, Ipswich, MA, USA). The following program was used: 1 min at 98 °C, 30 cycles of 10 s at 98 °C, 15 s at 55 °C, 20 s at 72 °C, 5 min at 72 °C. The PCR product was visualized on agarose gel, purified, sequenced, and analyzed with BLAST as describe above. The 25 best hits and the ITS sequences of other 18 *Anthostomella* strains obtained from the NCBI nucleotide database were compared by constructing a maximum-likelihood phylogenetic tree using MEGA 11 [90]. First, the sequences were aligned with the MUSCLE algorithm with default settings [91], then the tree was built using standard settings and number of bootstrap replications of 100.

### Morphological characterization

For morphological characterization, *A. pinea* was inoculated on malt extract agar (MEA: malt etraxct 30 g/L; peptone 5 g/L; microagar 15 g/L), potato dextrose agar (PDA, 39 g/L)), yeast extract sucrose agar (YES: yeast extract 4 g/L; sucrose 20 g/L; KH_2_PO_4_ 1 g/L; MgSO_4_ · 7H_2_O 0,5 g/L; microagar 15 g/L), Czapek yeast autolysate agar (CYA: sucrose 30 g/L; yeast extract 5 g/L; NaNO_3_ 3 g/L; K_2_HPO_4_ 1 g/L; KCl 0,5 g/L; MgSO_4_ · 7H_2_O 0,5 g/L; FeSO_4_ · 7H_2_O 0,01 g/L; microagar 15 g/L), and dichloran-glycerol medium (DG18: peptone 5 g/L; glucose 10 g/L; KH_2_PO_4_ 1 g/L; MgSO_4_ 7H_2_O 0,5 g/L; glycerol 180 g/L; dichloran 0,002 g/L). The plates were incubated at 20 °C and the growth of *A. pinea* was monitored regularly until the colonies stopped growing after 4 weeks, at which points the plates were photographed using a Nikon D7500 camera (Nikon, Minato City, Tokyo, Japan) coupled to a Sigma 17-50mm F/2.8 EX DC OS (Sigma Corporation, Kawasaki, Kanagawa, Japan). The colonies were incubated for further 8 weeks to assess whether any additional morphological change would occur, but none was observed. For morphological comparisons (Fig. S1), *A. pinea* CBS128205 was purchased from the strain collection of the Westerdijk Institute of Fungal Biodiversity (Utrecht, the Netherlands). Both isolate F5 and *A. pinea* CBS128205 were grown on MEA for 4 weeks at 20 °C, before being imaged as described above.

### Extraction of HMW gDNA

The fungus was grown in 25 mL malt extract broth (MEB: malt etraxct 30 g/L; peptone 5 g/L) at 20 °C in the dark in static conditions for 28 days. The mycelium was then collected by filtration with sterile Miracloth (MilliporeSigma, Burlington, MA, USA), washed with ∼20 mL sterile Milli-Q water, snap-frozen in liquid N_2_, and lyophilized overnight using a Lyovapor L-200 (Buchi AG, Flawil, Switzerland). The dried biomass was finally stored at −20 °C.

DNA extraction was carried out with 37 mg of freeze-dried mycelium using the Nucleospin Microbial DNA kit in combination with 3 mm tungsten carbide beads (Qiagen) for tissue disruption in a MM 301 vibratory mill, with significant modifications to the manufacturer’s protocol. Briefly, disruption time was reduced to two cycles of 5 s to minimize DNA shearing, with 30 s pause in between to allow the sample to cool down; each vortexing step was replaced by gentle flicking or mixing by inversion; washing steps with buffer BW was repeated four times, followed by four washes with buffer B5; elution was carried out in 60 µL of pre-warmed EB buffer (∼55 °C), and the eluate was then re-applied to the column, incubated for one additional minute and re-eluted to maximize DNA recovery. The integrity of the purified DNA was assessed by agarose gel electrophoresis, while quality control prior sequencing was performed using NanoDrop ND-1000 (ThermoFisher, Waltham, MA, USA), Qubit 3.0 (Invitrogen, Waltham, MA, USA), and the Qubit dsDNA HS Assay Kit (Invitrogen).

### Library preparation and sequencing

For long-read sequencing, the genomic DNA was prepared using the ligation sequencing kit SQK-LSK112 (Oxford Nanopore Technologies, Oxford, United Kingdom) according to the manufacturer’s guidelines. Briefly, genomic DNA (1020 ng) was subjected to end repair, 5’ phosphorylation and dA-tailing by NEBNext FFPE DNA Repair and NEBNext Ultra II End prep modules (New England Biolabs) and purified with AMPure XP (Beckman Coulter, Pasadena, CA, USA) magnetic beads. The sequencing adaptors were ligated using the NEB Quick T4 DNA Ligase (New England Biolabs) in combination with ligation buffer (LNB) from the SQK-LSK112 kit. The library was finally cleaned up using the long fragment buffer (LFB) and purified with AMPure XP magnetic beads. For sequencing, 12 µL of library (∼400 ng) were loaded into a primed FLO-MIN112 (ID: FAT29688) flow cell on a MinION device for a 42-hour run. Data acquisition was carried out with MinKNOW software v22.05.5 (Oxford Nanopore Technologies).

### Read processing and whole genome assembly

The raw reads were basecalled using Guppy v6.0.7 (Oxford Nanopore Technologies) in GPU mode using the dna_ r10. 4_ e8.1_sup.cfg model (super accuracy). The basecalled reads were subsequently filtered to a minimum length of 2 kb and trimmed by 20 nt at both ends using NanoFilt v2.8.0 [92]. NanoPlot v1.40.0 [92] was used to evaluate the filtered reads. Assembly was performed using Flye v2.9-b1778, using the parameters *--nano-hq and --read-error 0.03*, optimized for Q20+ chemistry and super accurate basecalling. The quality of the genome assembly was evaluated using QUAST v5.1.0rc1 [93] and Bandage v0.8.1 [94] (Fig. S4). The draft assembly was subsequently polished in two rounds. First, Racon v1.4.10 [95] was used with the following settings: *−m 8 −x −6 −g −8 −w 500*, optimized for combined use with Medaka. Second, Medaka 1.6.0 was used on the polished version with the r104_e81_sup_g5015 model. The completeness of assemblies was evaluated using BUSCO v5.3.2 (ascomycota_odb10 and sordariomycetes_odb10 datasets) [34]. To assemble the mitochondrial DNA, we subsampled the filtered reads to generate a set of 10,000 reads and then used Flye v2.9-b1778 as described above. The only circular contig obtained was analyzed with BLAST against the NCBI nucleotide collection to confirm its organellar origin. Lastly, the mitochondrial genome was annotated using GeSeq [35] using default settings.

The complete sequencing data and genome assembly for this study have been deposited in the European Nucleotide Archive (ENA) at EMBL-EBI under accession number PRJEB67537.

### Structural and functional genome annotation

Genome annotation was carried out on the polished assembly using the online platform Genome Sequence Annotation Server v6.0, which provides a pipeline for whole genome structural and functional annotation [96]. Standard settings were used unless otherwise mentioned. In brief, low complexity regions and repeats were masked using RepeatModeler v2.0.3 and RepeatMasker v4.1.1 [97], setting the DNA source to ‘Fungi’. The newly generated masked consensus sequence was used for *ab initio* gene prediction using the following tools: (I) Augustus v3.4.0 [98], selecting *Fusarium graminearum* as a trained organism; (II) GeneMarkES v4.48 [99]. For homology-based prediction, the NCBI reference transcript and protein databases for Fungi were searched, using (III) blastn v2.12.0 [89] and (IV) DIAMOND v2.0.11[100], respectively. To generate the final consensus gene model EvidenceModeler v1.1.1 [101] was used on the above-mentioned predictions, weighted as follows: (I)–five, (II)-five, (III)-ten, (IV)-ten. For prediction of proteins with PFAM domains, the Pfam module within GenSAS was used with the following parameters: E-value sequence: 1; E-value Domain: 10. The tRNA and rRNA were predicted using tRNA scan-SE v2.0.11 (“check for pseudogenes = OFF”) [102] and barrnap v0.9 [103], respectively. Prediction of proteins with signal peptides and/or transmembrane domains was carried out on the web server Phobius [104]. Gene Ontology (GO) annotation was performed using the webtool PANNZER2 [105].

### Prediction of secondary metabolites BGCs, CAZymes, P450s, and UPOs

Secondary metabolite biosynthetic gene clusters were identified using the fungal suite of antiSMASH web server v7.0 with the default settings [53]. To annotate CAZymes, the web server dbCAN2 was used [37]. The integrated HMMER, DIAMOND, and HMMER-dbCAN-sub tools were used on the total proteome of *A. pinea*. The three outputs were automatically combined, and CAZymes predicted only by 1/3 of the tools were removed to improve the annotation accuracy. The substrate specificity of the final hits was extracted from the individual results of HMMER-dbCAN-sub.

For the prediction of P450s, the BLAST tool of biocatnet CYPED 6.0 [106] was used, with E-value cutoff set at 1.0 x 10^-10^. To identify putative UPOs, the query sequences of the prototype enzymes AaeUPO [50] and HspUPO [51] were retrieved from the Uniprot database [107], and submitted to phmmer (HMMER v3.3.2) to search against the total proteome of *A. pinea*. The cutoff was set at E-value of 0.01. The multiple sequence alignment analysis between the putative UPOs from *A. pinea* and the prototype AaeUPO and HspUPO was performed with MEGA 11 [90] using the MUSCLE algorithm [91] with default settings, and visualized in Jalview [108].

### Extraction of secondary metabolites and HRMS-MS^2^ analysis

Fungal mycelium was transferred from storage plates to PDA or MEA plates. For inoculation in liquid MEB, mycelium of *A. pinea* was scraped from a storage plate, coarsely ground into 1 mL of MEB in an 1.5 mL microcentrifuge tube with a pipette tip, and then transferred to 25 mL of medium. The plates and flasks were incubated at 20 °C for 28 days alongside empty PDA, MEA plates and MEB-containing flasks to be used as controls. For extraction of SMs, the whole agar pads (agar and mycelium) were cut into pieces and transferred to 50 mL polypropylene tubes, then extracted twice with 25 mL of 9:1 ethyl acetate-methanol (v/v) with 0.1% formic acid and sonicated in a sonication bath for one hour for each extraction. For the liquid cultivations, the entire content of the flasks was collected in 50 mL PP tubes and extracted as above. Prior extraction, all samples were spiked with 10 µL caffeine standard solution with a concentration of 10 mg/mL to validate the extraction procedure. The dried extracts were resuspended in 1 mL of 1:1 MeOH-MilliQ water (v/v) supplemented with 0.1% formic acid (FA), filtered with 0.45 μm PTFE filters, and stored at −20 °C until further analysis.

HR-LC-MS/MS analysis was performed with a Shimadzu Nexera X2 high performance liquid chromatography (HPLC) system with binary LC20ADXR coupled to a Q Exactive Plus hybrid quadrupole-orbitrap mass spectrometer (Thermo Fisher Scientific, Waltham, MA, USA). A Kinetex EVO C18 reversed-phase column was applied for HPLC separations (100 mm × 2.1 mm I.D., 2.6 μm, 100 Å particles, Phenomenex, Torrance, CA, USA), which was maintained at 50 °C. The mobile phase consisted of a gradient of solution A (0.1% formic acid in MilliQ water) and solution B (0.1% formic acid in acetonitrile). A linear gradient was used: 0–3 min 5% B, 3–51 min linear increase to 90% B, 51–55 min held at 90% B, 55–55.01 min decrease to 5% B, and 55.01–60 min held at 5% B. The injection volume was 2 µL, and the flow was set to 0.25 mL·min−1. MS and MS/MS analyses were performed with electrospray ionization (ESI) in positive mode at a spray voltage of 3.5 kV, and sheath and auxiliary gas flow set at 60 and 11, respectively. The ion transfer tube temperature was 300 °C. Spectra were acquired in data-dependent mode with a survey scan at m/z 100–1500 at a resolution of 70,000, followed by MS/MS fragmentation of the top 5 precursor ions at a resolution of 17,500. A normalized collision energy of 30 was used for fragmentation, and fragmented precursor ions were dynamically excluded for 10 s.

### Data processing and molecular networking analysis

The raw MS/MS data files were converted to mzML format using the format conversion utility MSConvert from the ProteoWizard suite [109]. Binary encoding precision was set at 32-bit and zlib compression was set as off. Data was centroided on both MS^1^ and MS^2^ level using the peak picking filter (algorithm set to “Vendor”). The data files were then pre-processed with MZMine v2.53 [58] to generate a .csv feature list and MS/MS .mgf file. The converted MS/MS data, feature list, and the .mgf files were subsequently uploaded to GNPS [59] using the WinSCP tool [110].

A molecular network was generated using the FBMN workflow from the GNPS website [60], version release 28.2. The precursor ion mass tolerance and MS/MS fragment ion tolerance were set to 0.01 Da and 0.02 Da, respectively. Edges were filtered to have a cosine score above 0.7 and more than six matched peaks. Further, edges between two nodes were kept in the network if each of the nodes appeared in the other’s respective top 10 most similar nodes (task ID: <u>gnps.ucsd.edu/ProteoSAFe/status.jsp?task=9d0fe25c59bc4efc876c9a638807a05d</u>, generated on 6 March 2023) (Additional file 4). All mass spectrometry data have been deposited on GNPS under the accession number MassIVE ID: MSV000093120. The molecular network was visualized and curated in Cytoscape version 3.9.1 [111]. Briefly, for nodes with identical m/z ratio and near-identical RT (≤ 0.1 min), only the node with the highest signal intensity and peak area was kept. Nodes occurring in the blanks (PDA, MEA, MEB extracts) were considered background and omitted from the polished network. The network was submitted to the GNPS tool MolNetEnhancer [61] version release 22 to annotate compound families, with default settings (task ID: gnps.ucsd.edu/ProteoSAFe/status.jsp?task=ffb0040546004a119b34be5dc2869aaa, generated on 6 March 2023) (Additional file 5).

The spectra in the network were then searched against the GNPS spectral libraries. Matches were kept with a score above 0.7 and at least six matched peaks. The data was also analyzed by the GNPS molecular library search V2 pipeline, version release 28 [112]. The precursor ion mass tolerance and fragment ion tolerance were set to 0.01 and 0.02 Da, respectively. The minimal matched peaks were set to eight, and the score threshold was set to 0.7 (task ID: <u>gnps.ucsd.edu/ProteoSAFe/status.jsp?task=14dff366901b437394e4d0feae71ff5d</u>, generated on

7 July 2023). Several matched annotations in the library search mode were manually added to the molecular network (additional files 4 and 6). Lastly, the MS/MS data (.mgf file) were processed with the bioinformatic tool Sirius v5.7.1 for compound annotation [113]. The predictions of the chemical formula for each node are included in the molecular network (additional file 4).

### Antimicrobial plate assays

For the whole-colony plate assays, *A. pinea* was inoculated in the center of MEA or PDA plates and grown at 20 °C for 28 days in the dark. *M. luteus* ATCC 10240 was grown overnight at 30 °C, 200 rpm, in 5mL 2x yeast extract tryptone medium (2x YT: tryptone 16 g/L, yeast extract 10 g/L, NaCl 5 g/L). The overnight culture was then diluted 1:100 in warm (∼60 °C) 50 mL 2x YT supplemented with 0.6% agar, mixed well and immediately overlayed on the agar plates (three mL each) with the colonies of *A. pinea* and corresponding controls. The plates were incubated overnight and inspected for antibacterial activity.

To assay the antimicrobial activity of the extracts, *M. luteus* ATCC 10240 was grown overnight and diluted 1:100 in 2x YT + 0.6% agar as described above. Three mL were immediately plated on LB plates (peptone 10 g/L, NaCl 10 g/L, yeast extract 5 g/L, microagar 15 g/L) and left to dry for ∼15 minutes. Six sterile diffusion disks (10mm ø, MilliporeSigma, Burlington, MA, USA) were then gently positioned on the surface of the agar. Each disk was soaked with either 10 µL of fungal extract, fungal extract 10x, medium extract, medium extract 10x, 50% MeOH + FA 0.1% (blank solvent), and ampicillin 2 mg/mL (20 µg) as positive control. The plates were incubated overnight and inspected for antibacterial activity.

## Declarations

### Ethics approval and consent to participate

Not applicable.

### Consent for publication

Not applicable.

### Data availability

All data generated or analyzed during this study are included in the main article and additional files. The sequencing data and genome assembly for this study have been deposited in the European Nucleotide Archive (ENA) at EMBL-EBI under the accession number PRJEB67537. The mass spectrometry data have been deposited on GNPS under the accession number MassIVE ID: MSV000093120.

### Author contributions

RI conceived the study with input from KH and JLA; JLA collected lichen specimen; RI, THe and THackl performed genome sequencing and analysis; RI performed all fungal cultivations and metabolomics studies; KH acquired funding and supervised data collection and analysis, RI and KH wrote the manuscript with input from all authors. All authors have read the final version of the manuscript and agreed to its publication.

#### Acknowledgements

The authors are grateful to Xiao Li for discussions on the molecular networking analysis, and the Interfaculty Mass Spectrometry Center of the University of Groningen and the University Medical Center Groningen for their services in high resolution tandem mass spectrometry.

## Additional files

**Additional file 1:** tables S1-S5, and figures S1-S6. **Additional file 2:** structural and functional annotation of the genome of *A. pinea* F5. **Additional file 3:** BGC prediction by fungiSMASH7 on the genome of *A. pinea* F5. **Additional file 4:** molecular network of *A. pinea* F5 extracts and annotation by Sirius (Cytoscape session file). **Additional file 5**: molecular network of *A. pinea* F5 extracts annotated with MolNetEnhancer (Cytoscape session file). **Additional file 6:** list of hits from GNPS MS/MS library search of *A. pinea* extracts. **Additional file 7:** Extracted ion chromatograms, MS and MS/MS spectra of annotated nodes from the molecular network. **Additional file 8:** results of blastP search on core synthase/cyclase genes from predicted terpene BGCs in the genome of *A. pinea* F5.

