## Additional file 1 for "Genome sequencing and molecular networking analysis of the wild fungus *Anthostomella pinea* reveal its ability to produce a diverse range of secondary metabolites"

**Contents**

Supplementary tables………………………………………………………………………..1

Supplementary figures……………………………………………………………………….5

**Supplementary tables**

**Table S1. Results of the blastn analysis of the ITS region of *A. pinea* F5 (first 25 hits based on %ID).**

| **Accession** | **% ID** | **E-value** | **Bit score** | **Taxon** |
| --- | --- | --- | --- | --- |
| GQ153162.1 | 99.638 | 0 | 1009 | Sordariomycetes sp. 11216 |
| KP992032.1 | 99.269 | 0 | 987 | Sordariomycetes sp. voucher ARIZ YLH0051 |
| KP992078.1 | 99.259 | 0 | 974 | Sordariomycetes sp. voucher ARIZ YLH0105 |
| HQ599578.1 | 98.338 | 0 | 1160 | *Anthostomella pinea* culture collection CBS 128205 |
| MF327371.1 | 98.316 | 0 | 1040 | *Anthostomella pinea* strain DNA105 |
| HM123694.1 | 98.195 | 0 | 966 | *Xylariaceae* sp. ARIZ AZ1047 |
| KX096656.1 | 98.151 | 0 | 1037 | *Anthostomella pinea* isolate RP139_2_1 |
| OK576226.1 | 98.151 | 0 | 1037 | *Anthostomella pinea* isolate SD11 |
| KU663949.1 | 98.136 | 0 | 1027 | uncultured *Anthostomella* clone AK100 |
| KJ406993.1 | 98.085 | 0 | 817 | *Anthostomella pinea* strain C14 |
| KJ406991.1 | 98.085 | 0 | 817 | *Anthostomella pinea* strain CL199 |
| KT291430.1 | 97.749 | 0 | 915 | *Anthostomella pinea* strain FMY29 |
| MW447070.1 | 97.663 | 0 | 1027 | *Anthostomella pinea* strain FeC56 |
| MF327370.1 | 97.619 | 0 | 1007 | *Anthostomella pinea* strain DNA51 |
| MH374419.1 | 97.505 | 0 | 821 | *Anthostomella pinea* strain F317 |
| MT153620.1 | 97.482 | 0 | 948 | *Anthostomella* sp. strain GLMC451 |
| MG543933.1 | 97.388 | 0 | 911 | *Anthostomella* sp. isolate EP22 |
| AY590787.1 | 97.323 | 0 | 887 | *Xylaria* sp. olrim65 |
| FN435792.1 | 97.126 | 0 | 880 | *Xylaria* sp. agrAR275 |
| FN435766.1 | 96.85 | 0 | 846 | *Xylaria* sp. agrAR229 |
| FN435726.1 | 96.78 | 0 | 880 | *Xylaria* sp. agrAR143 |
| FN435657.1 | 96.008 | 0 | 850 | *Xylaria* sp. agrAR040 |
| HM123573.1 | 95.824 | 0 | 734 | *Nemania* sp. ARIZ AZ0890 |
| KP991846.1 | 95.662 | 0 | 704 | Sordariomycetes sp. voucher ARIZ SR0070 |
| GQ153193.1 | 95.246 | 0 | 929 | Sordariomycetes sp. 11262 |

**Table S2. Quality metrics of HMW gDNA from *A. pinea*.**

| **Metrics** | **#** |
| --- | --- |
| Qubit conc. (ng/µL) | 60.6 |
| Nanopore conc. (ng/µL) | 86 |
| Output (μg) | 3.1 |
| A260/280 ratio | 1.91 |
| A260/230 ratio | 2.06 |
| NanoDrop/Qubit conc. ratio | 1.42 |
| Input for library preparation (μg) | 1 |
| Output after DNA repair and end-prep (μg) | 0.93 |
| Output after adapter ligation and clean-up (μg) | 0.52 |

**Table S3. Analysis of reads generated by MinION sequencing**

| **Stats** | **Raw** | **Filtered (>2000bp)** |
| --- | --- | --- |
| Mean read length | 3,135.70 | 6,441.8 |
| Mean read quality | 15.9 | 17.3 |
| Mean read accuracy | 97.43 | 98.14 |
| Median read length | 1,425.00 | 4,662 |
| Median read quality | 15.8 | 17.6 |
| Median read accuracy | 97.37 | 98.26 |
| Number of reads | 1,951,487.00 | 802,004 |
| Read length N50 | 6,949.00 | 8,291 |
| STDEV read length | 4,688.60 | 5,838.2 |
| Total bases | 6,119,362,157.00 | 5,166,368,880 |

**Table S4. QUAST analysis of the *A. pinea* whole genome assembly.**

| **Stats** | **#** |
| --- | --- |
| contigs (>= 0 bp) | 19 |
| contigs (>= 1000 bp) | 19 |
| contigs (>= 5000 bp) | 16 |
| contigs (>= 10000 bp) | 15 |
| contigs (>= 25000 bp) | 15 |
| contigs (>= 50000 bp) | 15 |
| Total length (>= 0 bp) | 53,727,737 |
| Total length (>= 1000 bp) | 53,727,737 |
| Total length (>= 5000 bp) | 53,717,162 |
| Total length (>= 10000 bp) | 53,709,479 |
| Total length (>= 25000 bp) | 53,709,479 |
| Total length (>= 50000 bp) | 53,709,479 |
| contigs | 19 |
| Largest contig | 8,924,063 |
| Total length | 53,727,737 |
| GC (%) | 52 |
| N50 | 5,428,629 |
| N90 | 2,115,983 |
| L50 | 4 |
| L90 | 10 |
| N's per 100 kbp | 0 |

**Table S5. Gene Ontology classification of *A. pinea* F5 genome (top 50 GO terms).**

| **Hierarchy** | **GO-term** | **Count** |
| --- | --- | --- |
| BP | transmembrane transport | 476 |
| BP | phosphorylation | 310 |
| BP | methylation | 247 |
| BP | proteolysis | 242 |
| BP | regulation of transcription by RNA polymerase II | 234 |
| BP | carbohydrate metabolic process | 221 |
| BP | secondary metabolite biosynthetic process | 171 |
| BP | transcription, DNA-templated | 170 |
| BP | translation | 158 |
| BP | protein phosphorylation | 153 |
| BP | protein transport | 120 |
| BP | cell division | 103 |
| CC | membrane | 2,969 |
| CC | nucleus | 899 |
| CC | cytoplasm | 615 |
| CC | extracellular region | 248 |
| CC | mitochondrion | 242 |
| CC | cytosol | 219 |
| CC | ribosome | 213 |
| CC | ribonucleoprotein complex | 201 |
| CC | endoplasmic reticulum membrane | 136 |
| CC | nucleolus | 130 |
| CC | RNA polymerase I transcription factor complex | 120 |
| CC | mitochondrial inner membrane | 114 |
| CC | cell periphery | 107 |
| MF | ATP-binding | 706 |
| MF | hydrolase activity | 602 |
| MF | oxidoreductase activity | 599 |
| MF | metal ion binding | 584 |
| MF | DNA binding | 429 |
| MF | zinc ion binding | 399 |
| MF | transmembrane transport activity | 372 |
| MF | transferase activity | 297 |
| MF | RNA binding | 249 |
| MF | heme binding | 236 |
| MF | monooxygenase activity | 229 |
| MF | kinase activity | 225 |
| MF | RNA polymerase II transcription factor activity, sequence-specific DNA binding | 214 |
| MF | methyltransferase activity | 206 |
| MF | iron ion binding | 206 |
| MF | structural constituent of ribosome | 186 |
| MF | oxidoreductase activity, acting on paired donors, with incorporation or reduction of molecular oxygen | 184 |
| MF | nucleic acid binding | 163 |
| MF | FAD binding | 136 |
| MF | ATPase activity | 135 |
| MF | GTP-binding | 134 |
| MF | ligase activity | 116 |
| MF | flavin adenine dinucleotide binding | 110 |
| MF | protein kinase activity | 110 |
| MF | GTPase activity | 104 |

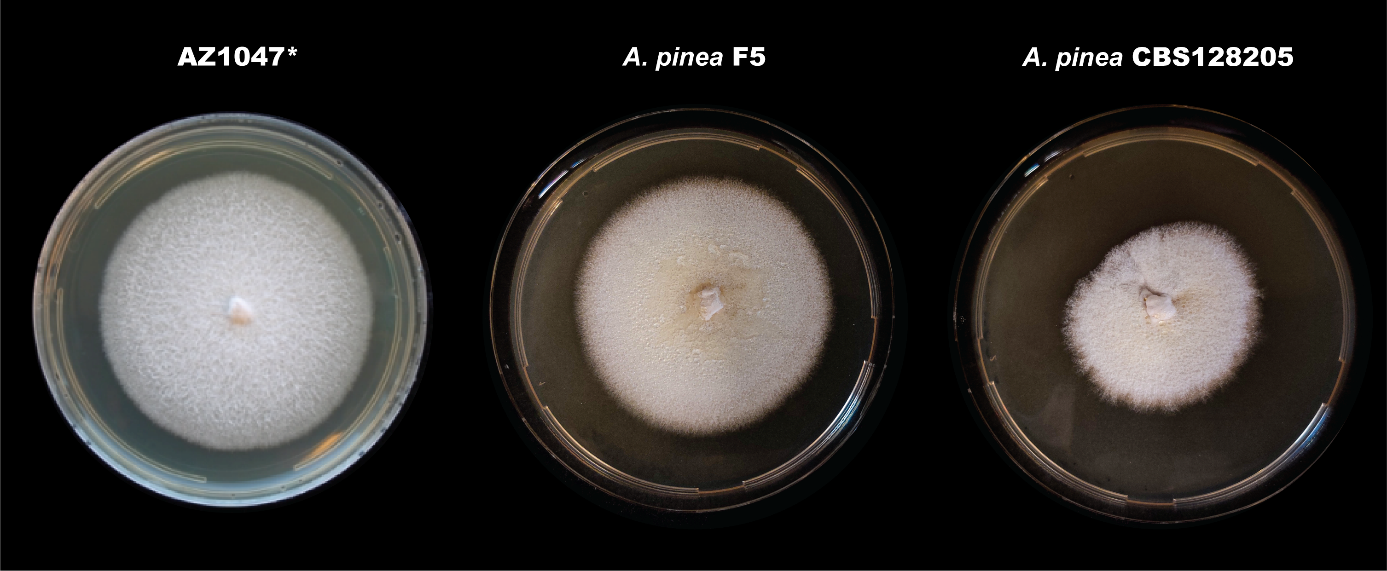
**Supplementary figures**

**Figure S1.** Morphological comparison of isolate AZ1047 (picture kindly provided by U’ren et al. [1]), A. pinea F5 isolated in our lab, and the culture collection strain A. pinea CBS128205.

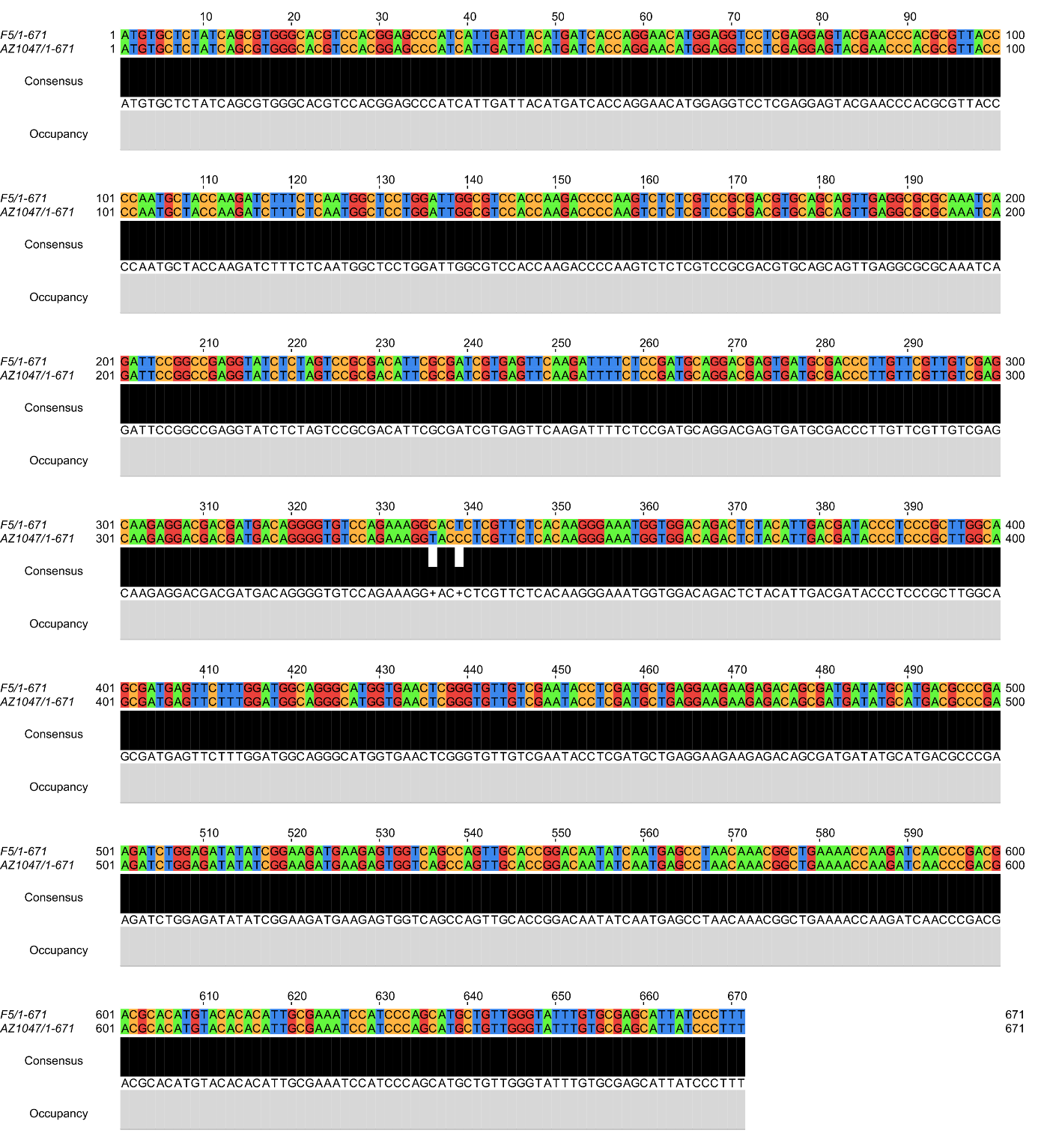

**Figure S2. Alignment analysis between the sequences of the rpb2 genes of A. pinea F5 and AZ1047.** The sequences are nearly identical, with only two mismatches observed at positions 336 and 339, suggesting that the two isolates likely belong to the same species.

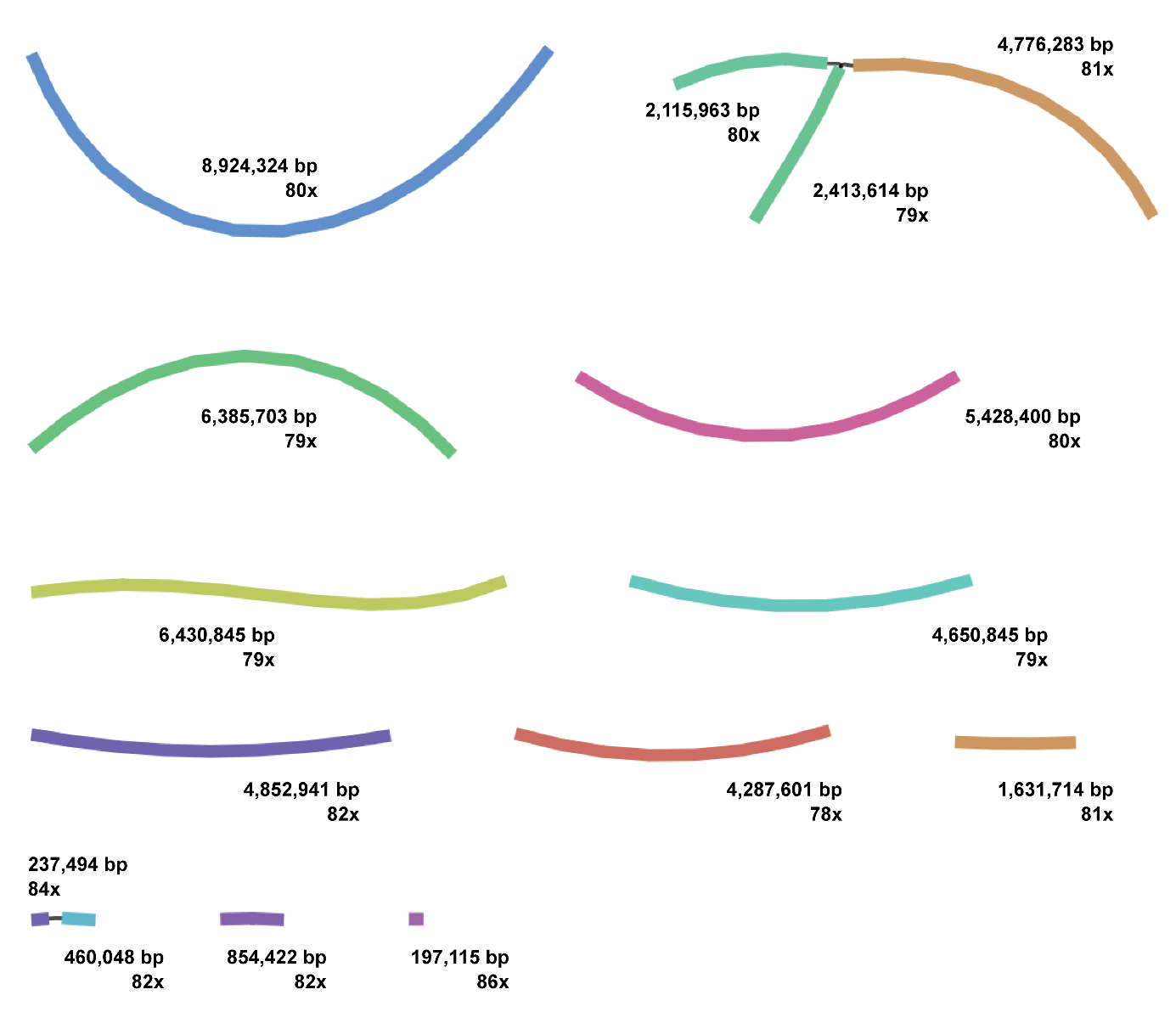

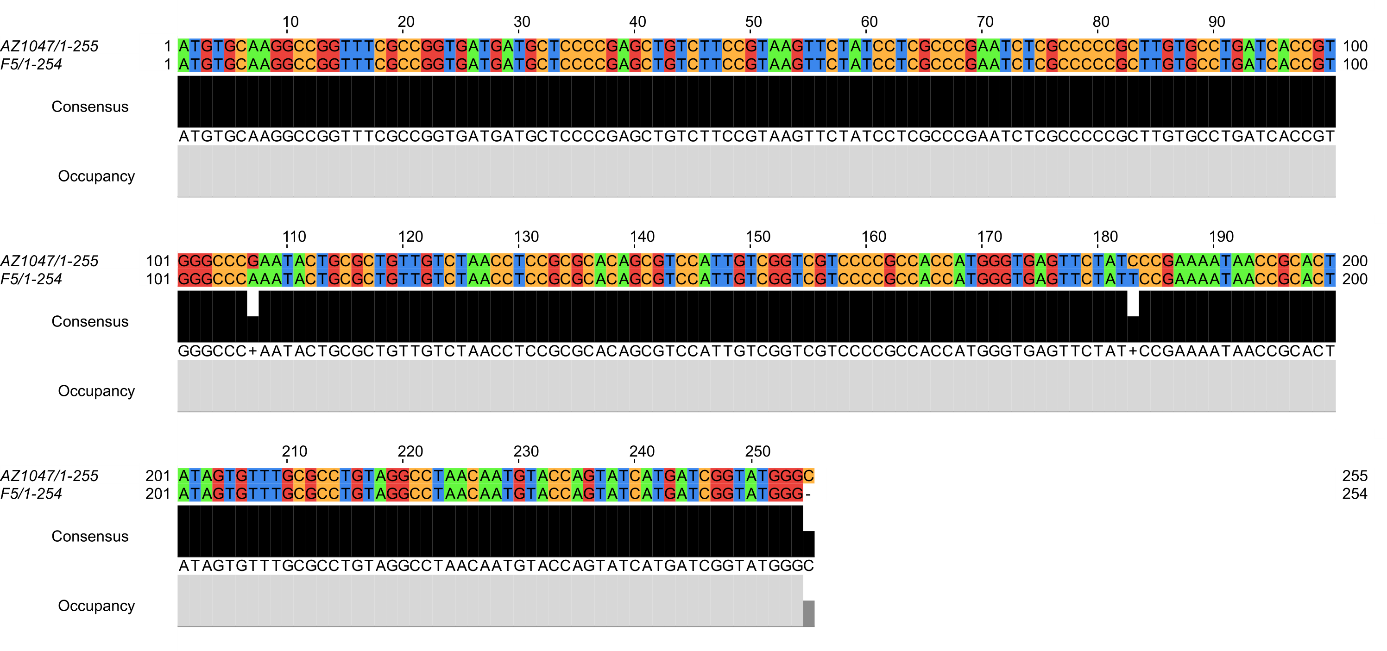

**Figure S5. Contigs of A. pinea F5 genome assembly visualized by Bandage [2].**

**Figure S3. Alignment analysis between the sequences of the α-actin genes of A. pinea F5 and AZ1047.** The sequences are nearly identical, with only two mismatches observed at positions 107 and 183 and one gap at position 255, suggesting that the two isolates likely belong to the same species.

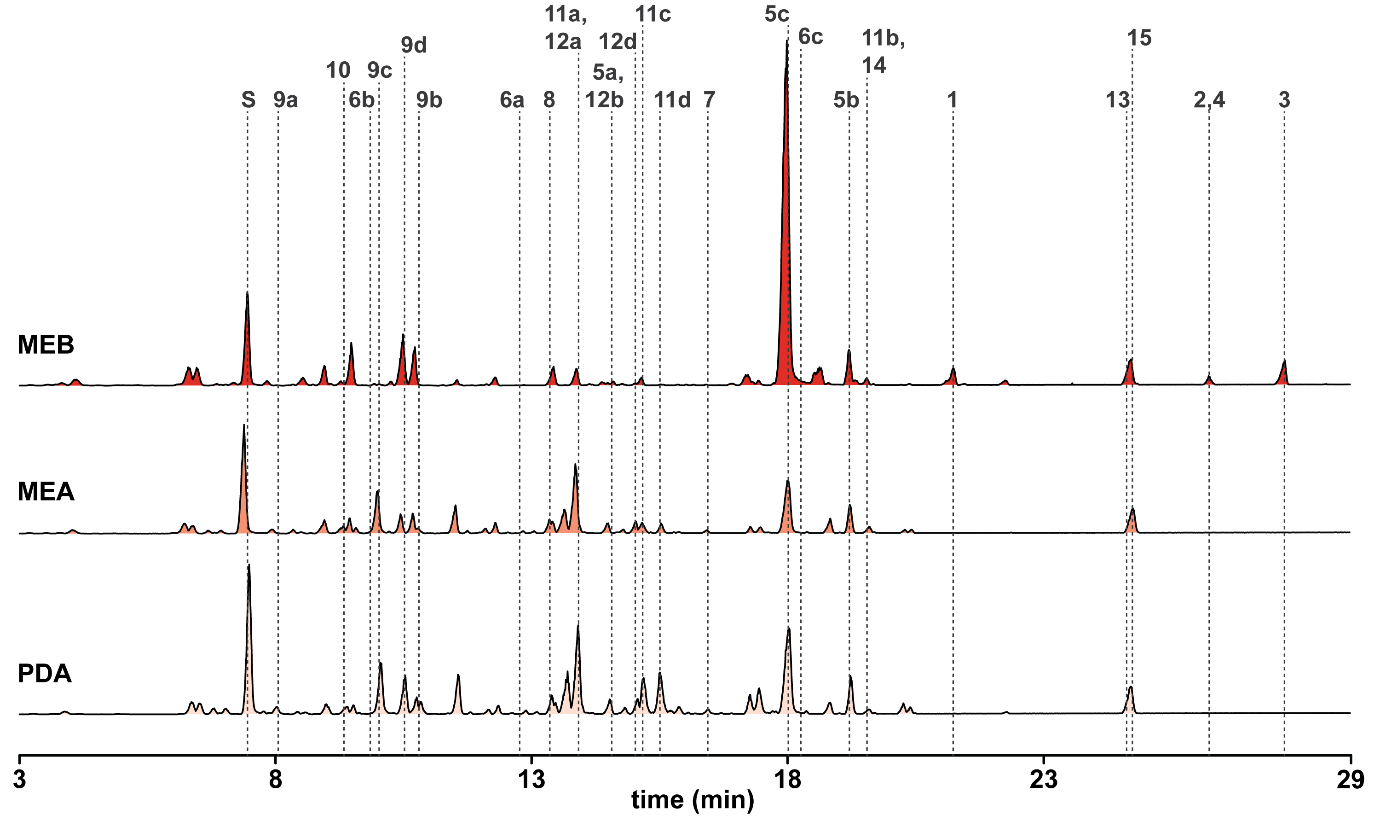

**Figure S5. Base peak intensity (BPI) LC-MS chromatograms of extracts from A. pinea F5 cultures.** The annotated nodes from the molecular network (Fig. 5A and Table 3) are marked by dashed lines.

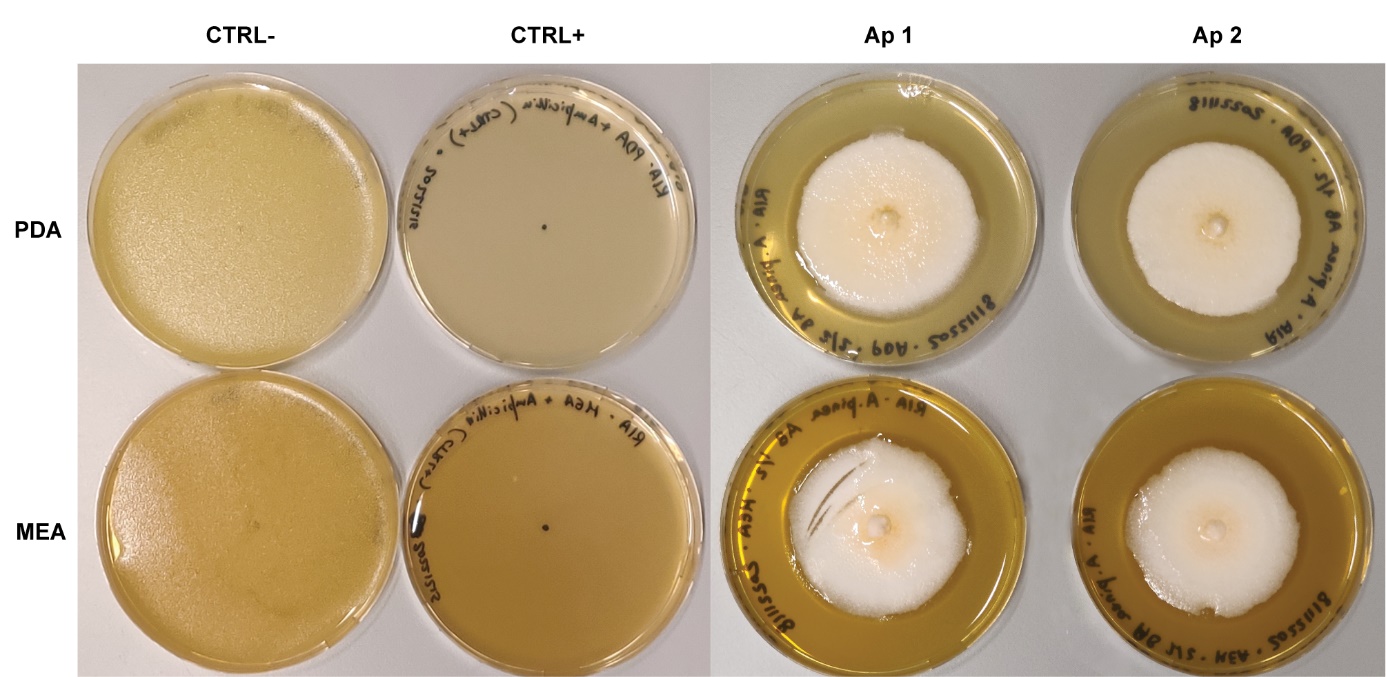

**Figure S6. Antibacterial assays of whole colonies of A. pinea F5 (in duplicates).** For the positive control, ampicillin 100 µg/mL was added to the media. A. pinea F5 completely inhibited the growth of the indicator strain M. luteus 10240.
