## Additional file 7 for "Genome sequencing and molecular networking analysis of the wild fungus *Anthostomella pinea* reveal its ability to produce a diverse range of secondary metabolites"

### GNPS hit 1 (MEB) - XIC range m/z 233.152-233.154 [M+H]

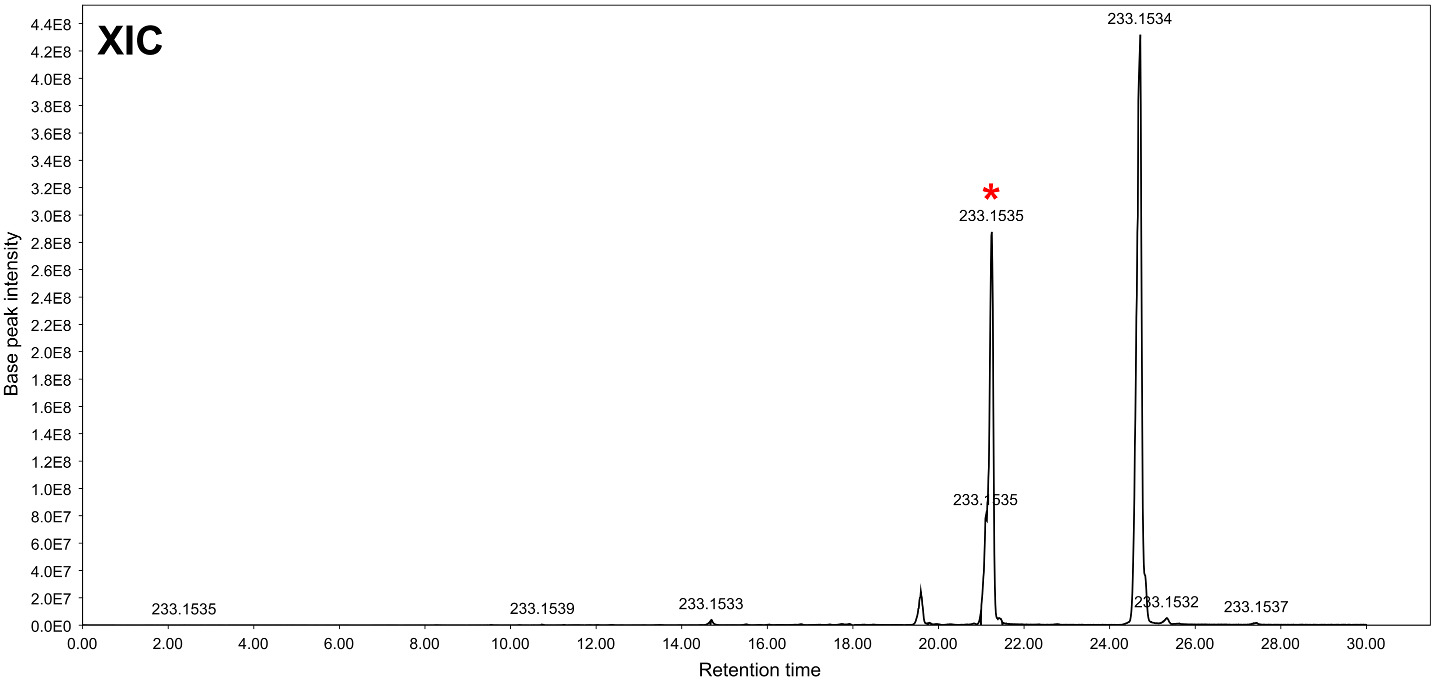


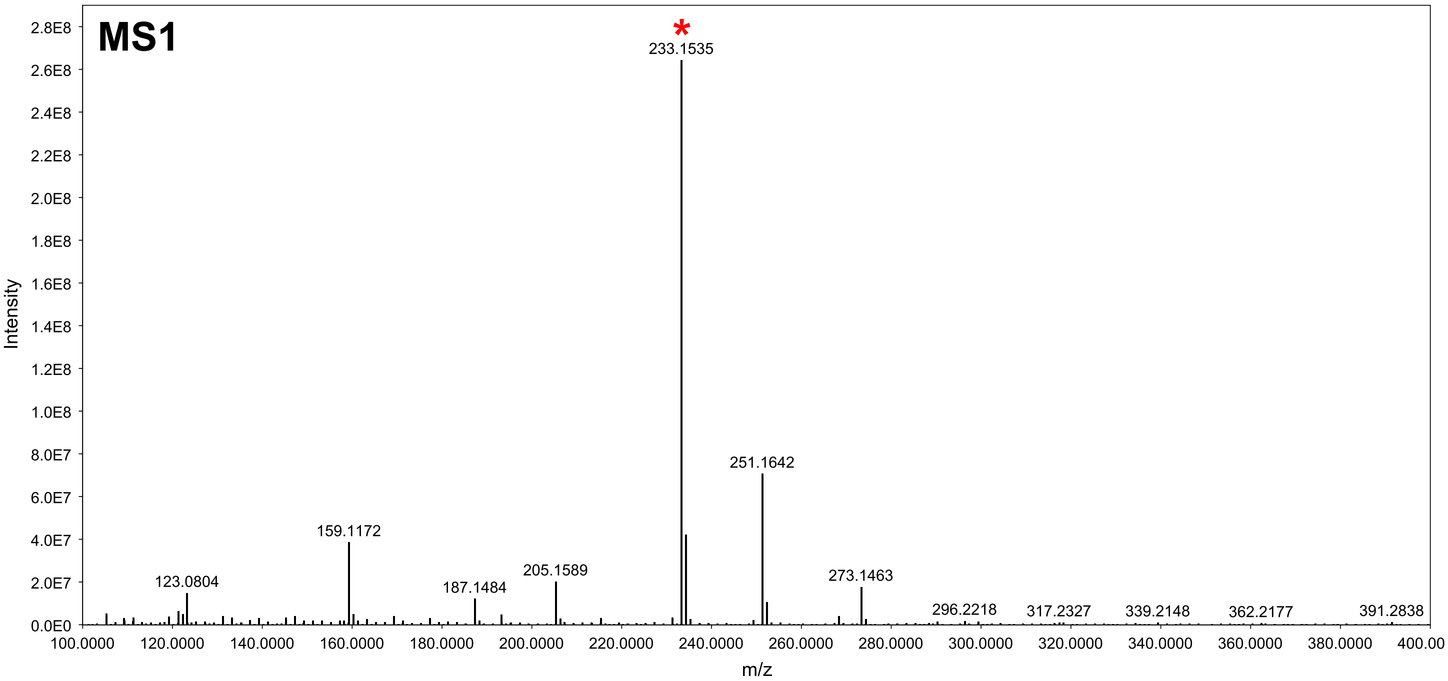


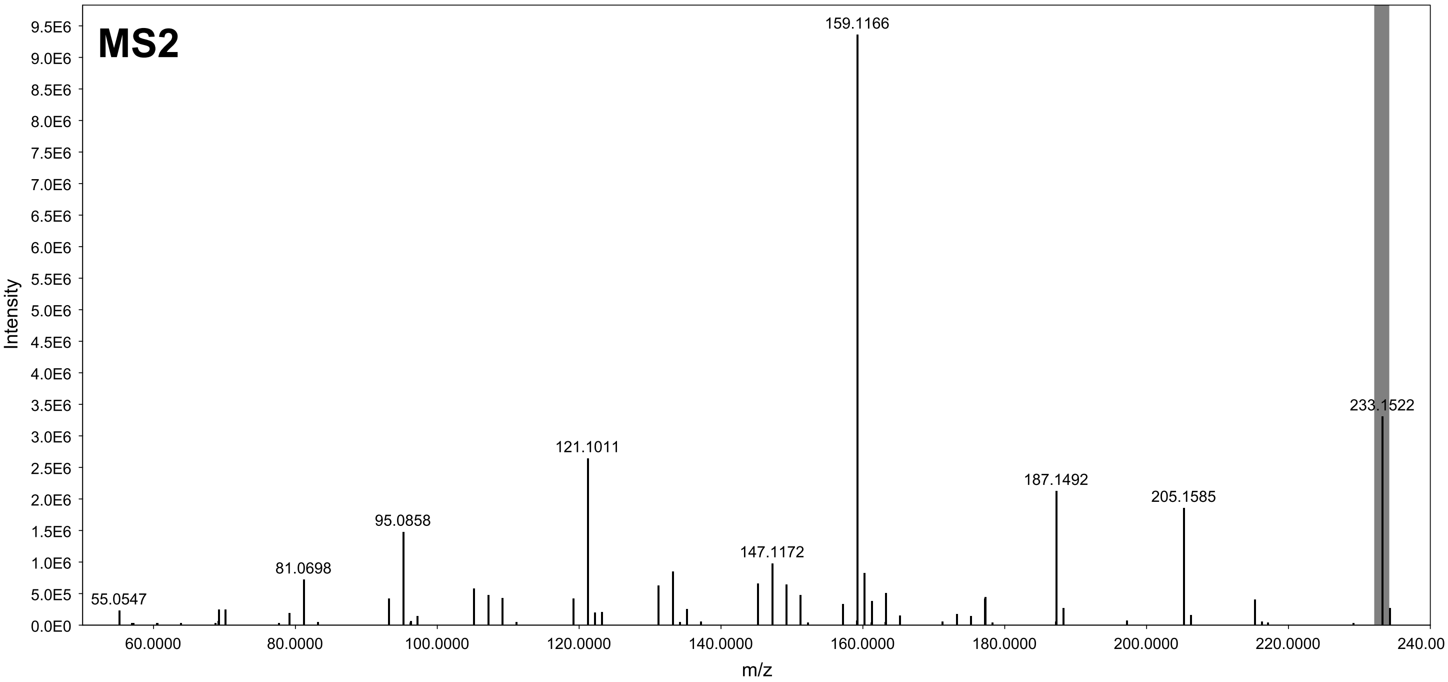

### GNPS hit 2 (MEB) - XIC range m/z 251.199-251.201 [M+H]

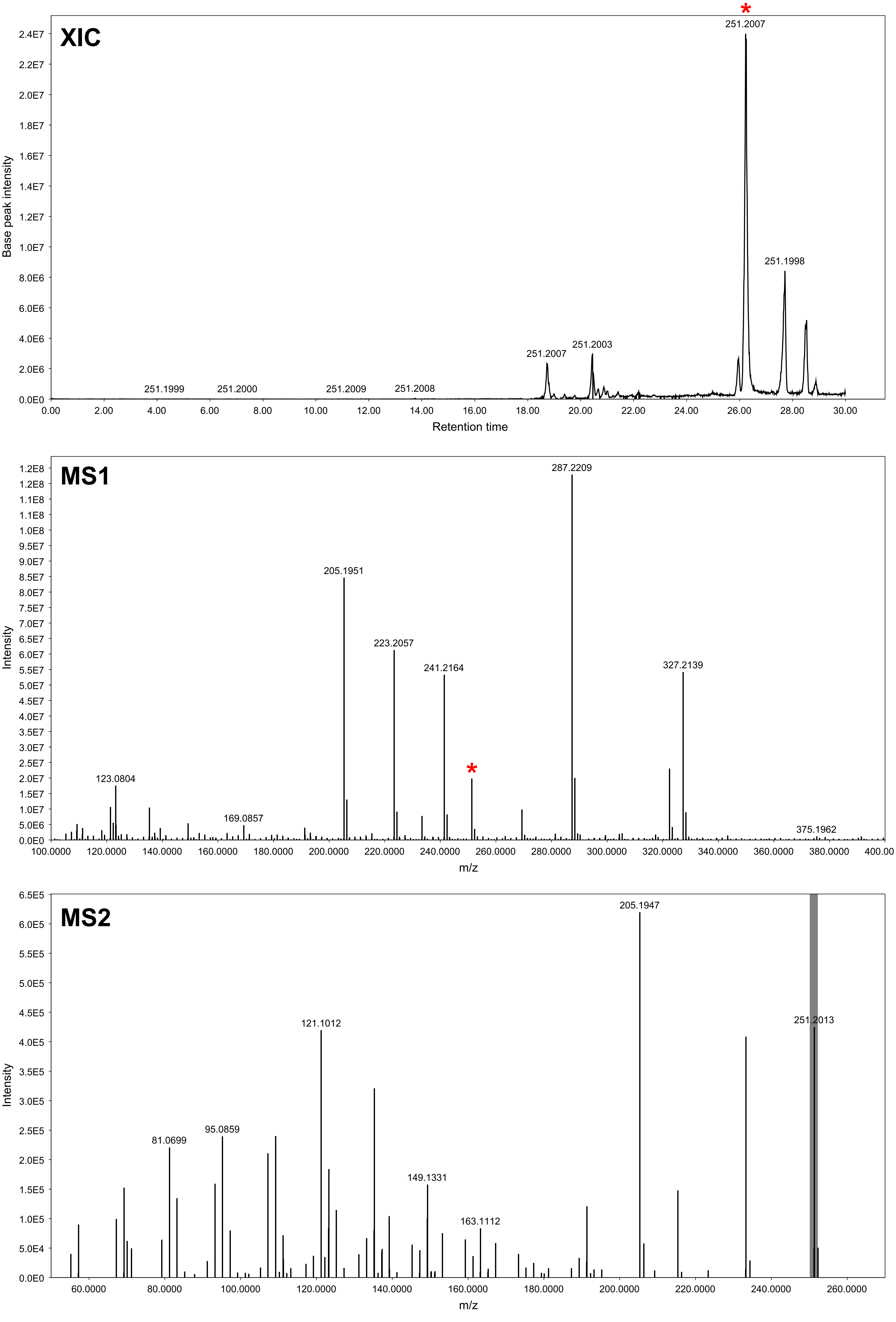

### GNPS hit 3 (MEB) - XIC range m/z 223.204-22.206 [M+H]

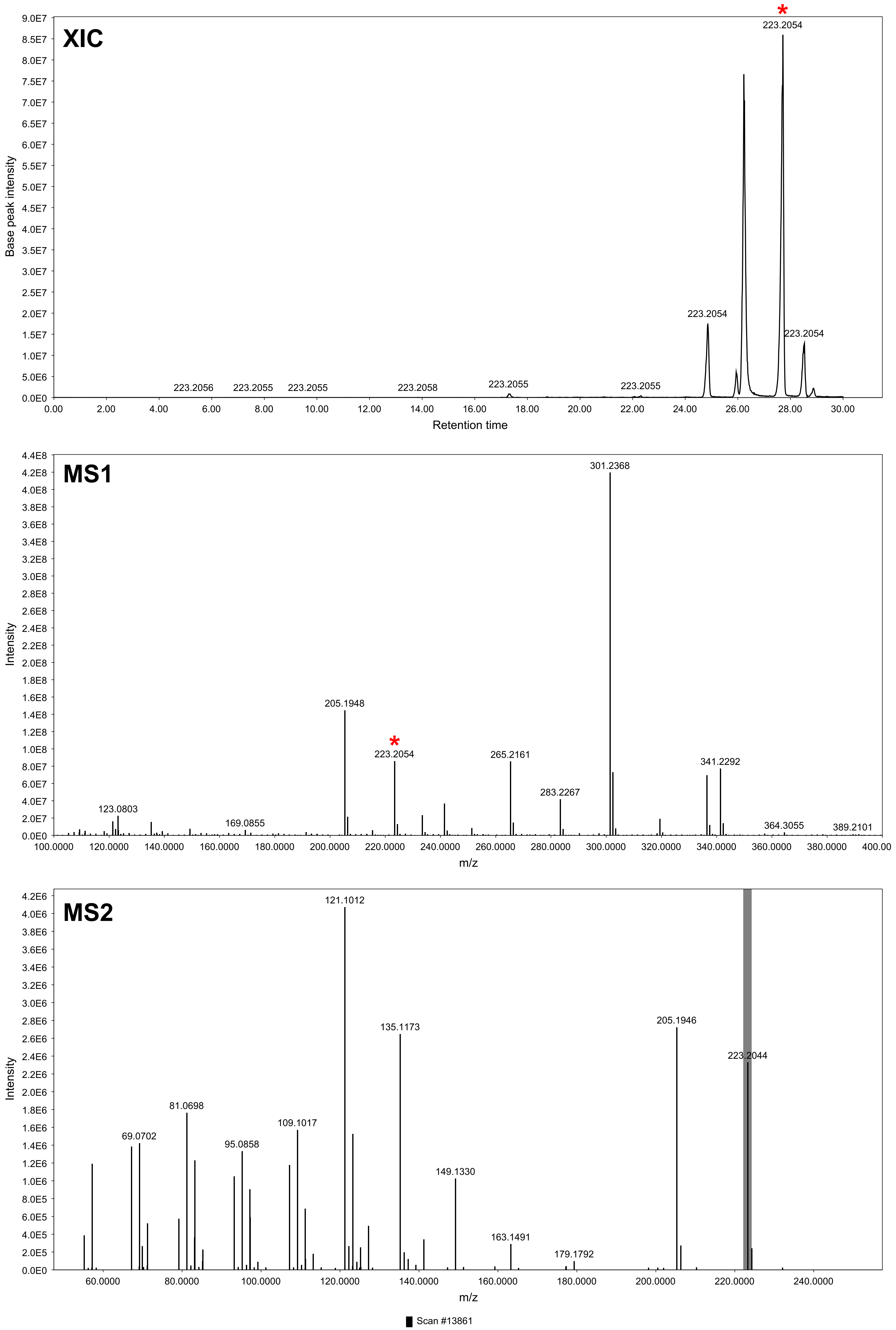

### GNPS hit 4 (MEB) - XIC range m/z 205.194-205.196 [M-H2O+H] [M+H]

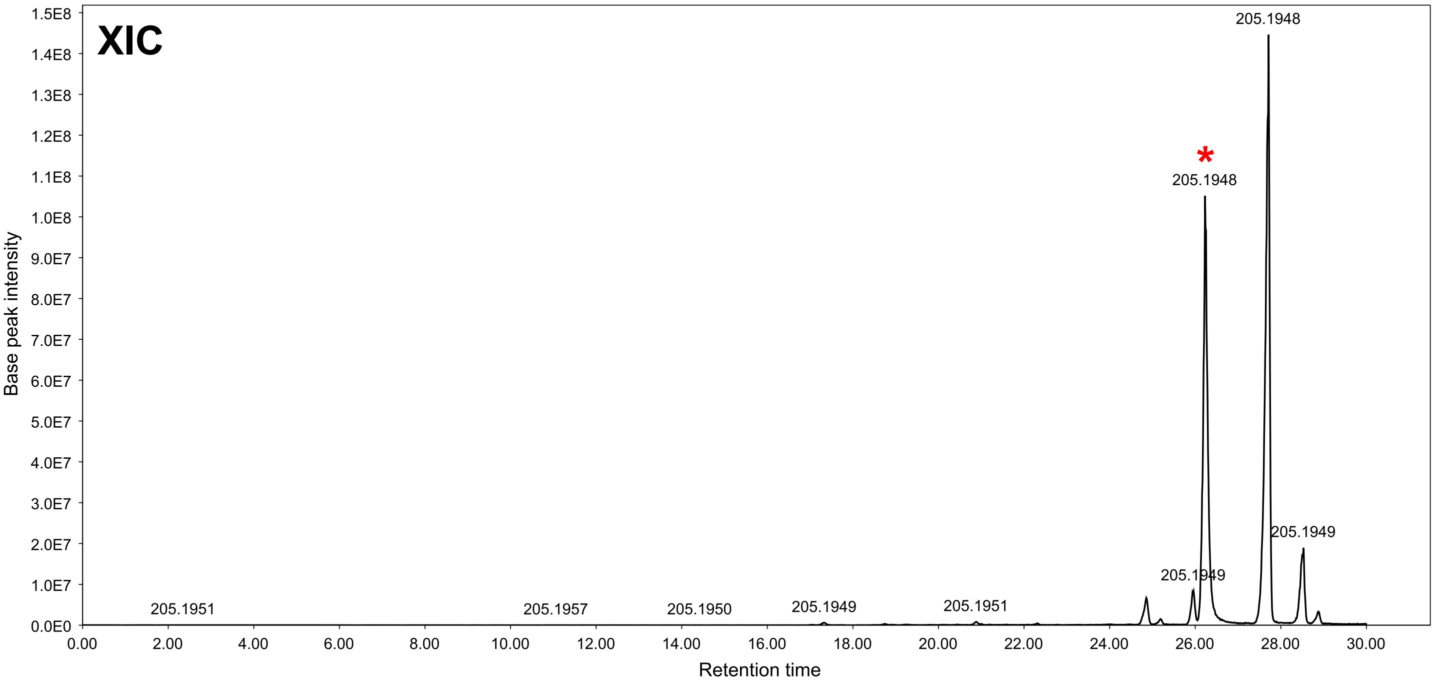


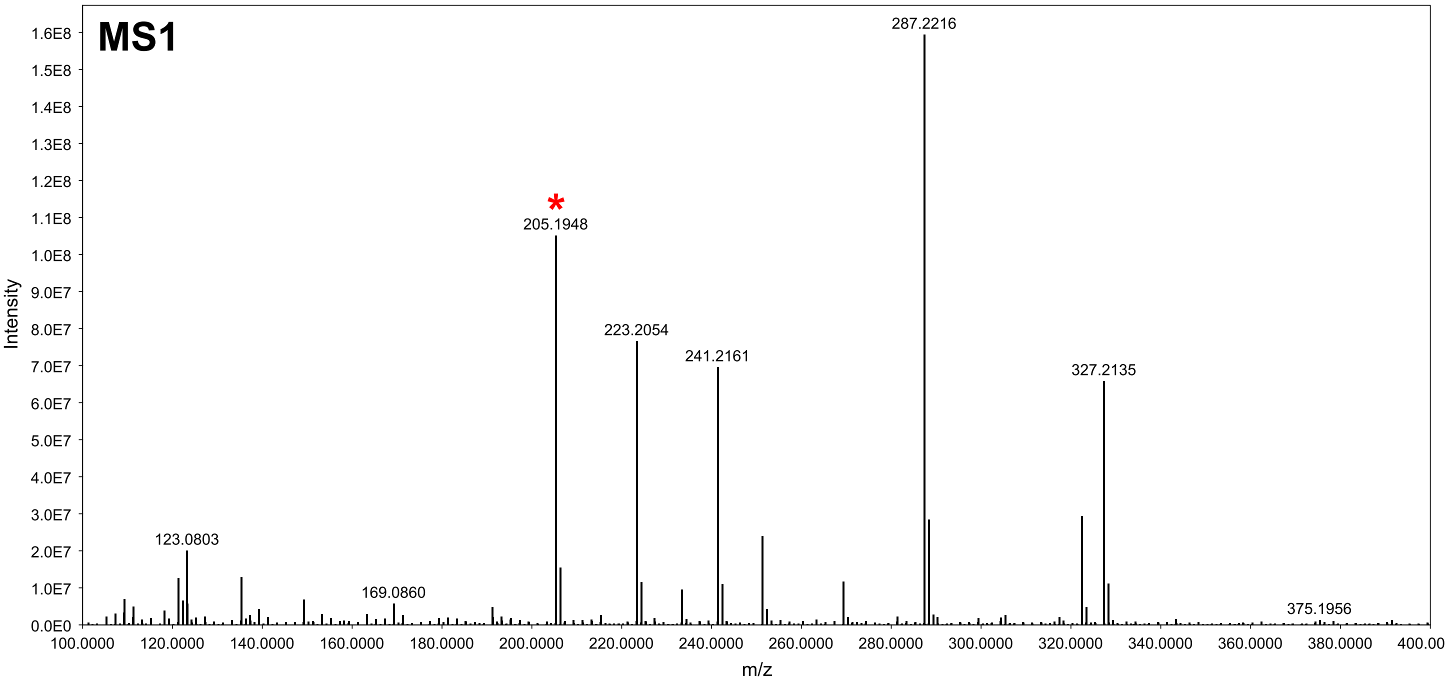


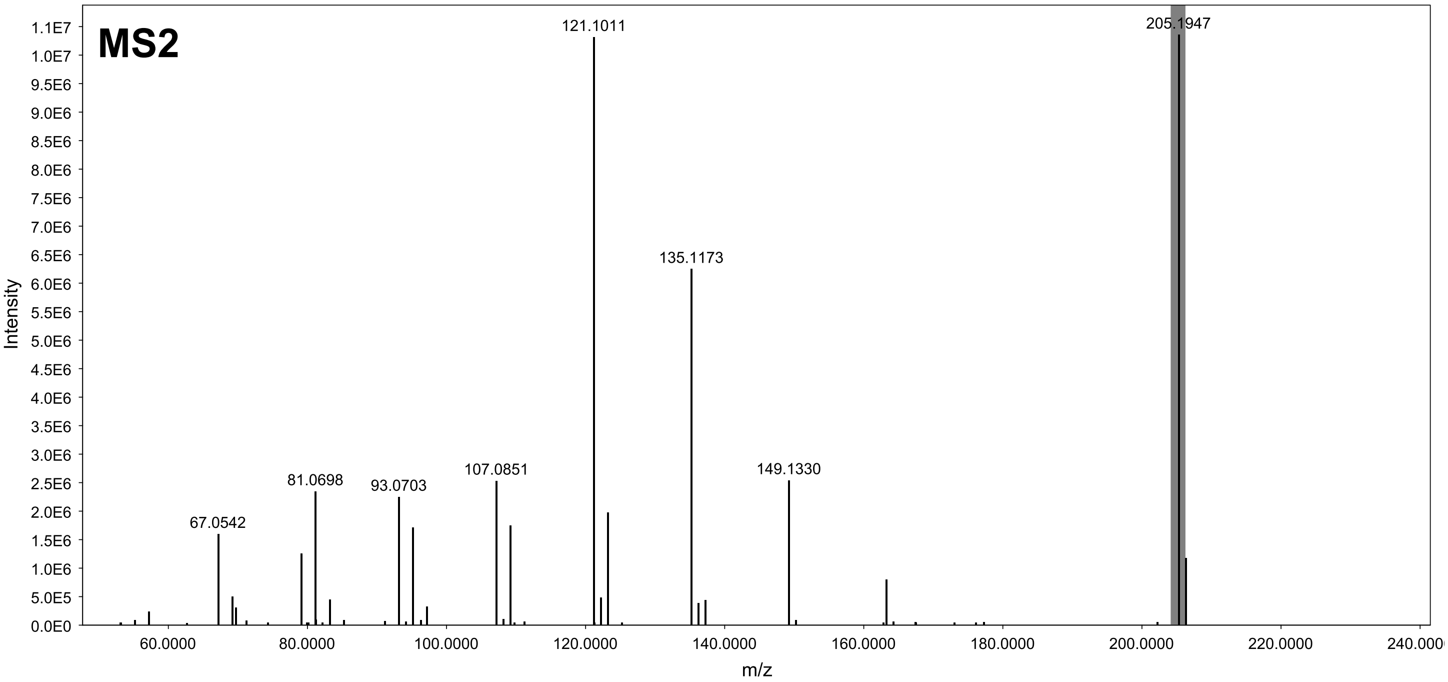

### GNPS hit 5a (MEB) - XIC range m/z 203.179-203.181 [M-H2O+H]

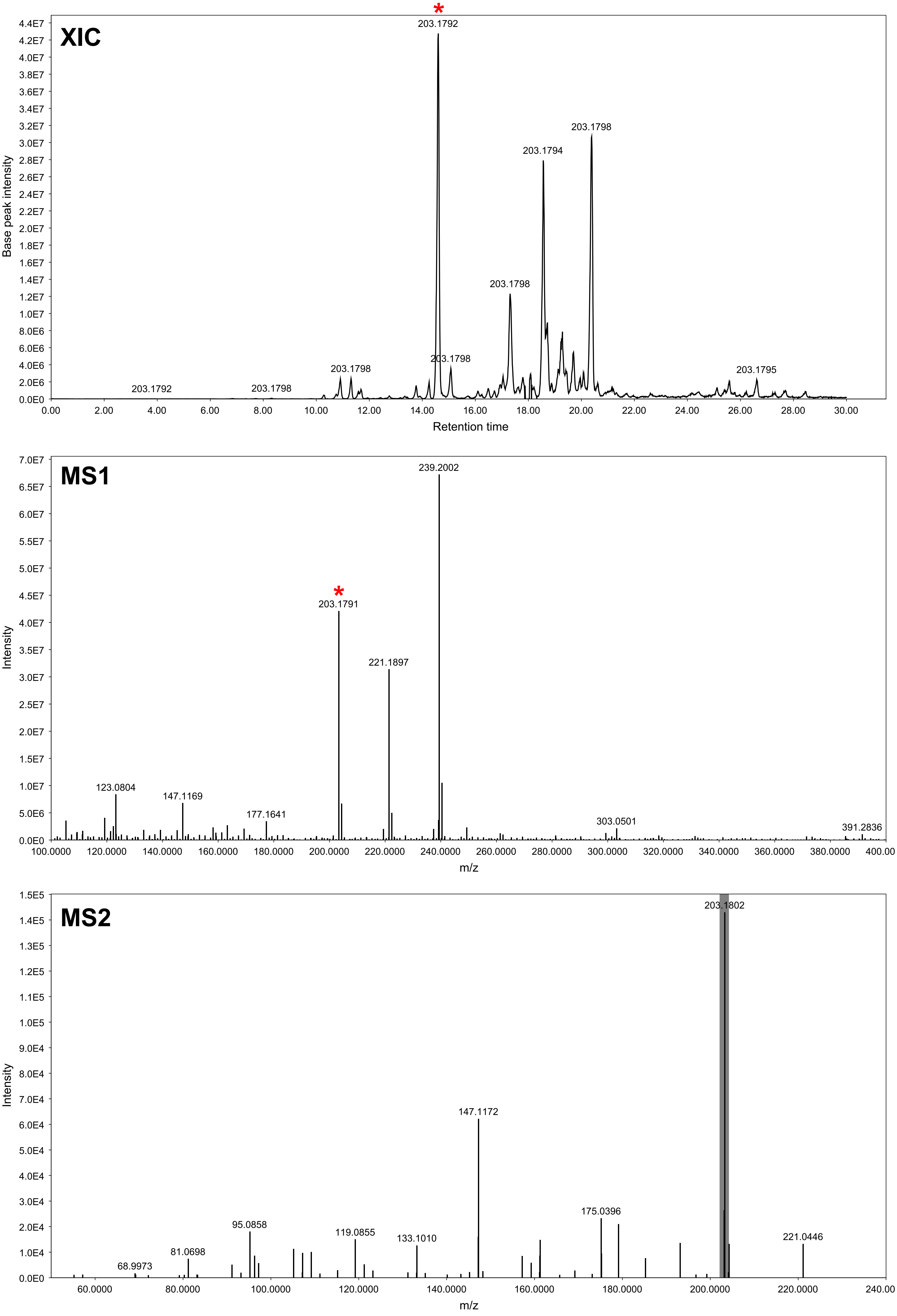

### GNPS hit 5b (MEB) - XIC range m/z 203.179-203.181 [M-H2O+H]

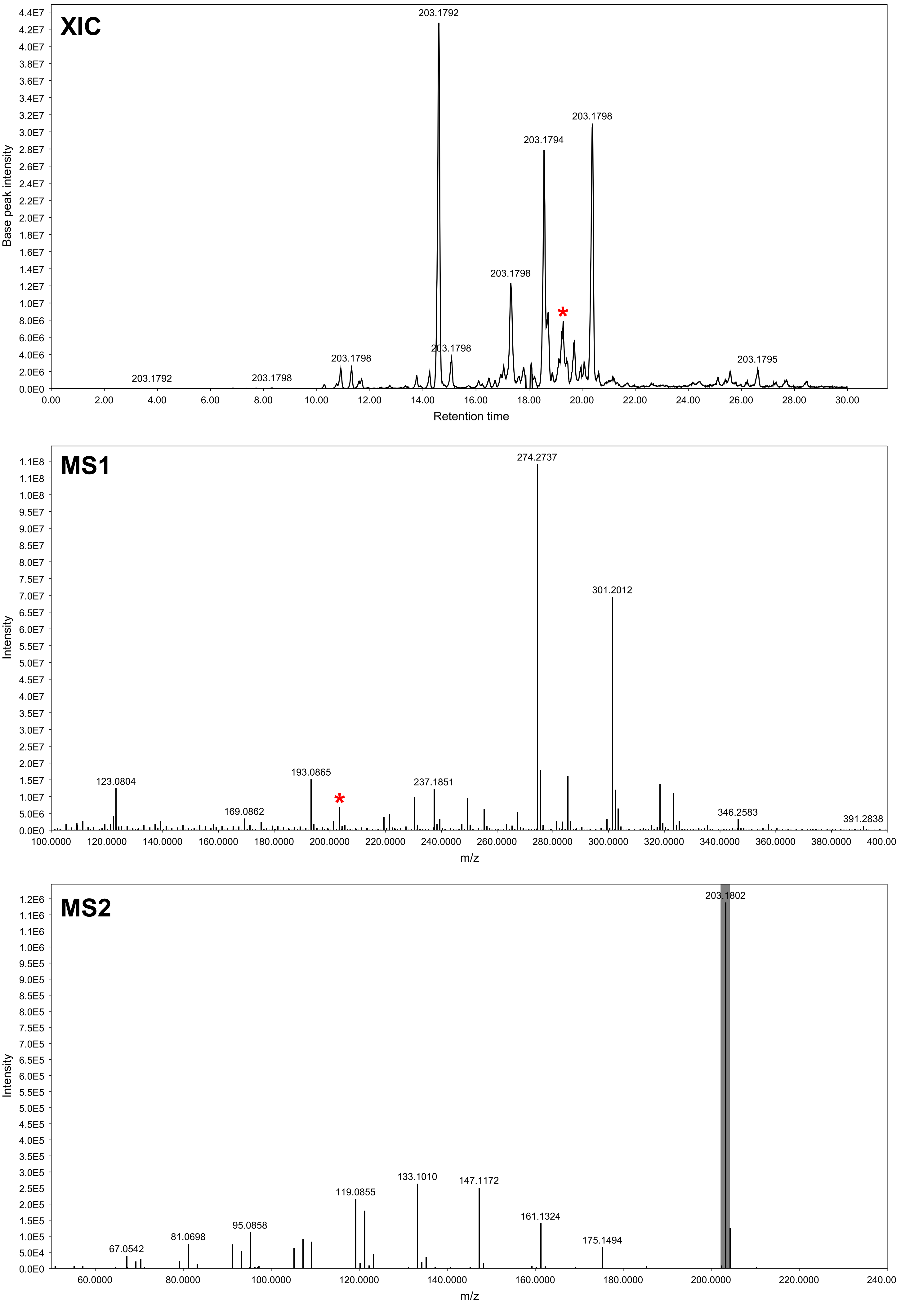

### GNPS hit 5c (MEB) - XIC range m/z 203.179-203.181 [M-H2O+H]

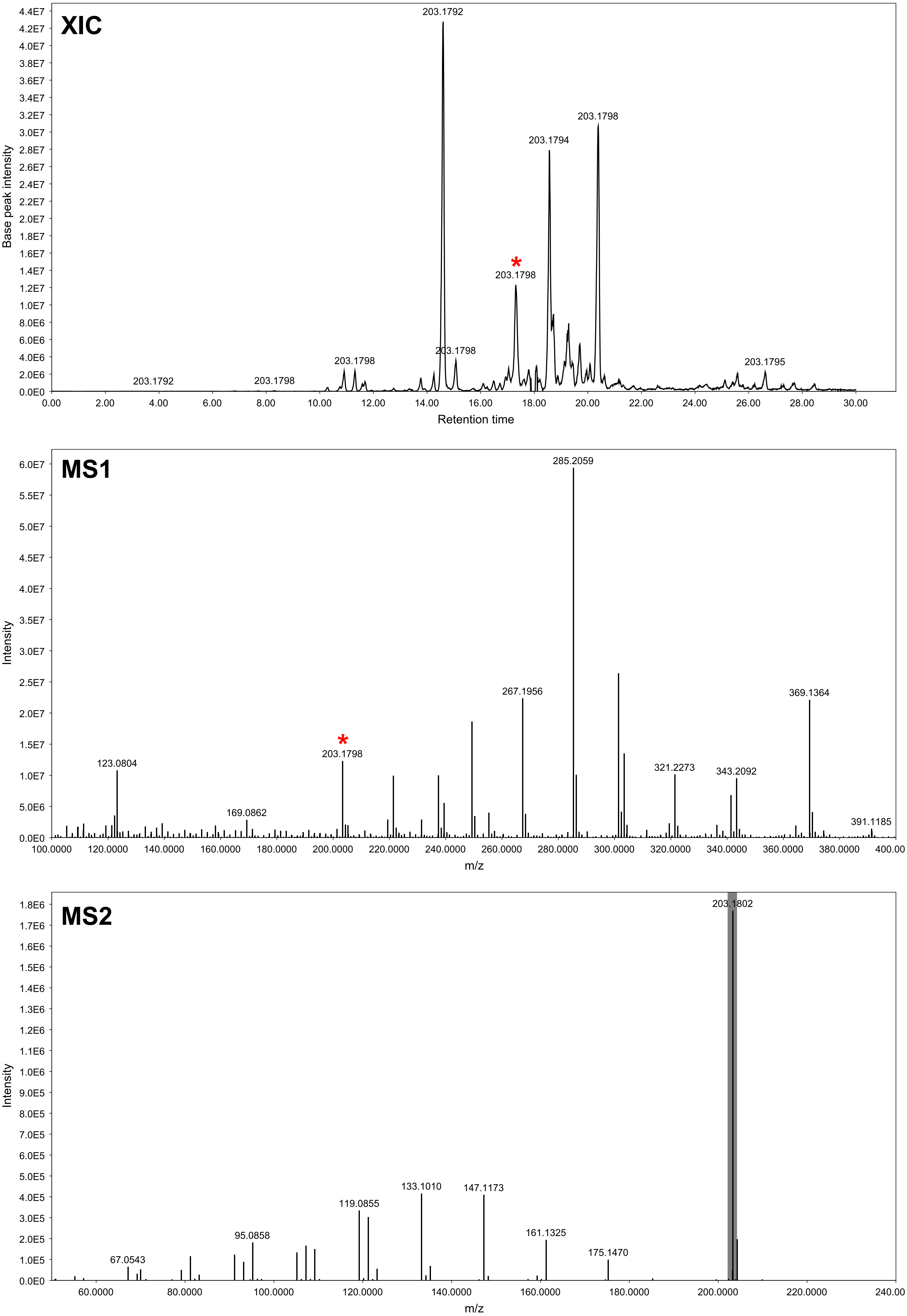

### GNPS hit 6a (PDA) - XIC range m/z 201.163-201.165 [M-2$H_2$O+H]

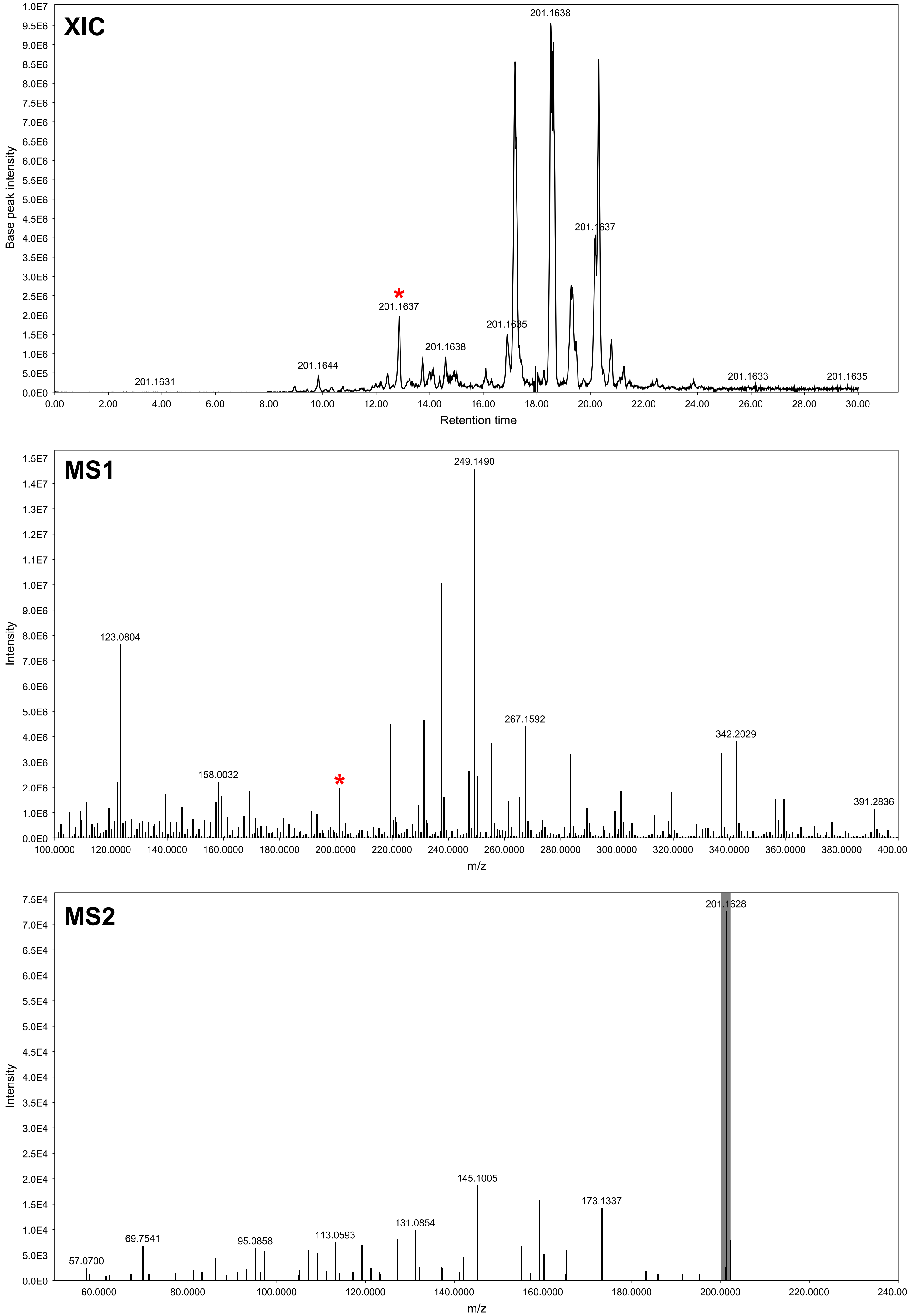

### GNPS hit 6b (PDA) - XIC range m/z 219.172-219.174 [M-$H_2O$+H]

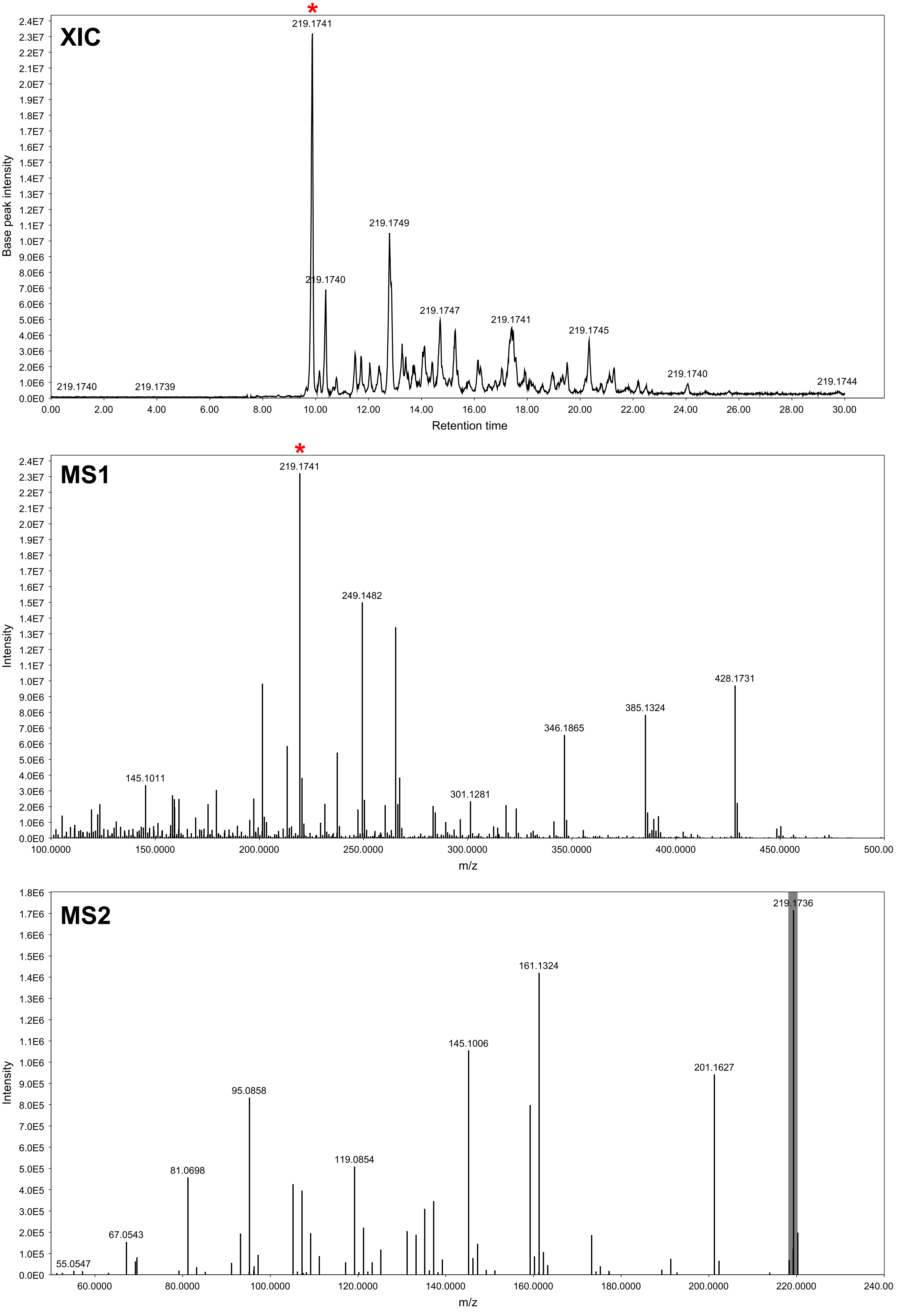

### GNPS hit 6c (MEB) - XIC range m/z 237.184-237.186 [M+H]

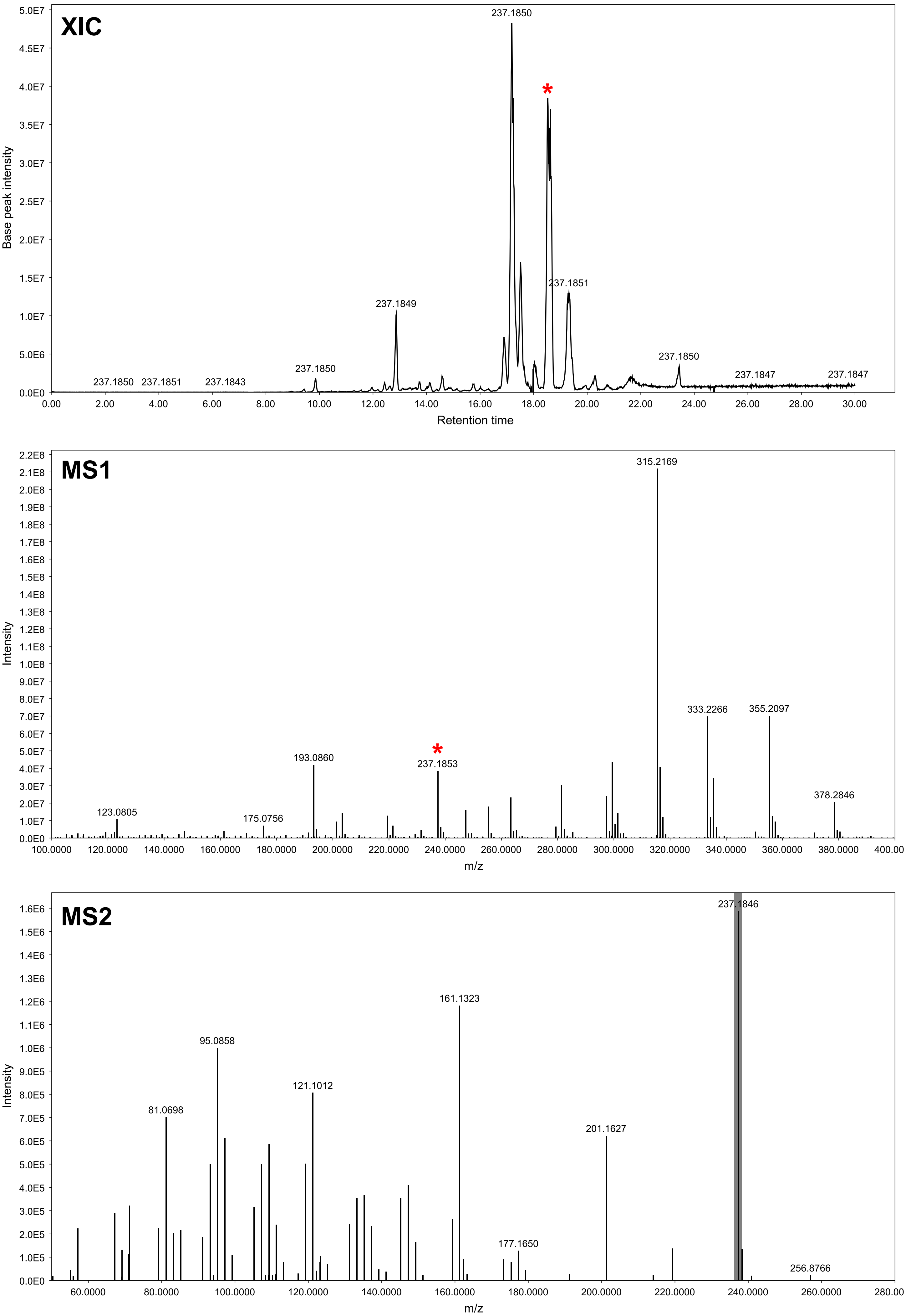

### GNPS hit 7 (MEB) - XIC range m/z 235.168-235.170 [M+H]

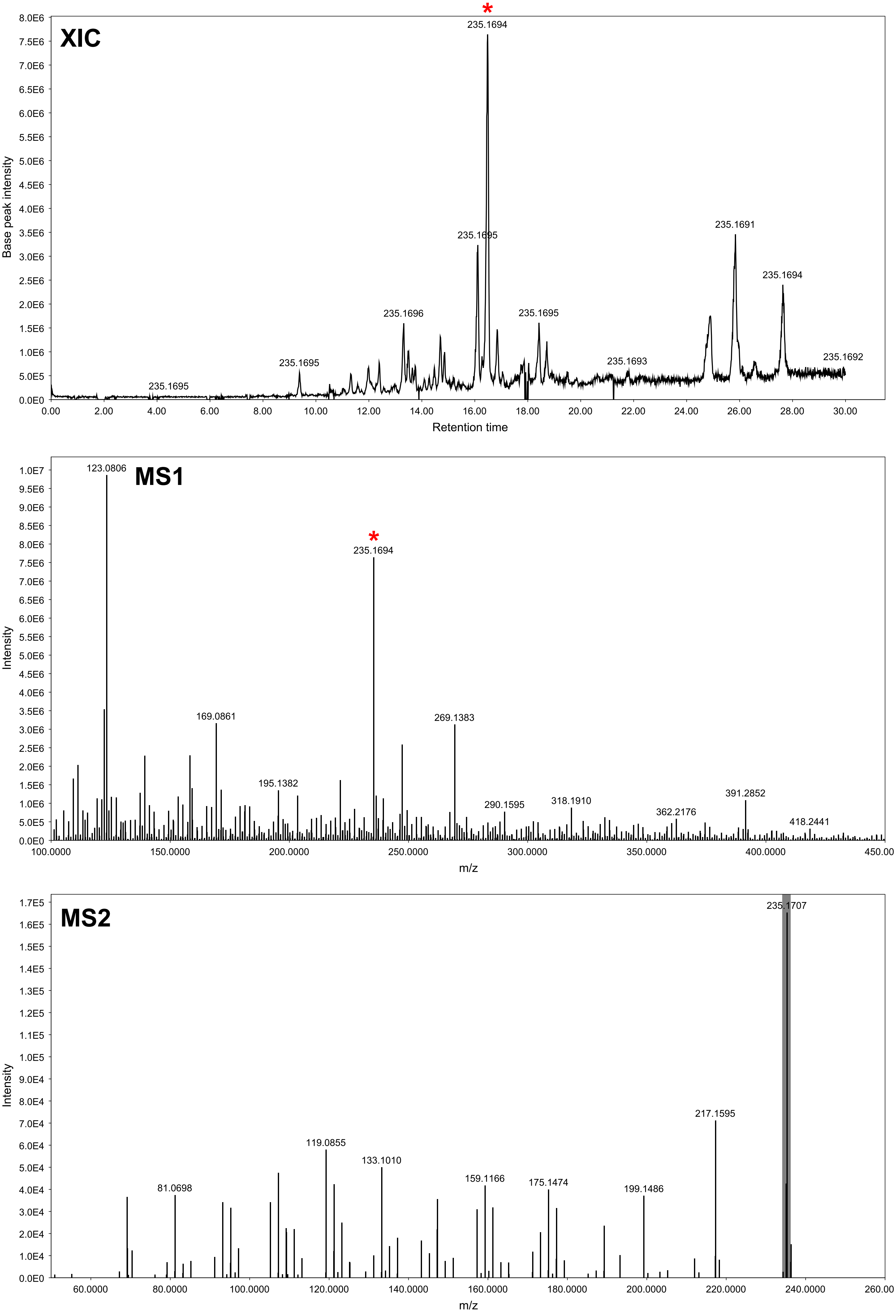

### GNPS hit 8 (MEB) - XIC range m/z 235.168-235.170

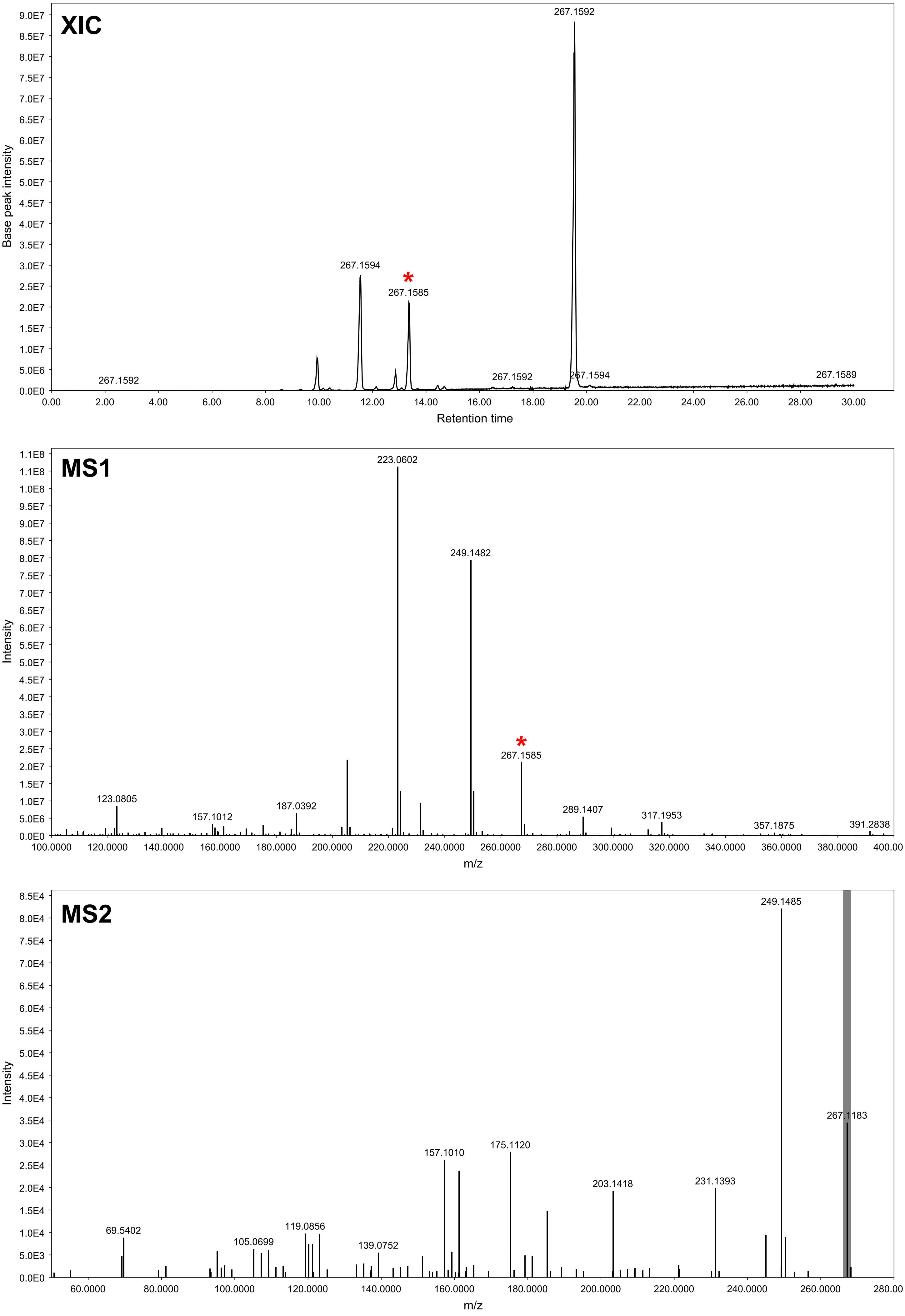

### GNPS hit 9a (PDA) - XIC range m/z 265.142-265.144 [M+H]

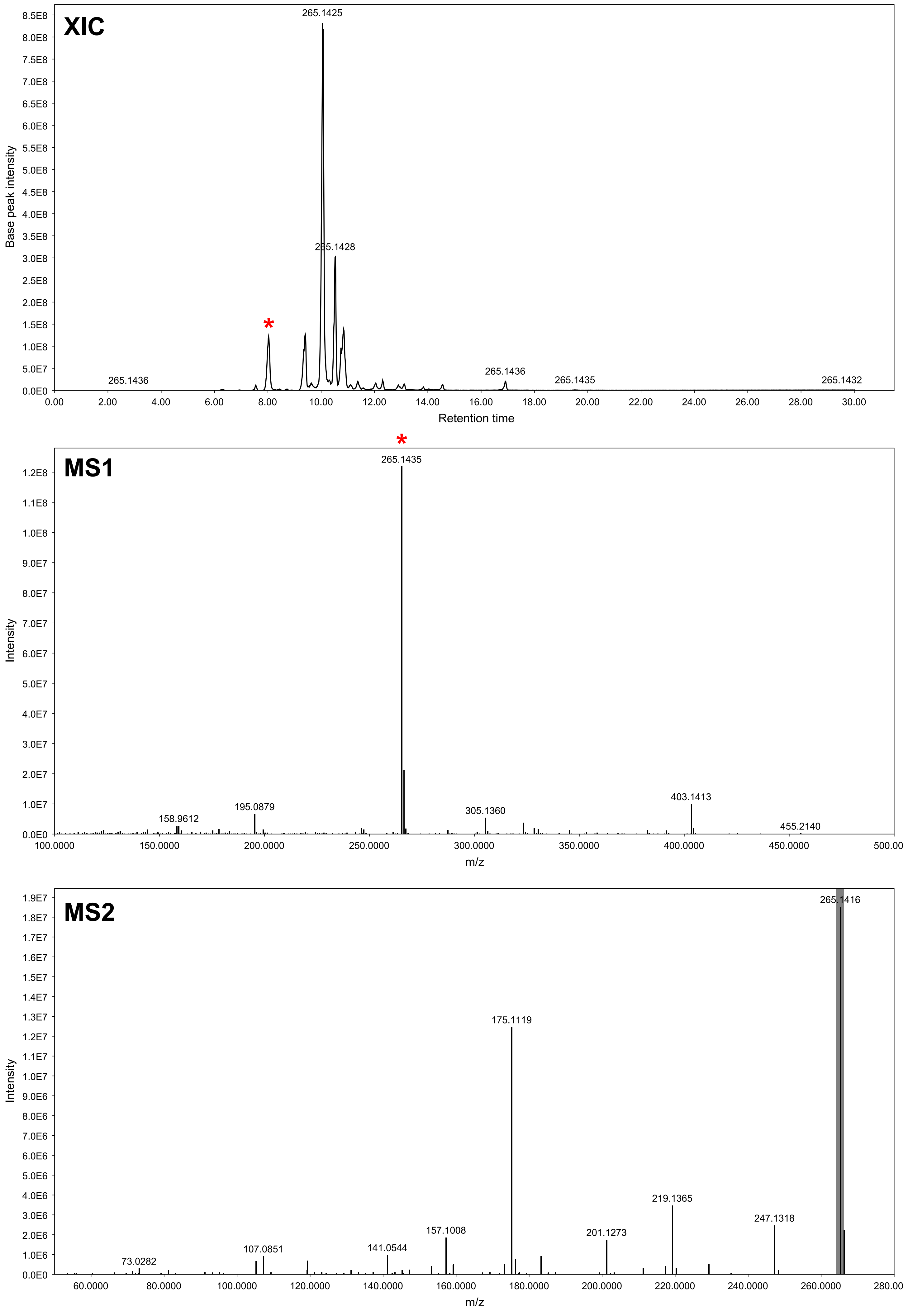

### GNPS hit 9b (PDA) - XIC range m/z 265.142-265.144 [M+H]

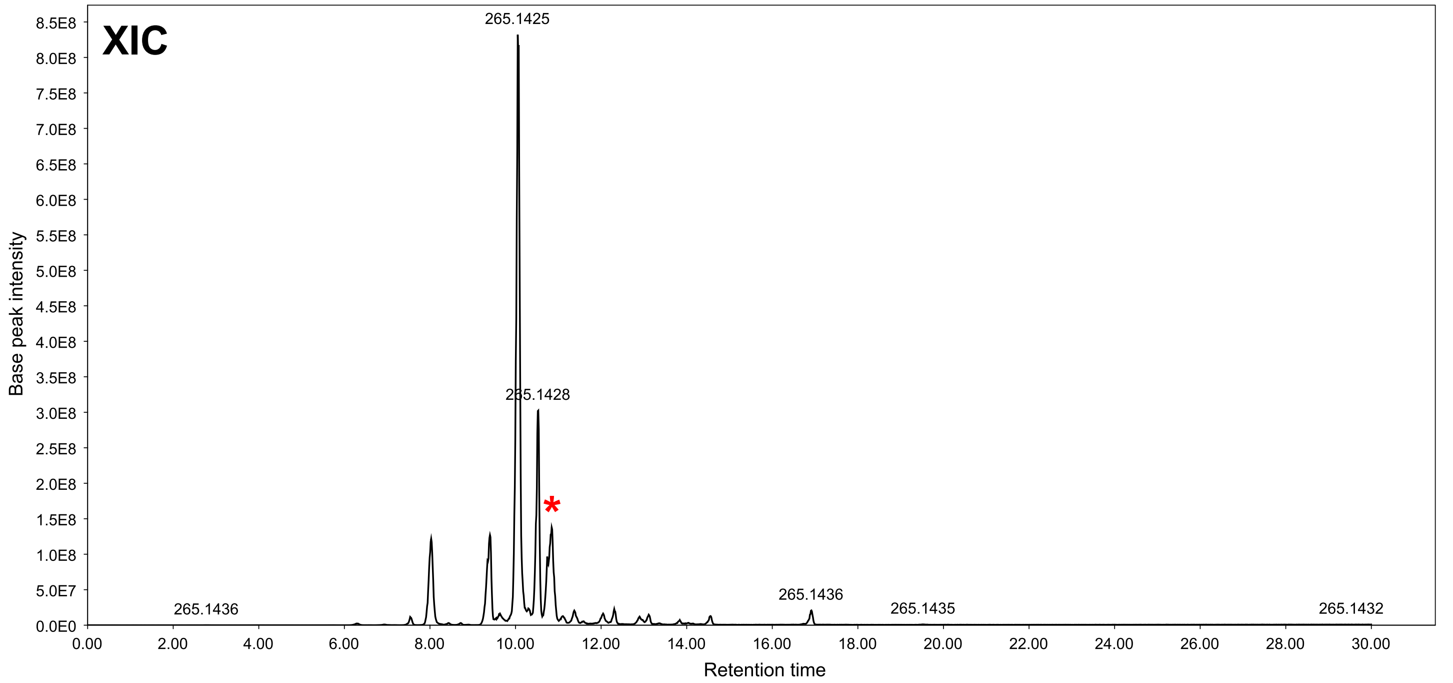


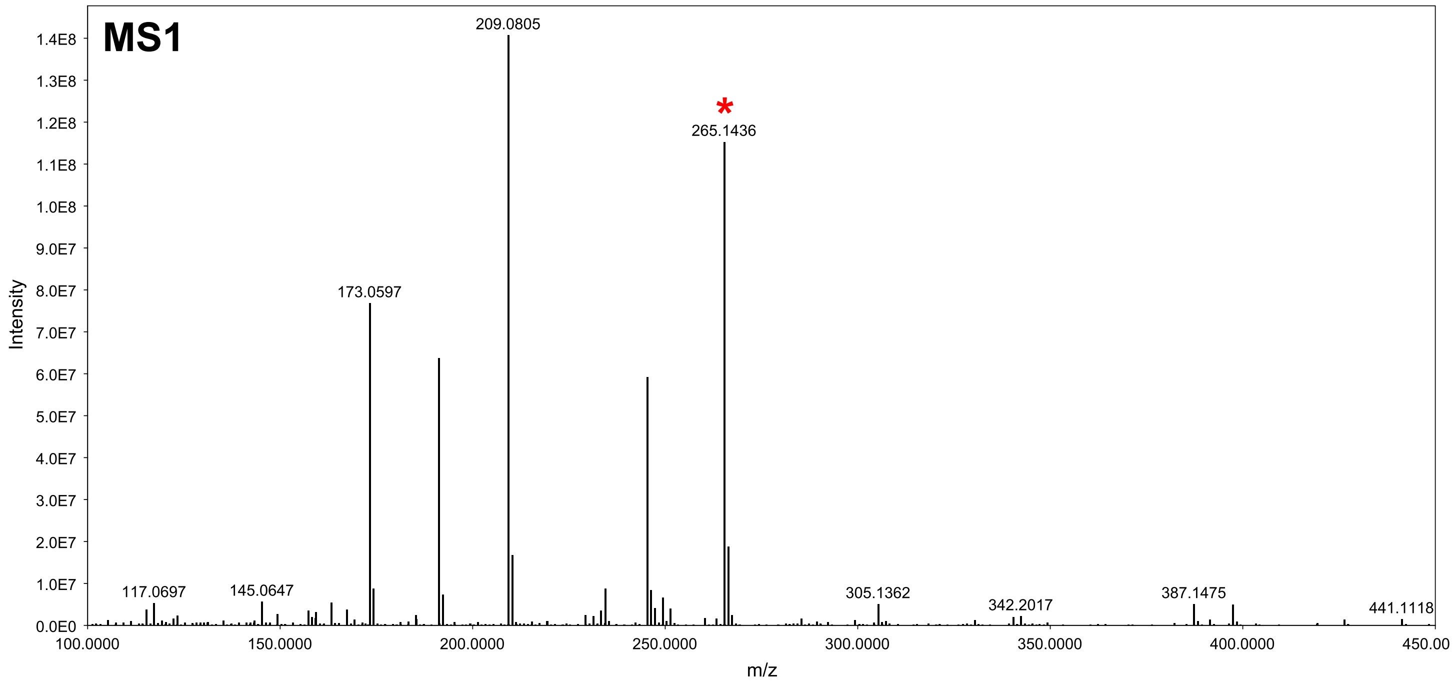


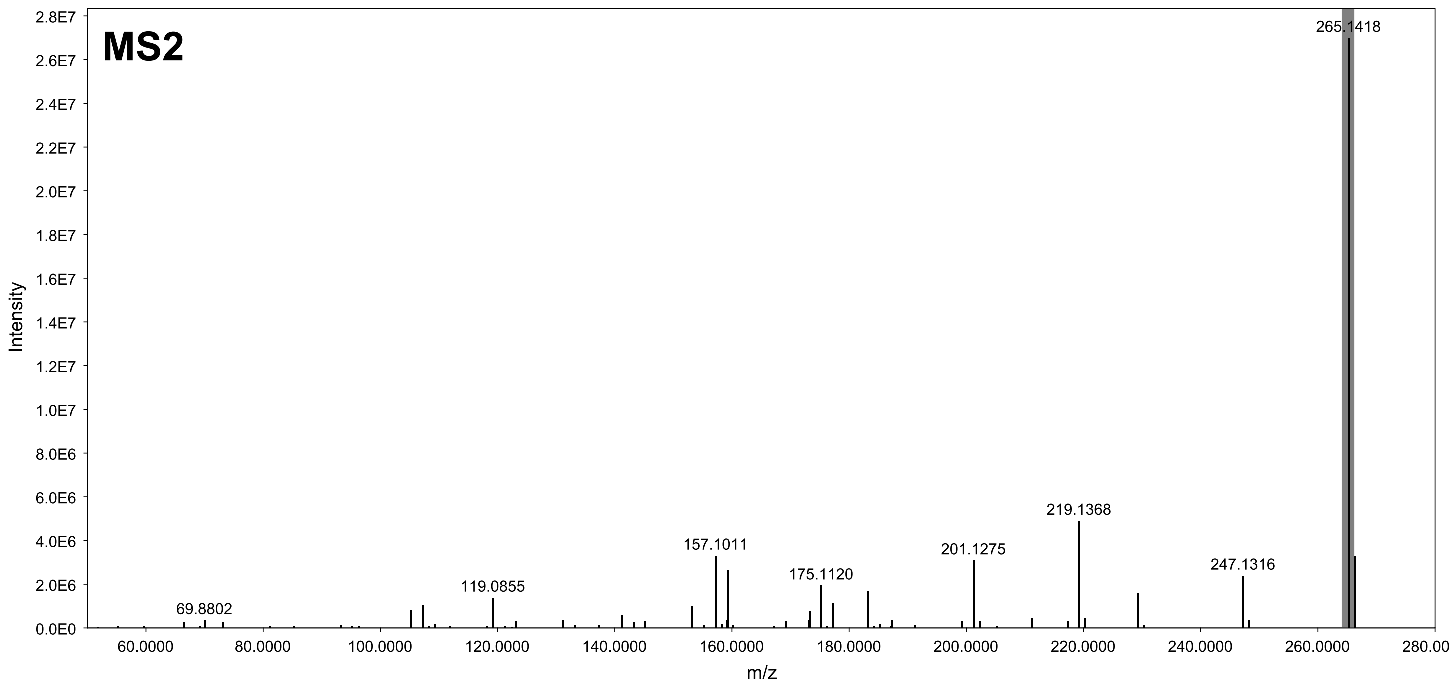

### GNPS hit 9c (MEB) - XIC range m/z 265.142-265.144 [M+H]

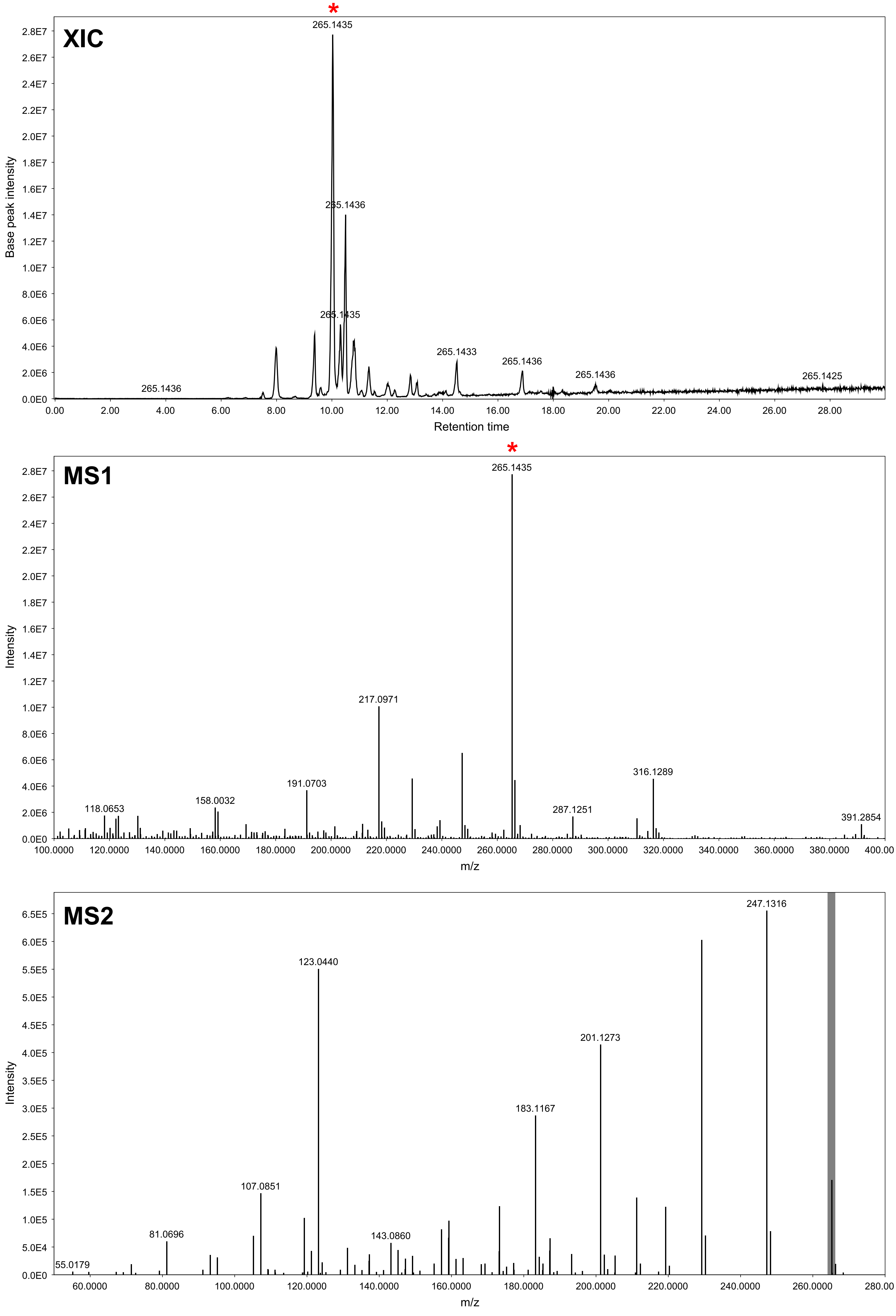

### GNPS hit 9d (MEB) - XIC range m/z 265.142-265.144 [M+H]

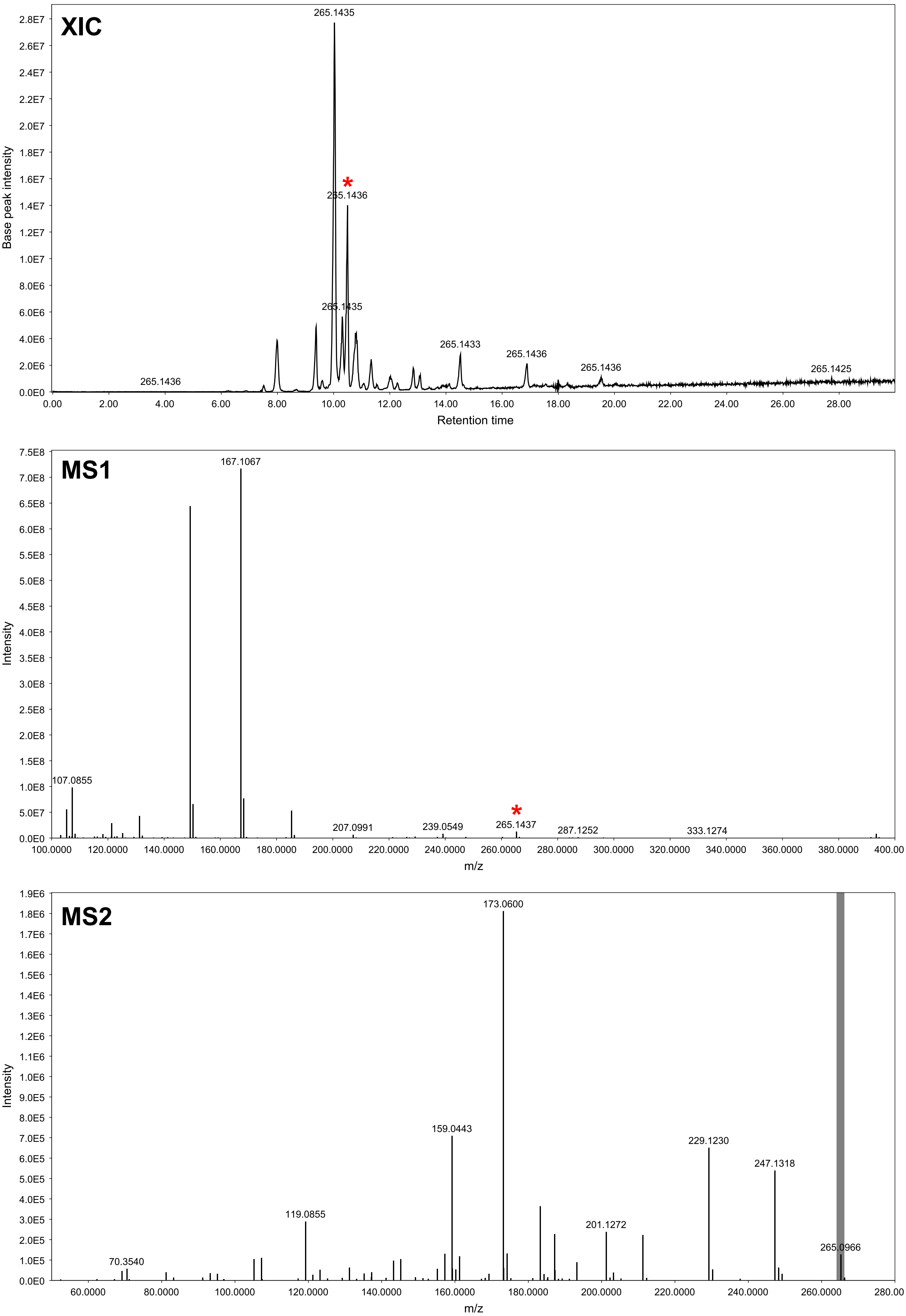

### GNPS hit 10 (PDA) - XIC range m/z 265.142-265.144 [M+H]

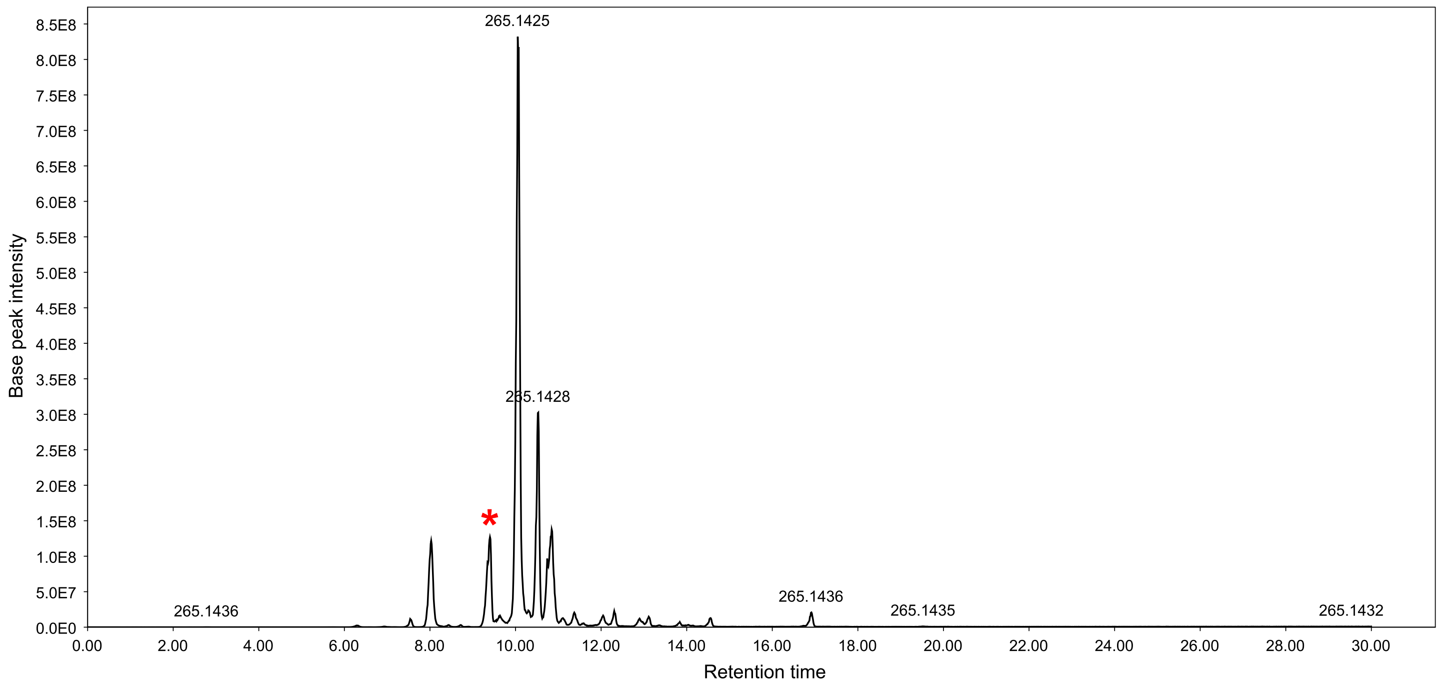


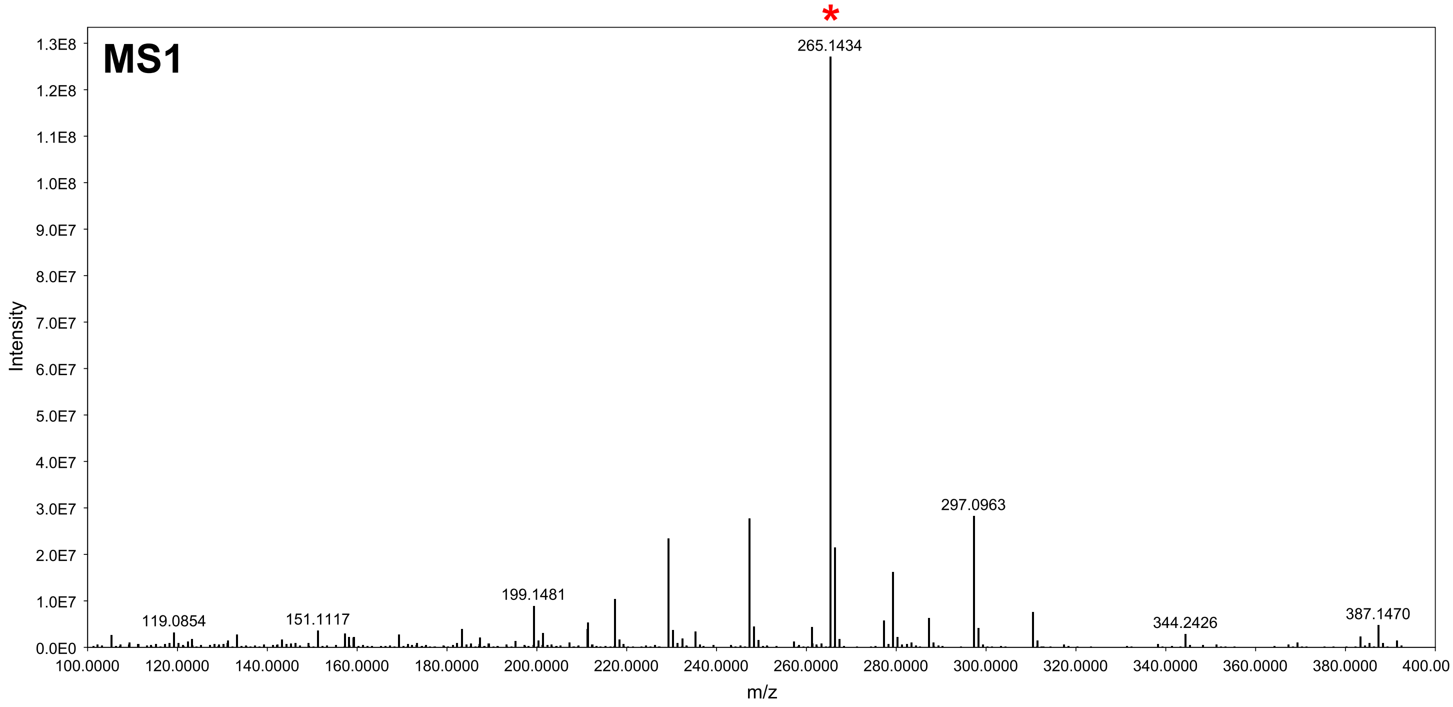


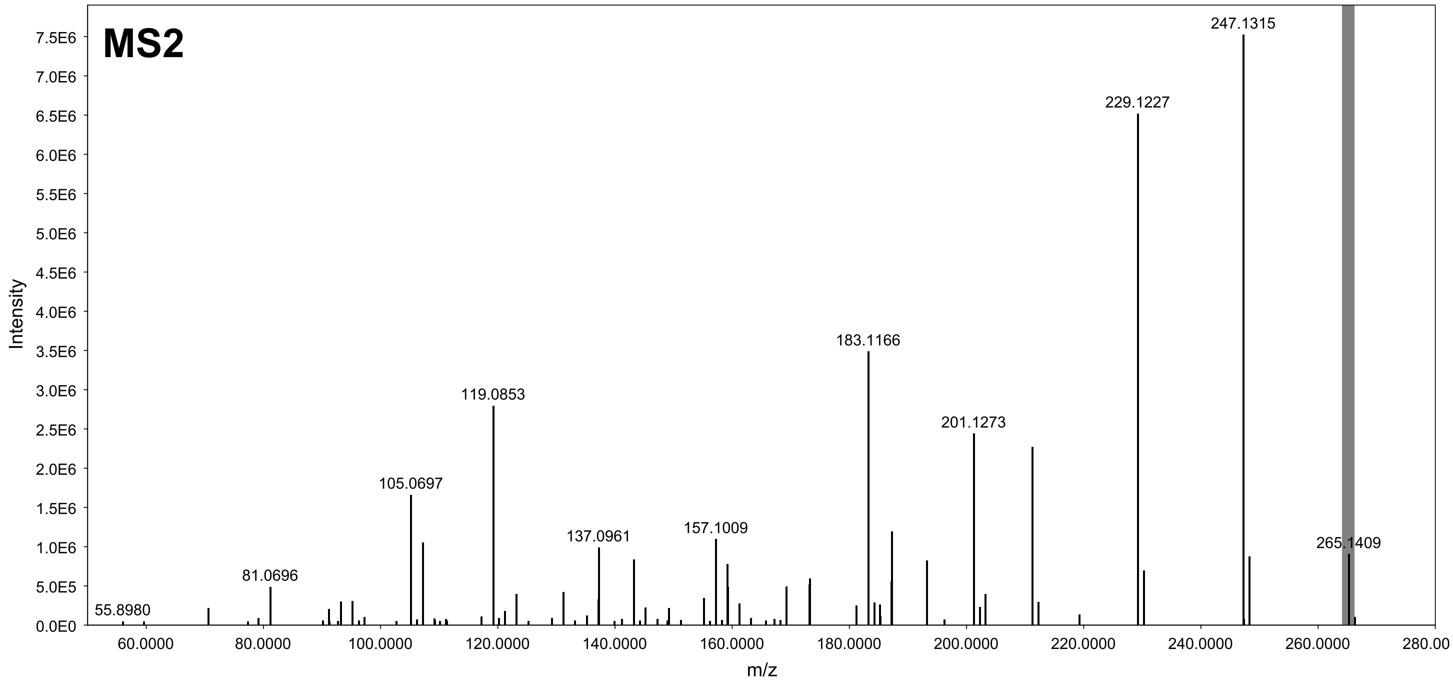

### GNPS hit 11a (MEB) - XIC range m/z 231.137-231.139 [M-H2O+H]

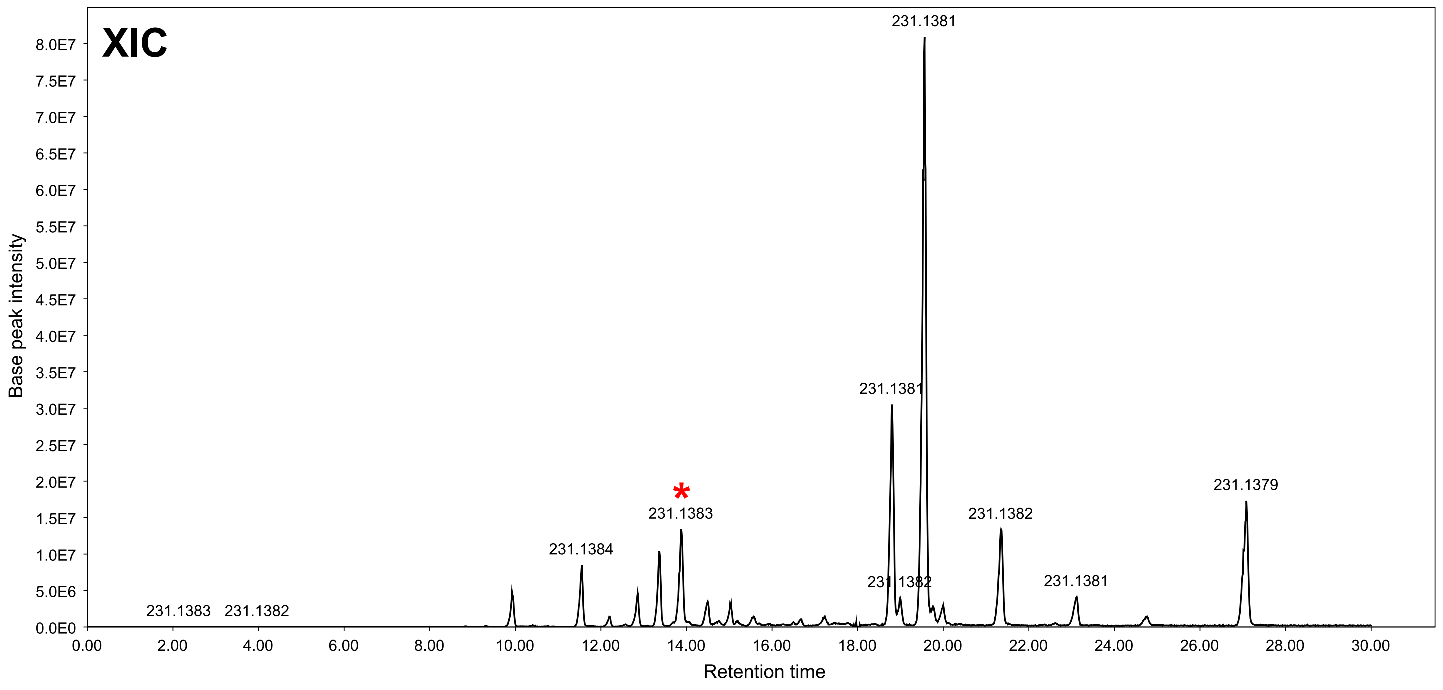


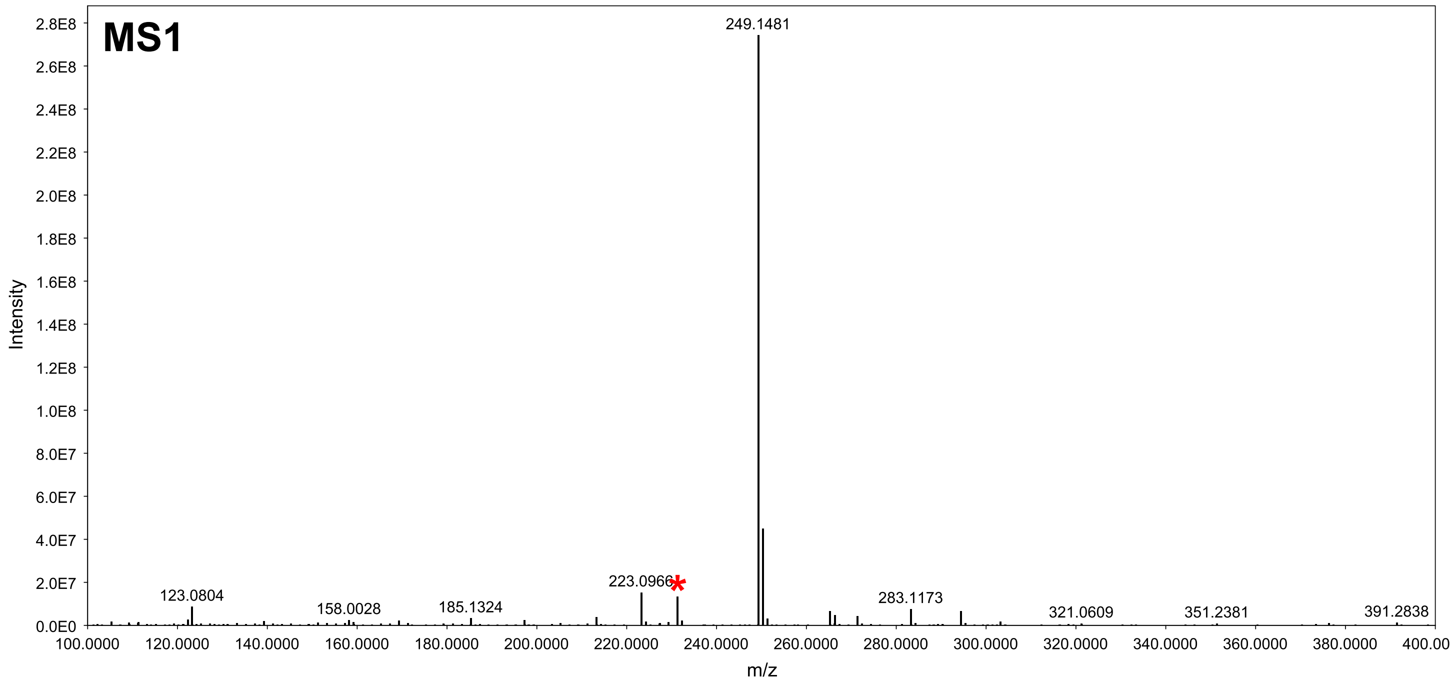


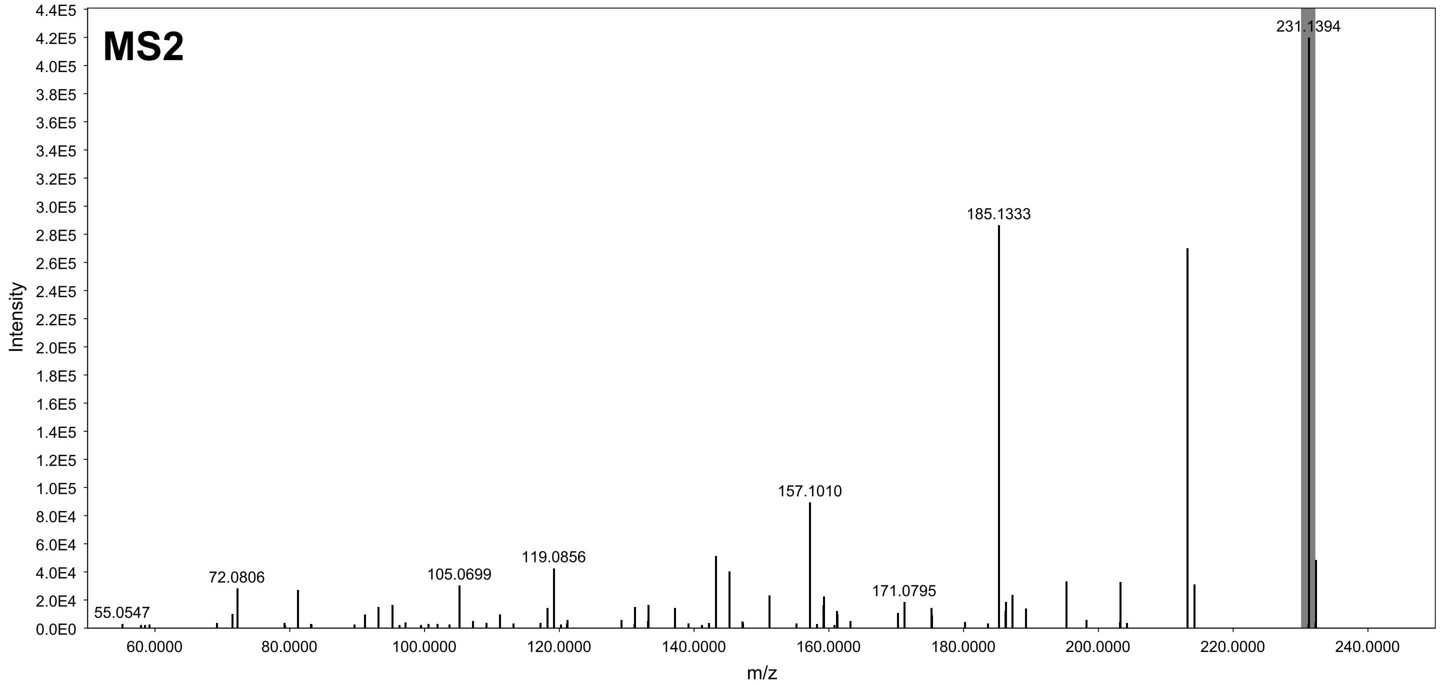

### GNPS hit 11b (MEB) - XIC range m/z 231.137-231.139 [M-H2O+H]

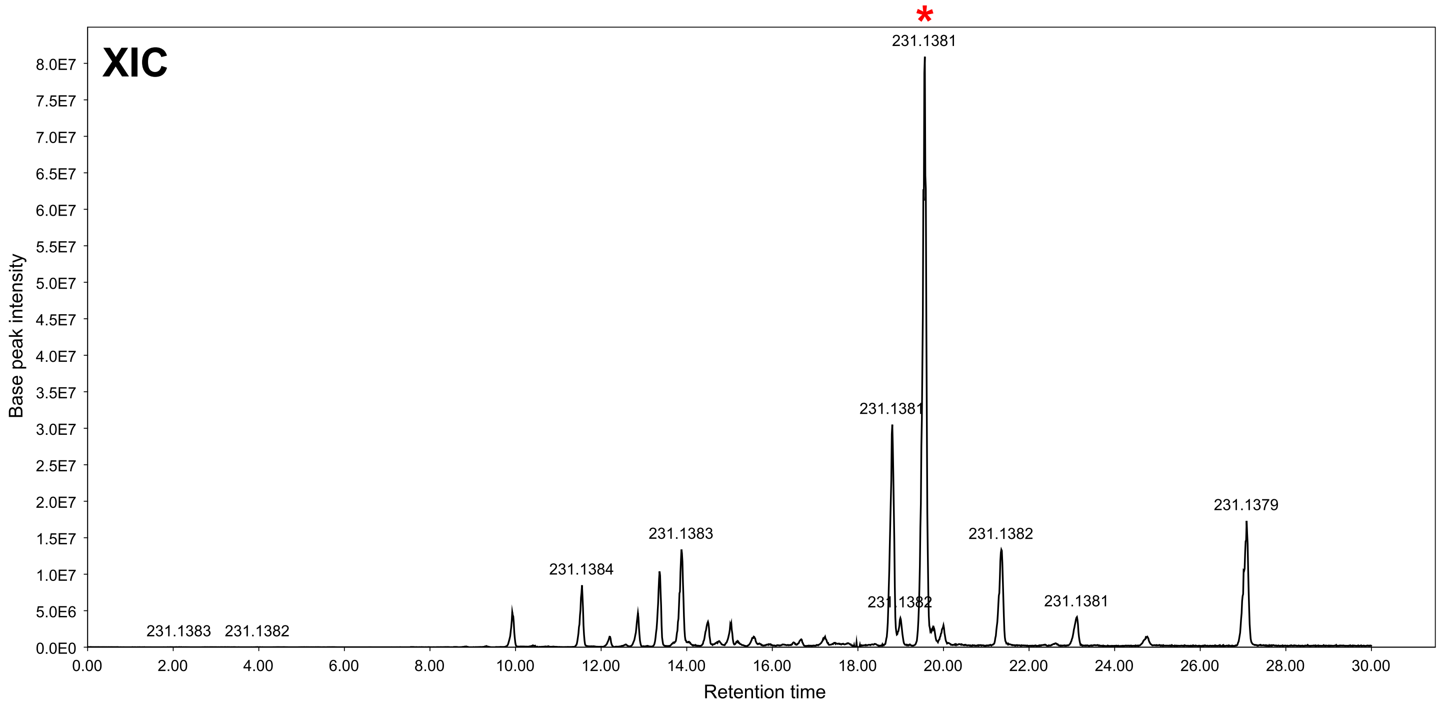


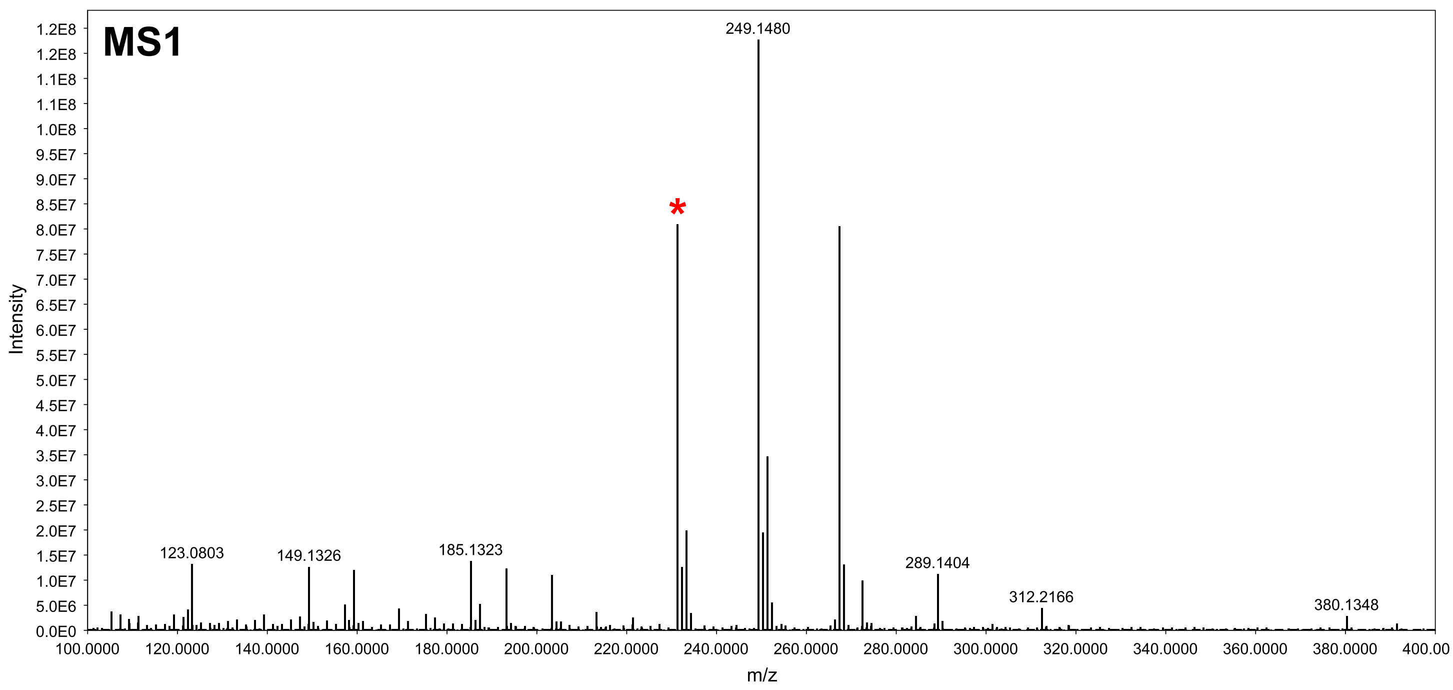


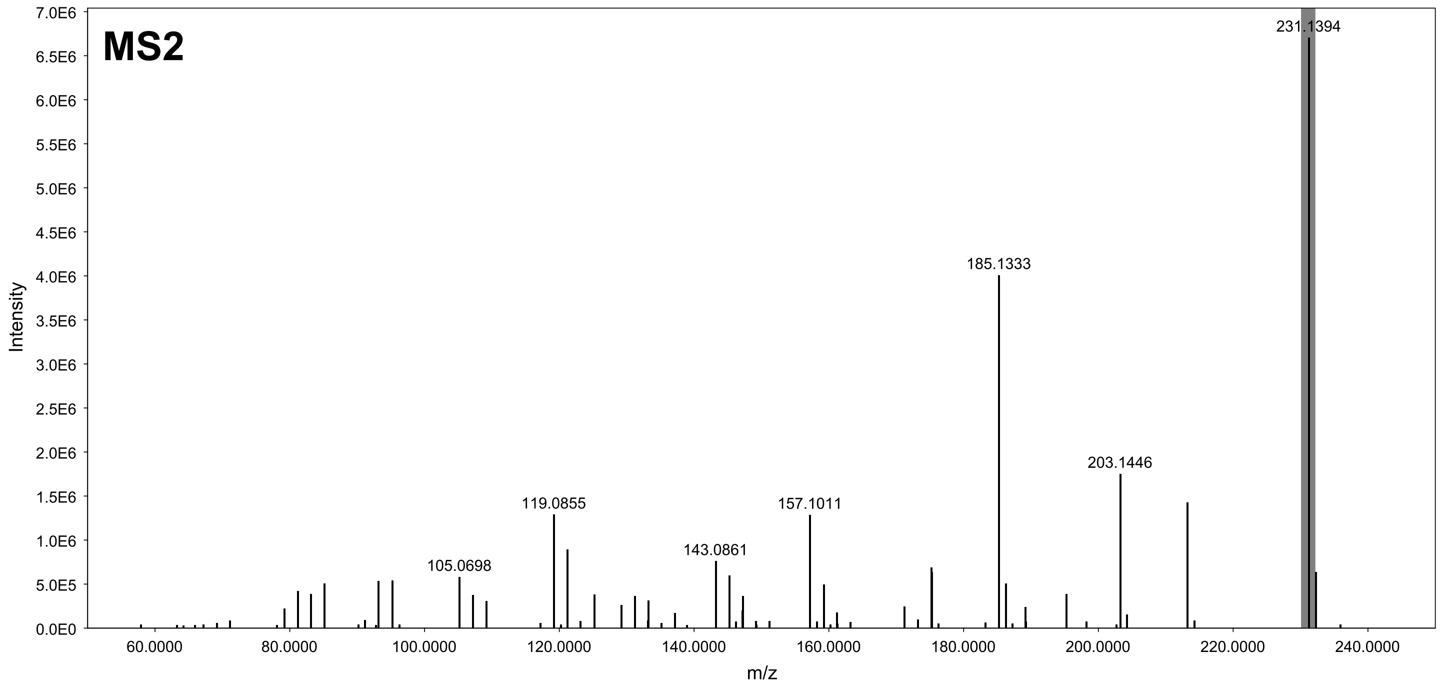

### GNPS hit 11c (PDA) - XIC range m/z 231.137-231.139 [M-H2O+H]

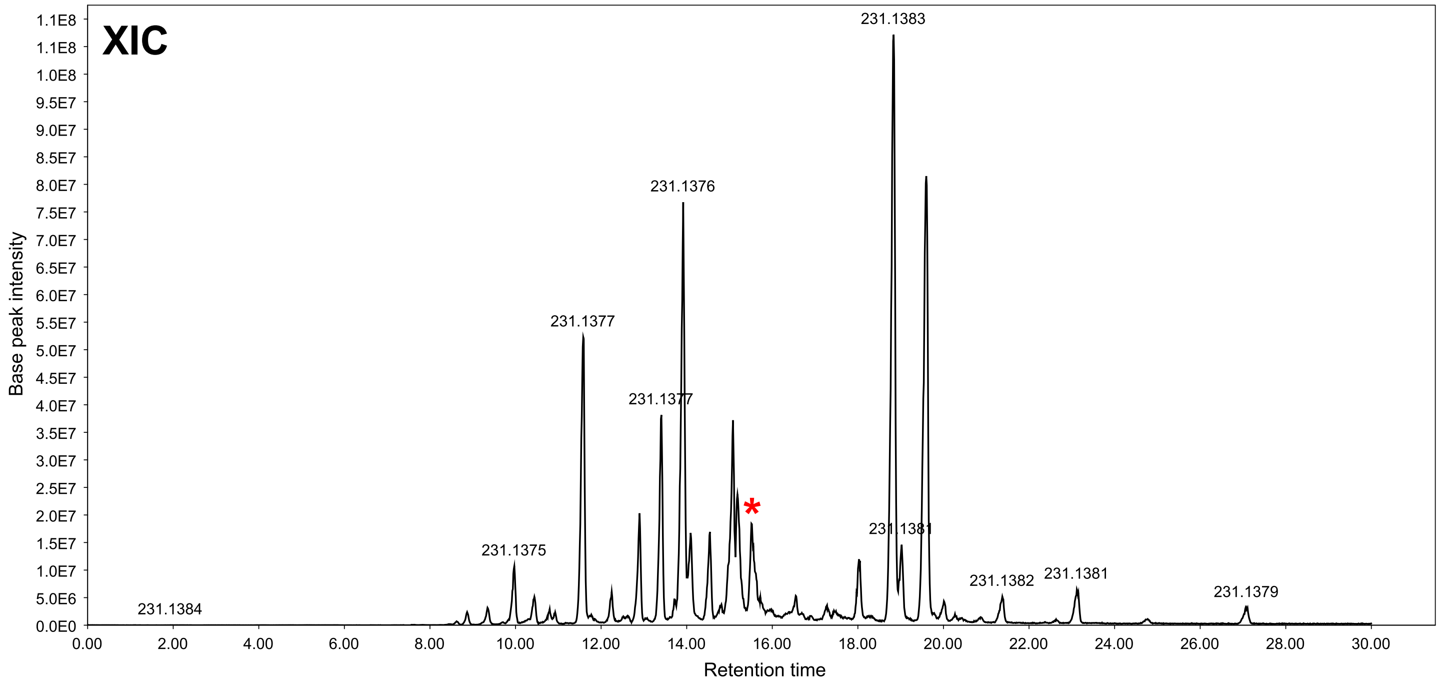


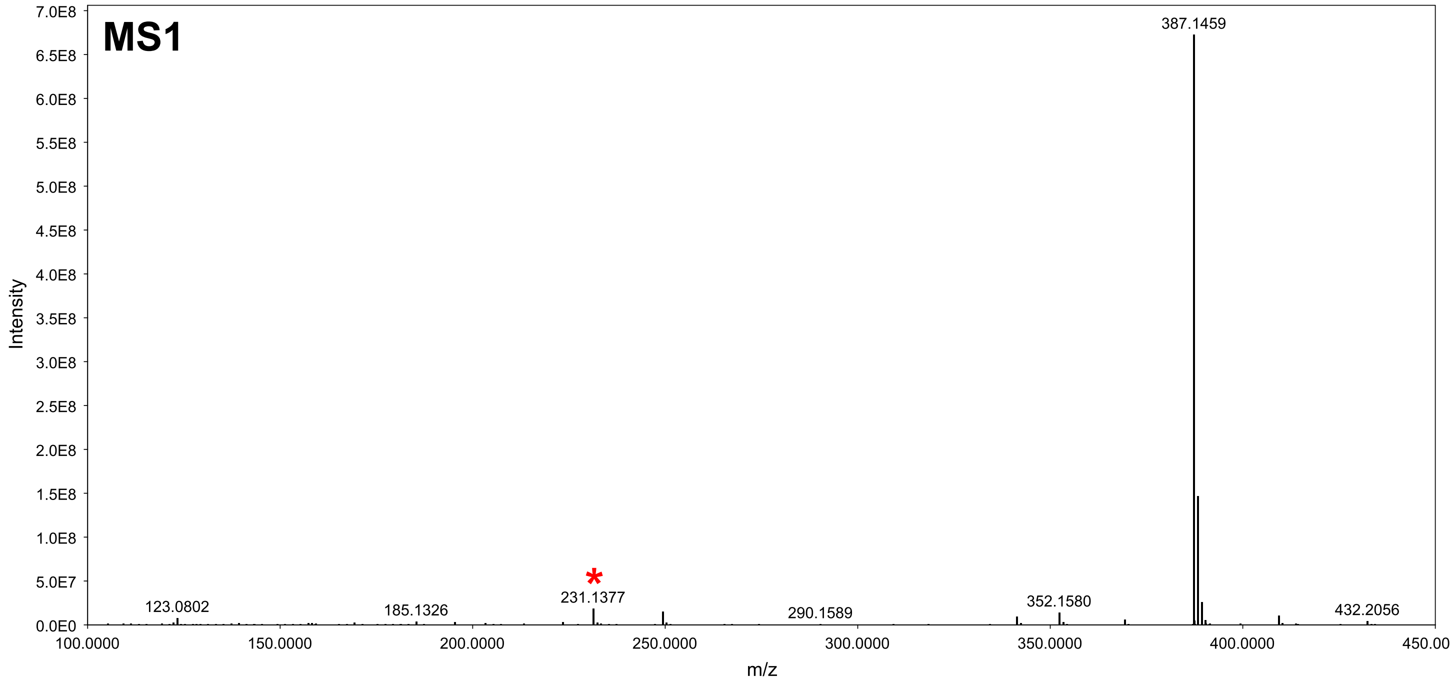


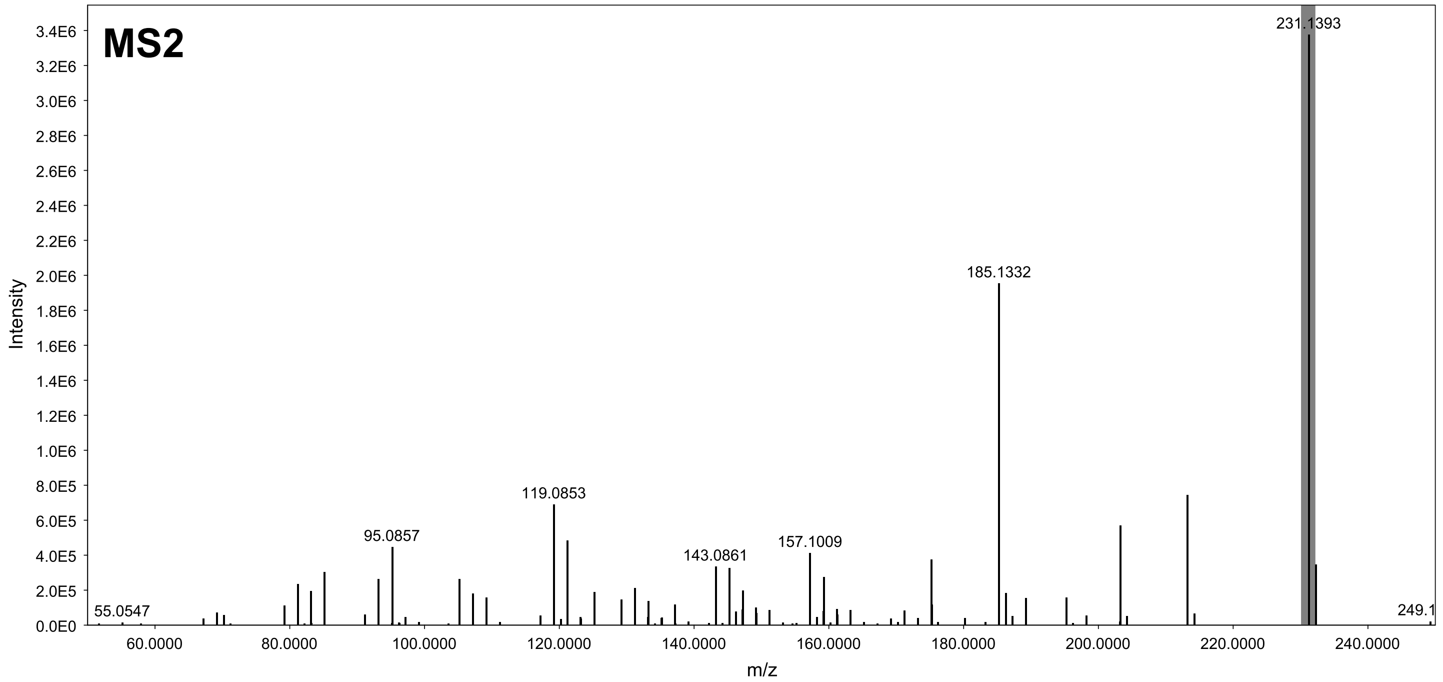

### GNPS hit 11d (PDA) - XIC range m/z 231.137-231.139 [M-H2O+H]

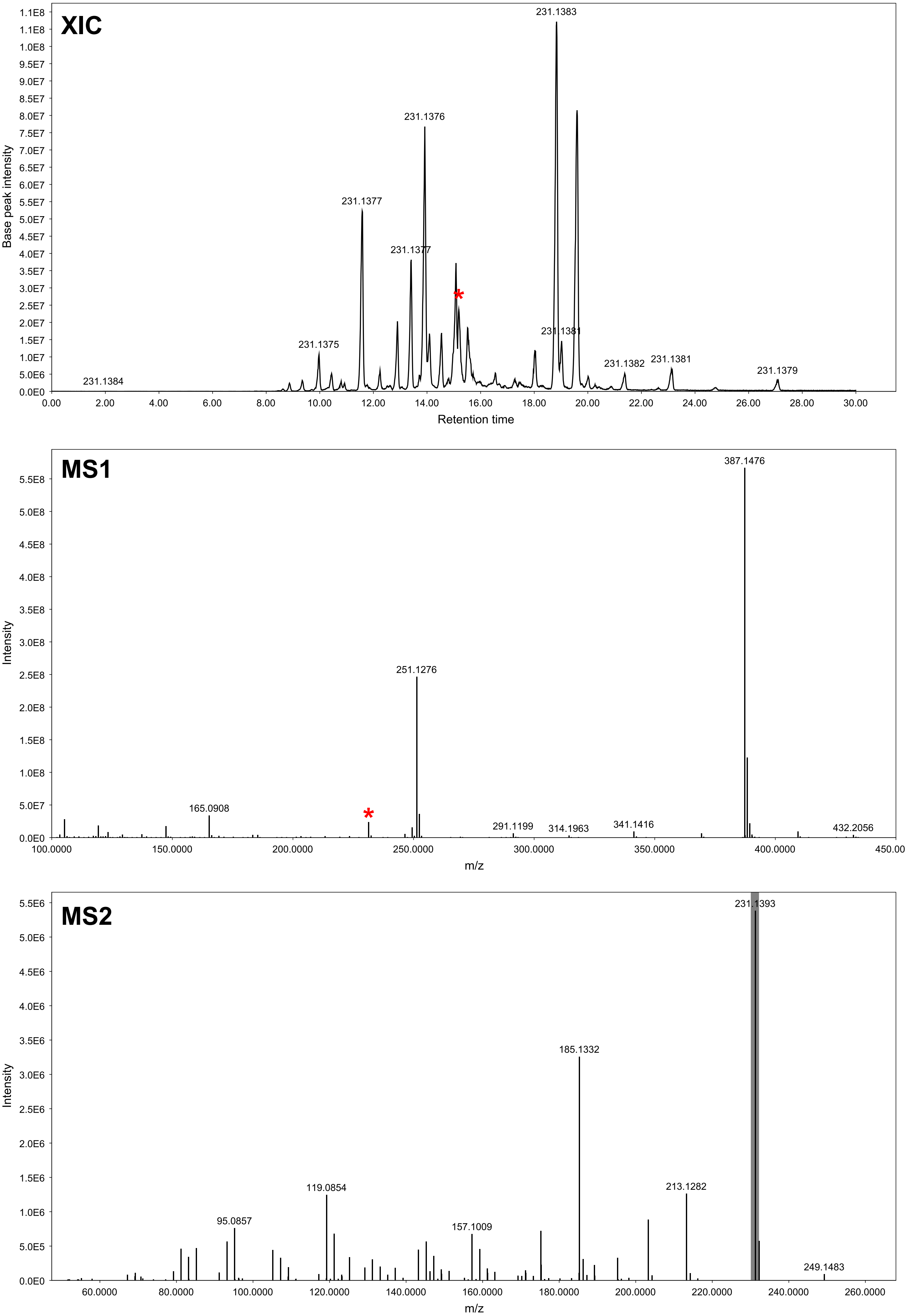

### GNPS hit 12a (PDA) - XIC range m/z 231.137-231.139 [M+H]

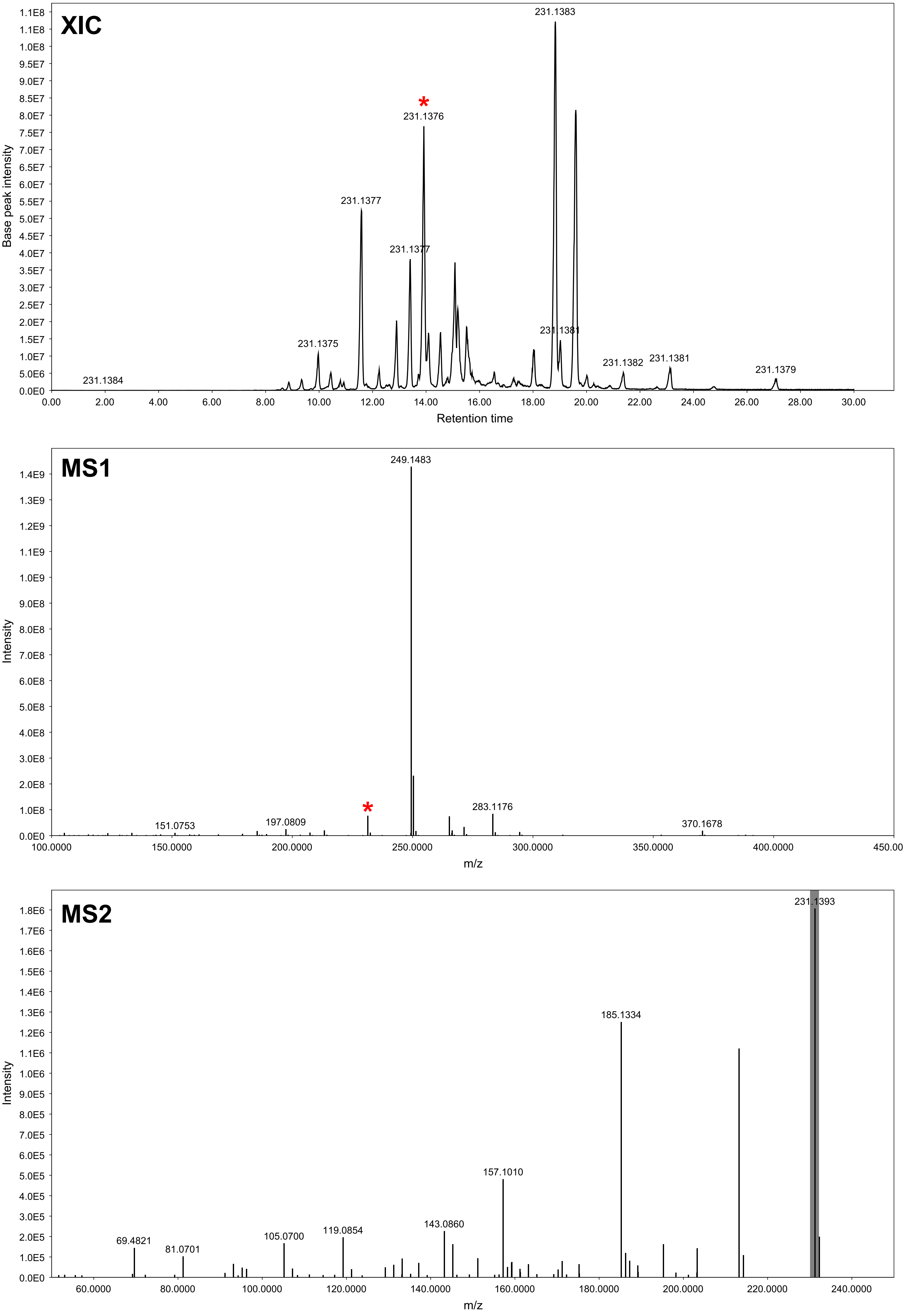

### GNPS hit 12b (PDA) - XIC range m/z 231.137-231.139 [M+H]

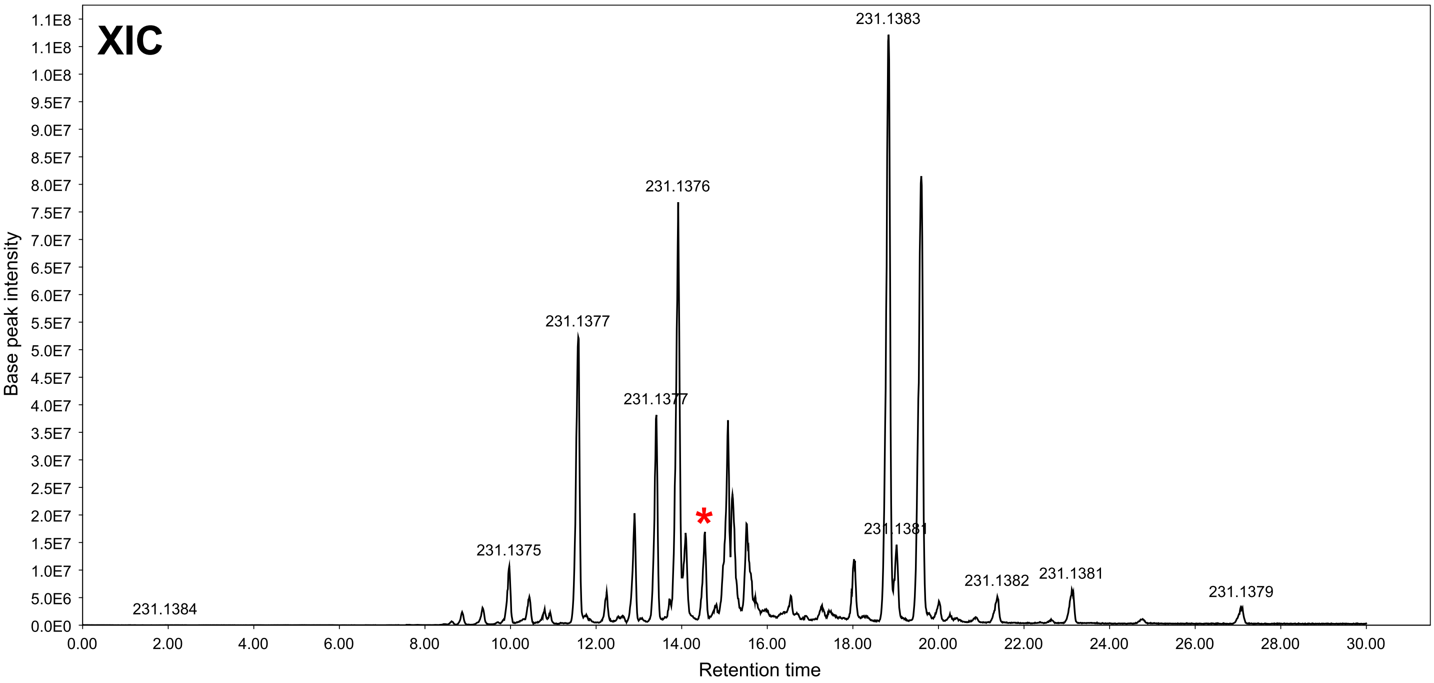


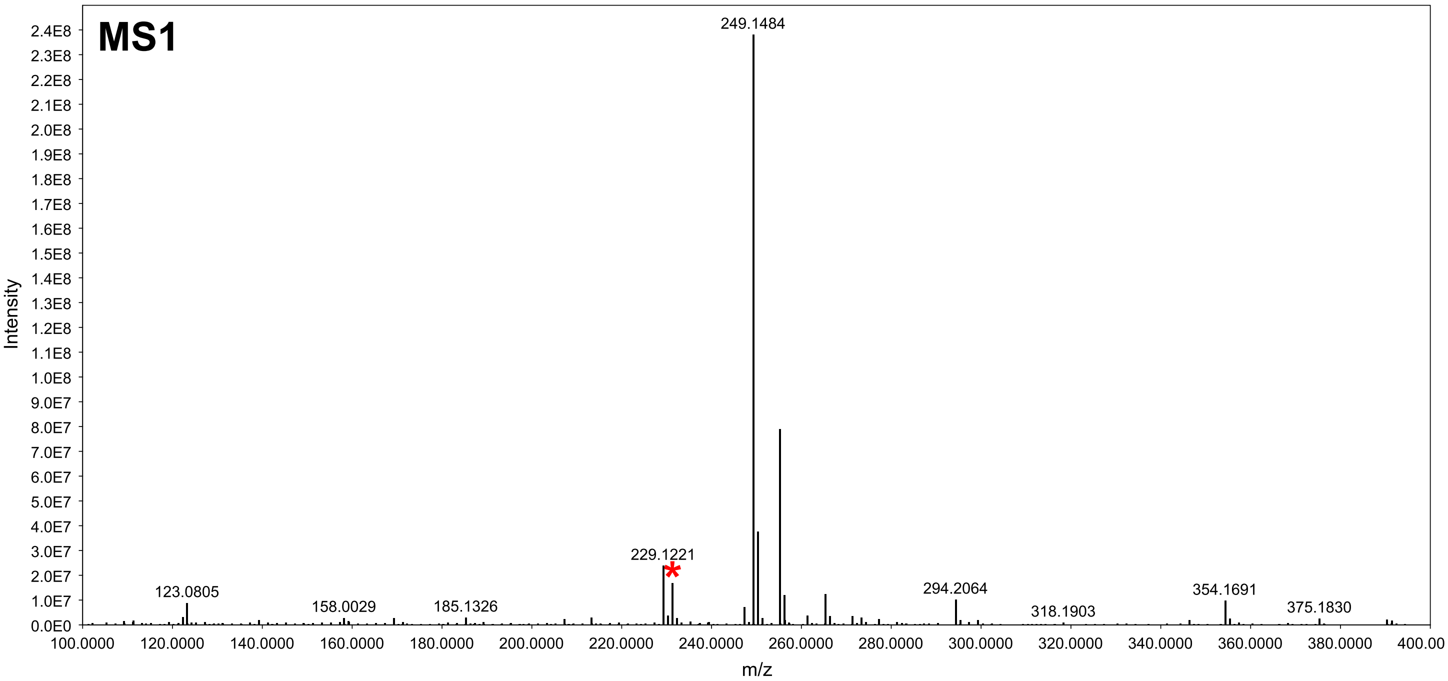


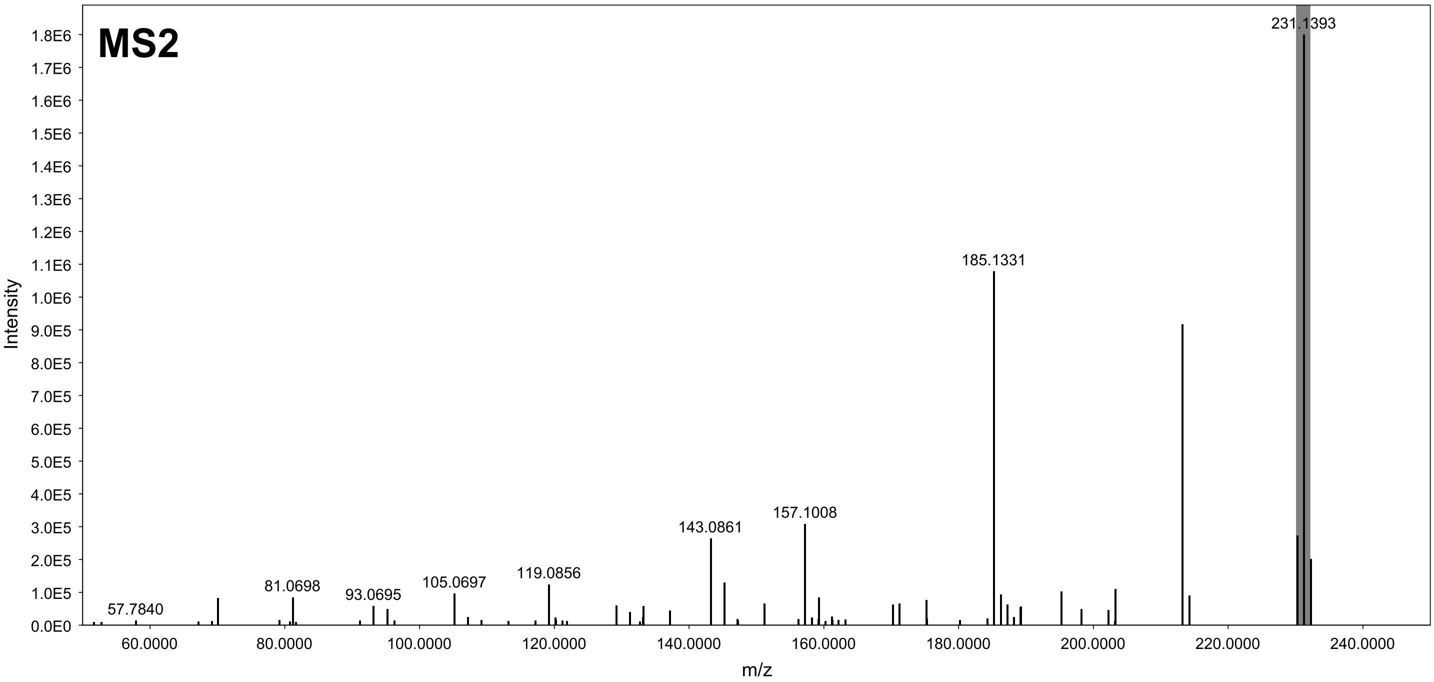

### GNPS hit 12c (MEB) - XIC range m/z 231.137-231.139 [M+H]

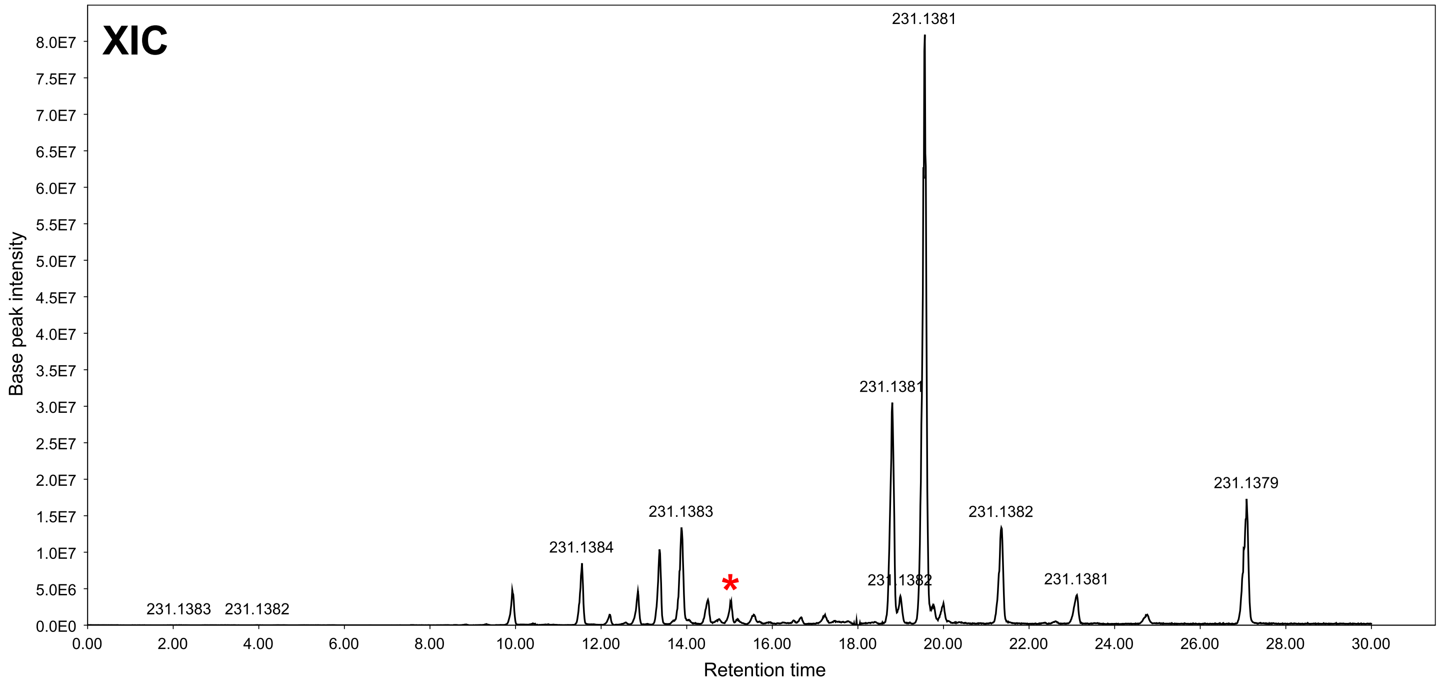


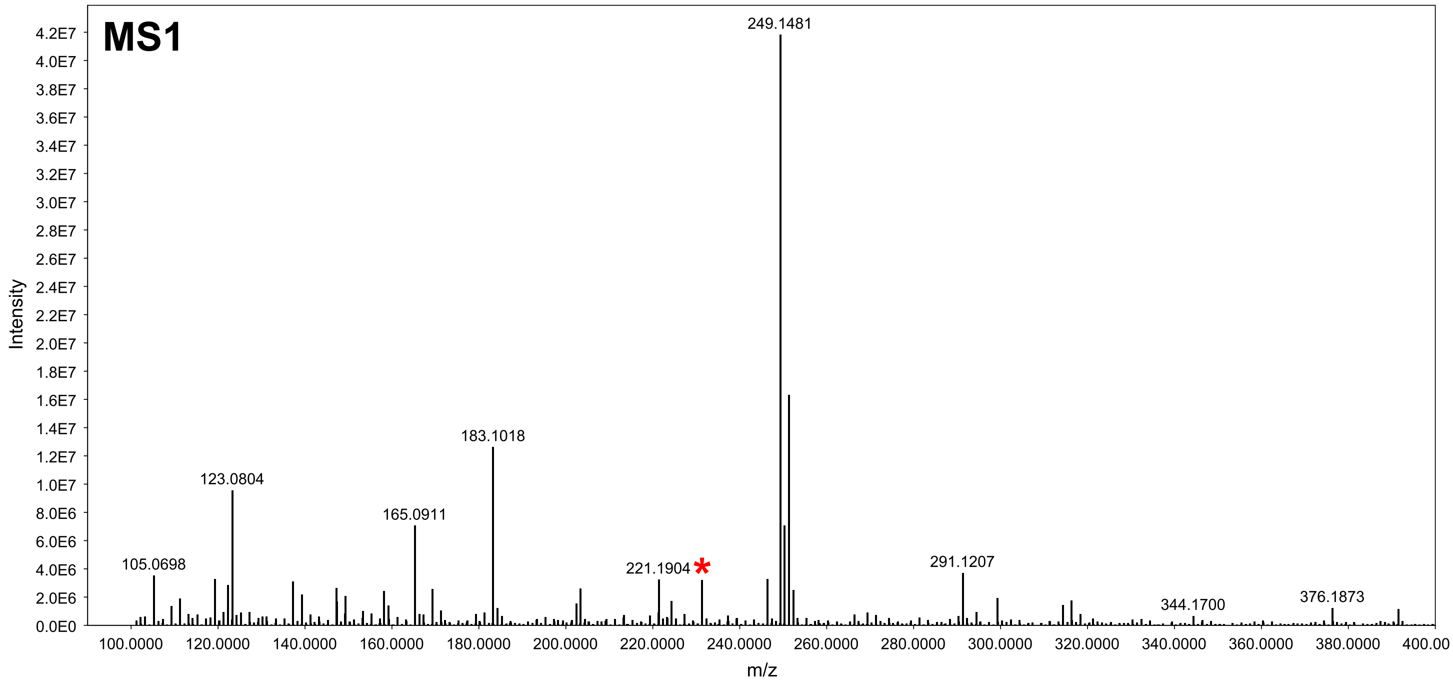


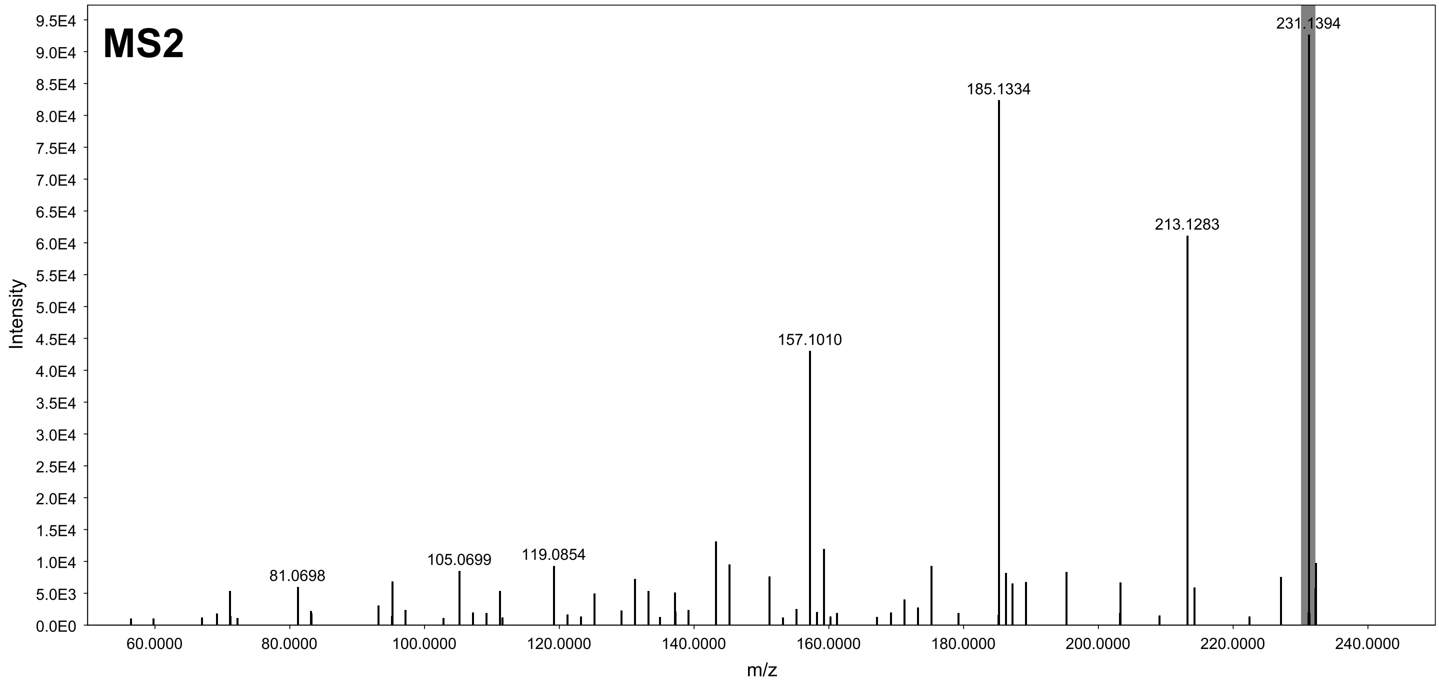

### GNPS hit 13 (MEB) - XIC range m/z 607.107-607.109 [M+H]

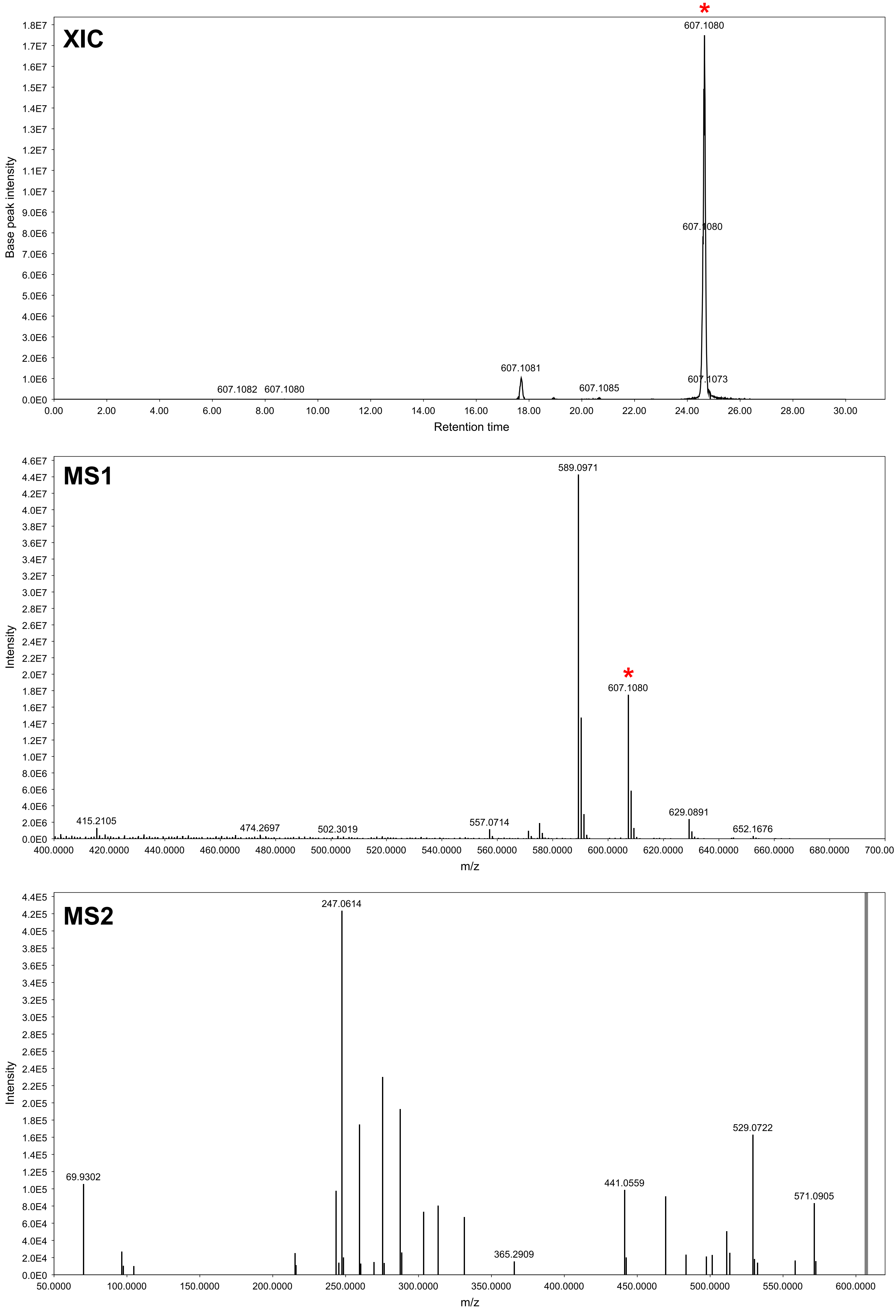

### GNPS hit 14 (MEB) - XIC range m/z 267.158-267.160 [M+H]

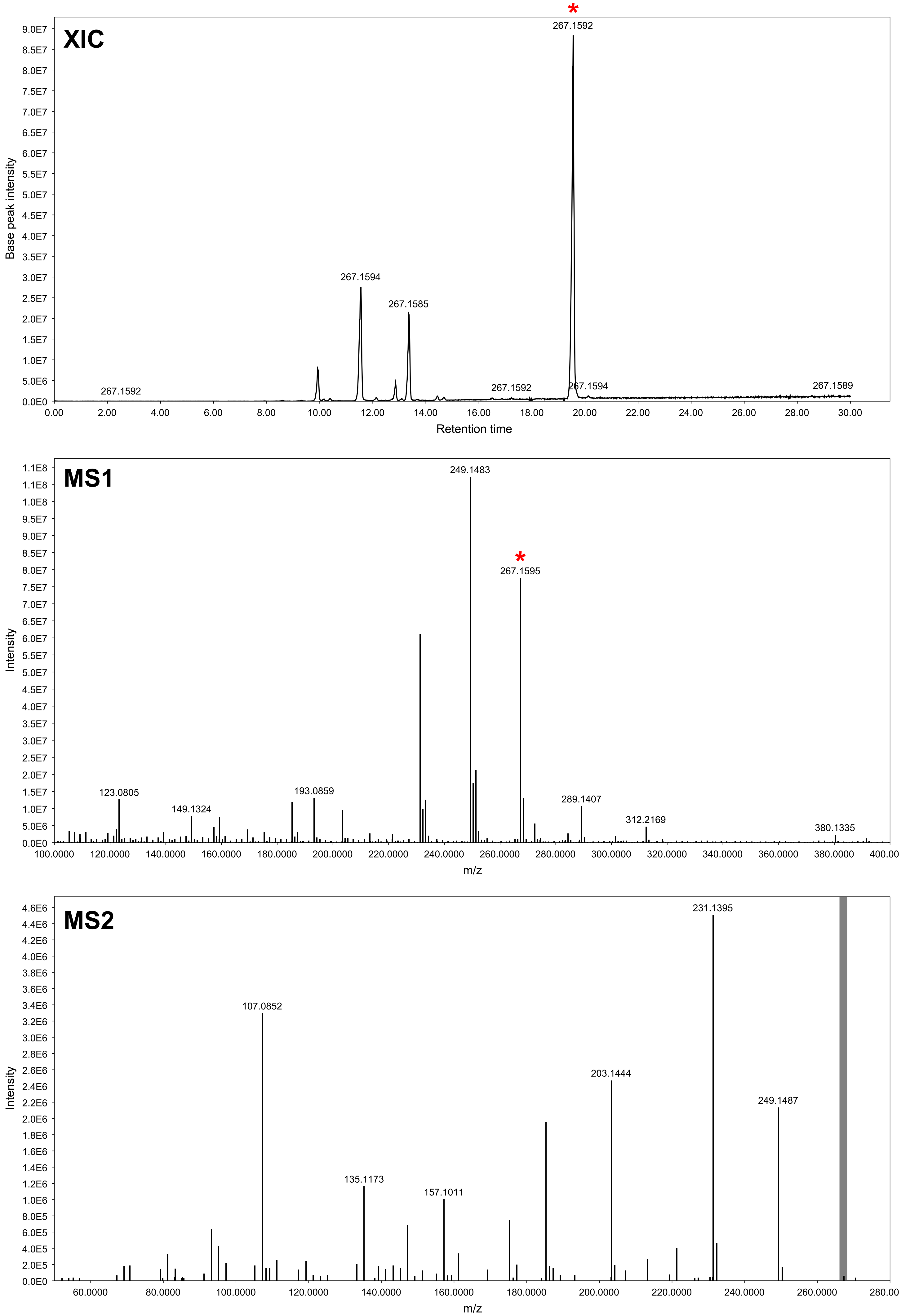

### GNPS hit 15 (MEB) - XIC range m/z 233.152-233.154 [M+H]

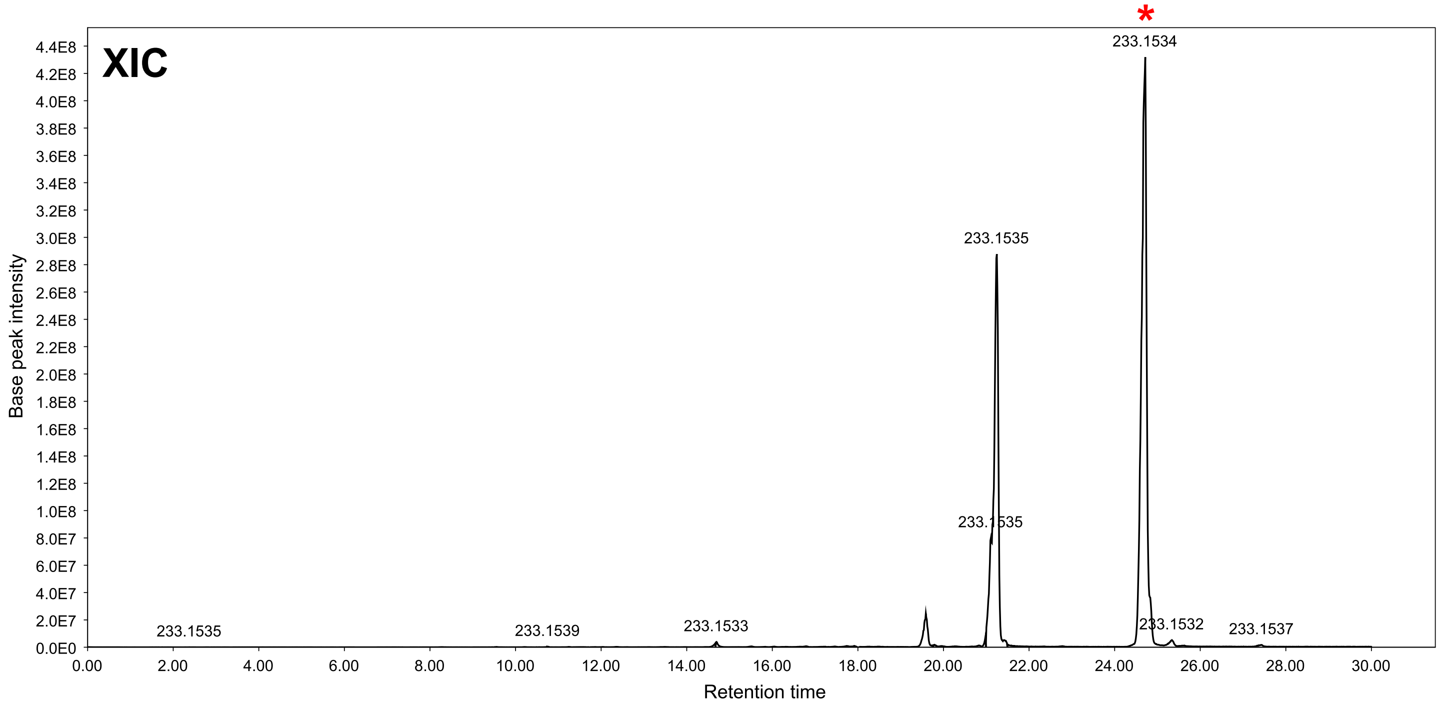


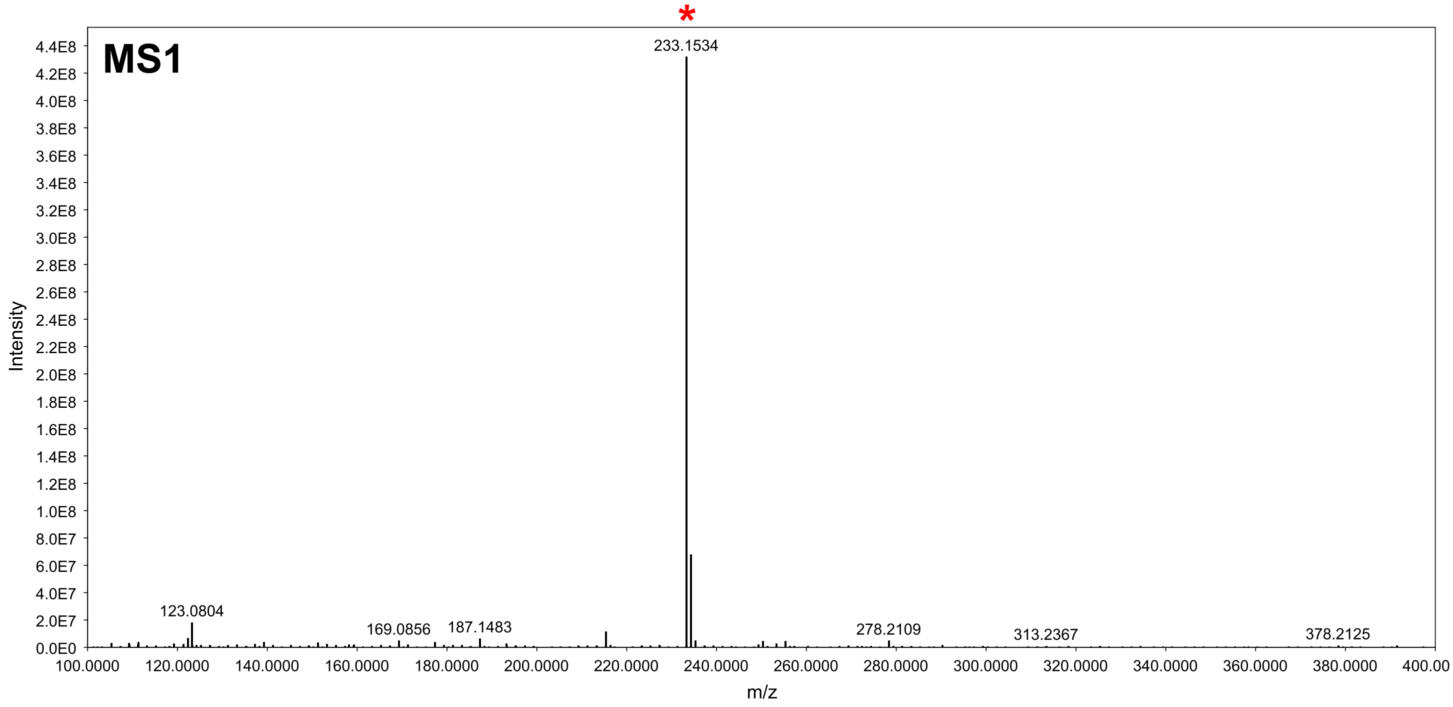


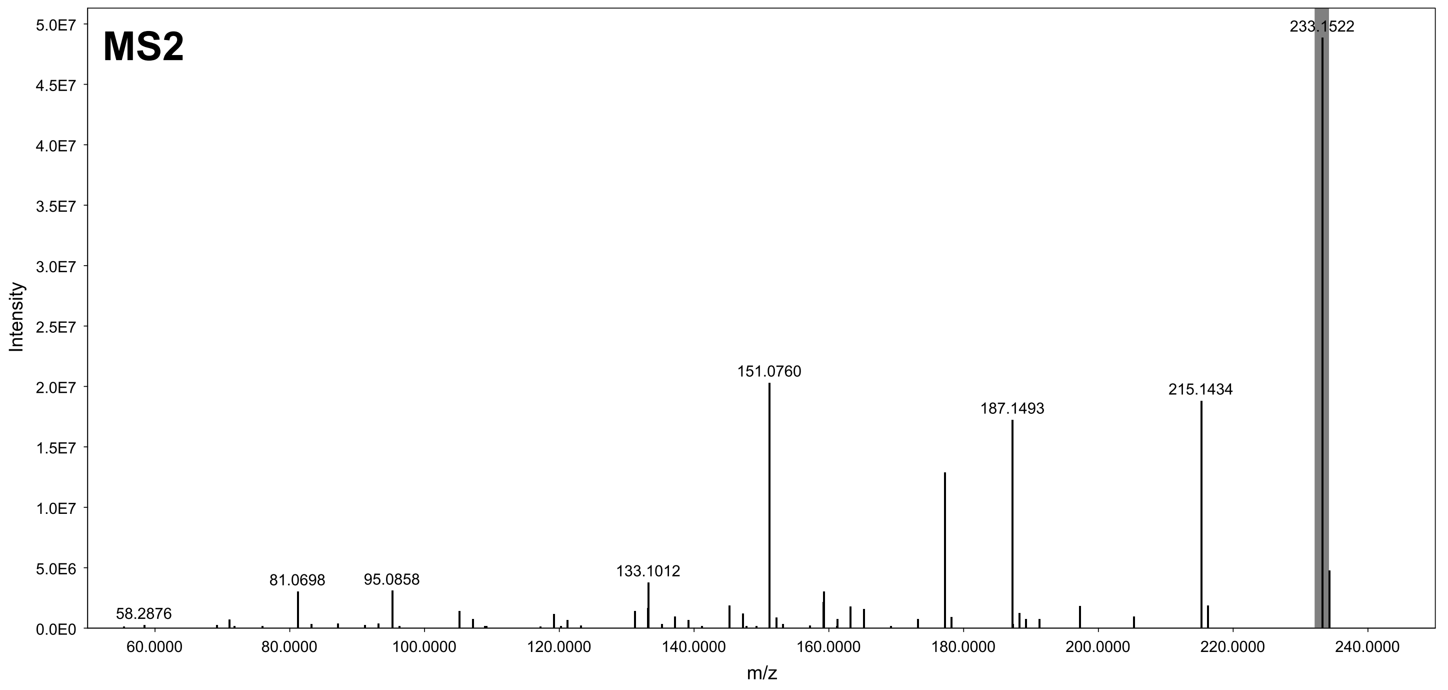
